# *Castling*, a novel therapeutic concept for rewiring pathological gene-expression networks, enabled by the TRIPLE technology

**DOI:** 10.64898/2026.01.28.702195

**Authors:** Dinu Antony, Maria Silvia Roman Azcona, Hagar Kalinski, Marianne Pultar, Swetlana Adamsky, Sharon Avkin Nachum, Eyal Shalom, Manuel Rhiel, Andreas Tsouris, Andreas Diendorfer, Geoffroy Andrieux, Melanie Boerries, Matthias Hackl, Tatjana I. Cornu, Daniel Zurr, Toni Cathomen, Elena Feinstein, Claudio Mussolino

## Abstract

Pathological conditions often arise from dysregulation of complex gene networks, with microRNAs (miRNAs) acting as central modulators. Disease progression is frequently characterized by upregulation of “disease-promoting” miRNA, suppressing beneficial pathways, and concomitant downregulation of “protective/therapeutic” miRNAs, normally restraining pathological programs. Since individual miRNAs coordinately regulate multiple genes, their manipulation represents powerful therapeutic intervention, yet synthetic or ectopically overexpressed miRNA mimics or inhibitors may perturb physiological miRNA processing and/or cause off-target effects. We hypothesized that pathological gene regulatory imbalances could instead be corrected by rewiring endogenous miRNA regulation. Specifically, by placing downregulated “protective/therapeutic” miRNAs under the control of promoters activated in pathology, and driving overexpression of “disease-promoting” miRNAs, thereby disabling the pathogenic program while inducing the therapeutic one in a single editing event. We termed this concept *castling*, after the chess move. For effective implementation of *castling*, we developed TRIPLE (Targeted Replacement Induced by Persistent Locus Editing), a novel genome-editing procedure enhancing homology-directed repair through sequential cleavage. As proof of concept, we *castled* miRNAs inversely regulated during onset of CAR T cell dysfunction in a model of chronic antigen stimulation. *Castled* CAR T cells exhibited a delayed dysfunction enabled by up- and downregulation of relevant gene subsets.

## Introduction

The dysregulation of complex gene expression networks drives the development of many diseases and pathological conditions, where coordinated changes in expression of multiple genes underlie dysfunctional cellular states. In the early stages of disease, transient changes in transcription often represent adaptive responses aimed at restoring homeostasis^1^. However, as pathological stimuli persist, these regulatory programs become increasingly distorted, leading to the sustained activation of genes that promote disease progression and the repression of genes that maintain normal cellular function^2^. MicroRNAs (miRNAs) have emerged as central regulators in these dynamic transitions, acting as critical post-transcriptional switches that fine-tune gene network activities with disease progression being frequently associated with imbalance in miRNA expression. In this context, miRNAs that suppress pathways regulating cellular integrity are progressively overexpressed, while those that inhibit disease-supportive pathways are progressively silenced. For instance, miR-21-5p and miR-155-5p are consistently upregulated in cancer and chronic inflammatory states^3–6^, where they reinforce proliferation and pro-inflammatory signaling, whereas miR-29a-3p, a key antifibrotic regulator, is repressed in tissue fibrosis, or miR-126-5p, essential for vascular homeostasis, is downregulated in cardiovascular disease^7,8^. This coordinated dysregulation contributes to the establishment of pathological gene expression programs that sustain disease and suppress recovery. Since miRNAs naturally control the expression of multiple genes simultaneously, they have emerged as attractive therapeutic targets for modulating complex gene circuits and the broader signaling pathways they govern.

Existing approaches, such as miRNA mimetics to restore a protective miRNA exhibiting low expression or antimirs to inhibit overexpressed pathogenic miRNAs, have shown conceptual promise^9–11^ but still suffer from several technical problems including poor physiological control^12,13^. For example, synthetic miRNA mimetics or antimirs may act for durations that may not match the therapeutical need whereas, overexpression of miRNA mimics or antimirs using, for example, gene therapy vectors may overwhelm the endogenous miRNA processing machinery, leading to off-target effects and even cytotoxic outcomes^14,15^.

To overcome these limitations, we hypothesized that therapeutic restoration of normal cellular function in the context of pathology could be achieved by strategically rewiring miRNA regulation within the genome itself. Specifically, we reasoned that placing the sequence of an insufficiently expressed therapeutically beneficial miRNA under the control of a promoter that is activated under pathological conditions, would achieve two complementary goals: (i) high expression of the therapeutic miRNA during persistence of the pathological conditions, and (ii) disruption of the disease-promoting miRNA expression at that locus. This dual-action strategy would engage corrective regulatory mechanisms precisely during the pathological state, effectively using the disease’s own transcriptional machinery against itself (**Figure 1a**). We termed this concept *castling*, after the strategic chess move that simultaneously repositions and protects key chess pieces on the board. Implementing *castling* required an advanced genome-editing strategy capable of performing precise sequence replacement at high efficiency. To this end, we developed TRIPLE (Targeted Replacement Induced by Persistent Locus Editing), a novel homology-directed repair–based approach that through recurrent cleavage, prolongs the repair time window at double-stranded breaks, markedly enhancing the frequency of precise insertion editing and enabling the reliable execution of *castling*.

**Figure 1.**
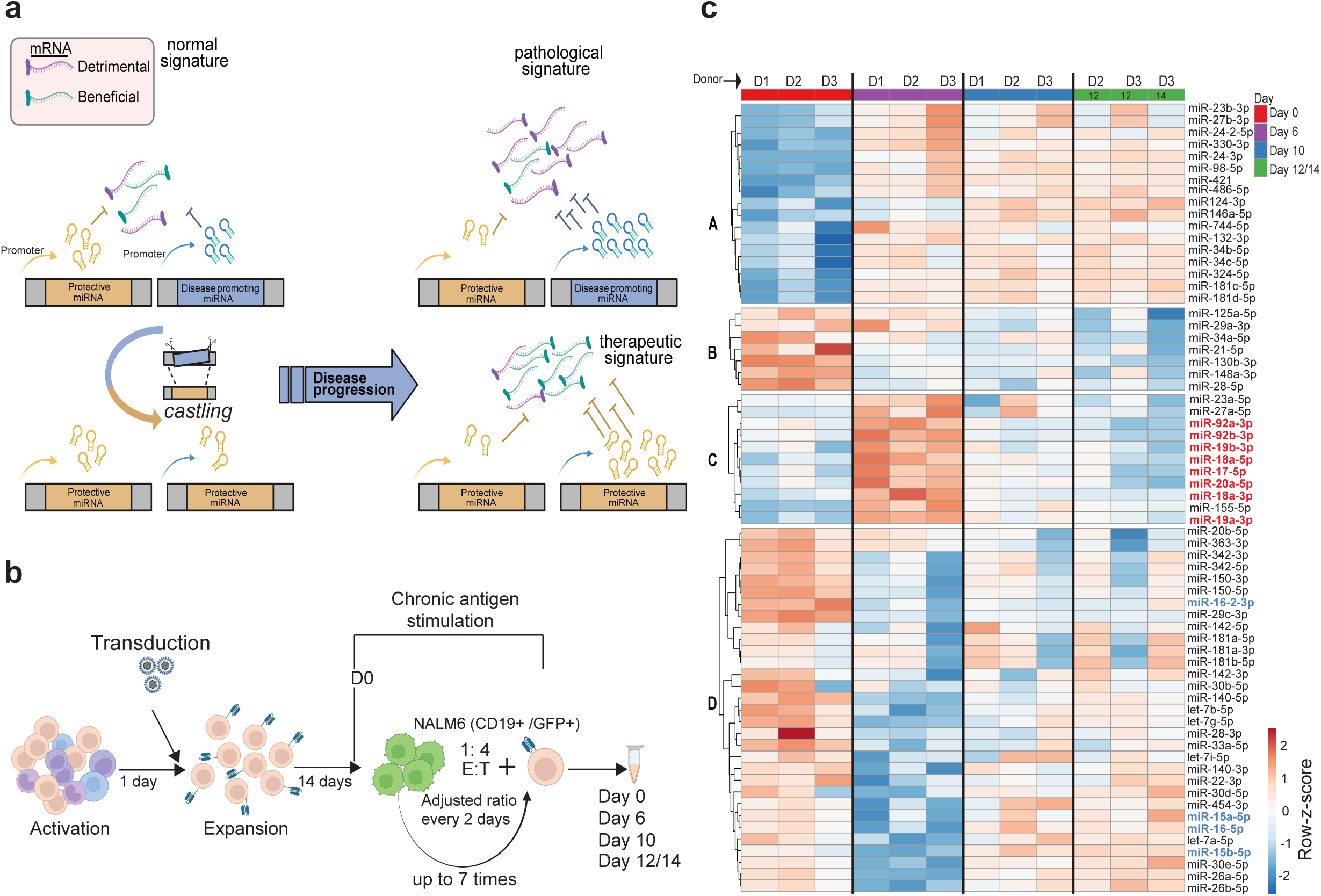
Identification of candidate miRNA pairs for *castling*. **a.** Schematic presentation of the miRNA *castling* concept. **b.** Schematic presentation of the T cell manipulation flow. Upon thawing of peripheral blood mononuclear cells (PBMC), T cells are activated and 24-hours later, transduced with a lentiviral vector harboring the CD19-specific CAR construct. Following expansion (14 days), CD19+/GFP+ NALM6 tumor cells are added to the CAR T cells at a 1:4 effector-to-target (E:T) ratio (day 0) and chronic antigen stimulation with the tumor cells continues for additional 12-14 days, adjusting the E:T ratio every two days. The timepoints when the cells were harvested for subsequent miRNA/mRNA extraction and sequencing are indicated. **c.** miRNA sequencing results. Heatmap depicts ln-transformed reads per million (RPM) after row-based scaling values for 66 miRNAs, which were found to be differentially expressed (FDR < 0.05) on day 6 as compared to day 0 (pre-stimulation), and which demonstrated medium or high abundance (mean logCPM > 8). Letters A-D on the left side of the plot designate the types of miRNA expression profiles. Each group of three columns represents miRNA expression data obtained from cells originating from three independent T cell donors (D1, D2 and D3) and harvested at the indicated time points (day). Unit variance scaling was applied to rows, and clustered using Pearson correlation distance with average linkage. miRNAs selected for *castling* (miR-92b-3p and miR-17∼92 cluster from profile C; miR-15/16 clusters from profile D) are highlighted in red or blue, respectively.

Here we demonstrate that *castling* can be used to rewire a pathogenic miRNA regulatory network into a configuration that sustains cellular function. For the proof of *castling* concept, we used an *in vitro* model of CAR T cell dysfunction which is progressively induced by chronic antigen stimulation. This model reproduces, at least in part, a known T cell exhaustion phenotype, a dysfunctional state characterized by diminished effector function, sustained expression of inhibitory receptors, restricted proliferative capacity and altered transcriptional programs^16,17^. Notably, in the clinics, engineered CAR T cells also acquire an exhausted phenotype upon entering the tumor microenvironment where they are exposed to target tumor antigens and tumor-derived inhibitory factors, underscoring that CAR T cell exhaustion is a major barrier to their durable therapeutic efficacy^18,19^. In line with this, several therapeutic strategies aim to mitigate exhaustion, for example, by blocking checkpoint receptor–ligand interactions through antibodies or gene-editing approaches, thereby delaying the dysfunction onset^20,21^. We first conducted a time-resolved analysis of miRNA expression in this model and identified inversely regulated miRNA pairs whose temporal expression patterns suggested opposing roles in promoting versus counteracting CAR T cells dysfunction. Guided by these findings, we employed TRIPLE to efficiently replace disease-promoting miRNAs, at their endogenous loci, with sequences of protective miRNA counterparts. This intervention reactivated gene expression programs associated with sustained effector function and prolonged cytotoxic activity while disabling the inhibitory programs, collectively translating into a measurable delay in the onset of CAR T cell dysfunction.

Together, our findings establish *castling* as a versatile approach for disease-responsive genome reprogramming and TRIPLE as its enabling technology. This work lays the foundation for the development of a next-generation therapeutic platform in which disease-associated regulatory networks can be precisely rewired at the genomic level to sustain cellular health and combat complex pathologies.

## Results

### Identification of miRNA pairs for *castling*

To demonstrate that *castling* can rewire pathological gene networks, we first sought to identify suitable miRNA candidates for swapping. We modified an existing chronic antigen stimulation assay that induces progressive CAR T cell dysfunction, mimicking continuous antigen exposure (**Figure 1b**)^16,17,22^. To this end, CD19-directed CAR T cells were co-cultured with CD19⁺/GFP^+^ NALM6 leukemia cells for up to 14 days, with tumor cells replenished every two days. Despite donor-to-donor variability, monitoring CAR T cell expansion revealed a reproducible kinetic profile in which proliferation peaked between days 6 and 10, followed by proliferative decline and complete proliferative arrest by day 12-14 (**Supplementary Fig. 1a**). At that point, chronically stimulated CAR T cells displayed additional hallmark features of dysfunction, including declined cytotoxicity and cytokine (IFN-γ and IL-2) production compared to control CAR T cells that were cultured in parallel without antigen exposure (**Supplementary Fig. 1b-d**)^16^. Flow cytometric analysis confirmed the acquisition of a dysfunctional phenotype, marked by increased CD39 expression and concomitant loss of the T cell memory marker IL7R in the course of the CAR T and NALM6 cell co-culture (**Supplementary Fig. 1e-f**)^16^.

We next leveraged this system to capture the temporal dynamics of miRNA expression across the progressive loss of CAR T cell functionality by miRNA sequencing. The analysis was performed at four key time points: day 0 (pre-stimulation), day 6 (a peak of CAR T cell activity), day 10 (a transition phase between function and dysfunction), and day 12/14 (a fully dysfunctional state; **Figure 1b**). The observed dynamics in miRNAs expression were visualized as a heatmap revealing four principal expression profiles (**Figure 1c**): miRNAs progressively upregulated (A) or downregulated (B) in response to antigen stimulation, and miRNAs either peaking at day 6 after initiation of antigen stimulation with expression returning to day 0 baseline by days 10-12 (C) or reaching their lowest expression at day 6 of antigen stimulation with rebound to day 0 baseline by day 10 (D). Since day 6 coincided with optimal CAR T cell proliferative capacity (**Supplementary Fig. 1a**), indicative of their functional activation and potency^23^, we reasoned that the miRNAs upregulated at this stage (profile C) likely supported their effector function, while those transiently downregulated at this time point (profile D) and upregulated thereafter might promote dysfunction. We therefore suggested that preserving the profile C signature at later time points of antigen exposure concomitantly with inhibiting the profile D signature might preserve CAR T cell functionality during the later phase of chronic antigen stimulation. Accordingly, miRNAs from profile C were selected for swapping into the locus of profile D miRNAs via the *castling* procedure.

Among the top candidates from both groups, we selected those displaying the highest fold change of expression on day 6 relative to the pre-stimulation time-point. Notably, most of the top hits from both profiles were miRNA clusters (**Supplementary Table 1**), consistent with their role in coordinating key T cell pathways, as observed in other biological systems^24,25^. As candidates for *castling*, we selected miR-92b-3p together with the miR-17∼92 cluster from profile C, and the miR-15/16 clusters from profile D (**Figure 1c, Supplementary Fig. 2, Table 1**). Supporting the biological relevance of these choices, literature evidence indicated that miR-92b-3p and the whole miR-17∼92 cluster (profile C) are associated with enhanced lymphocyte activation, expansion and persistence^26,27^ whereas the miR-15/16 clusters (profile D) act to constrain T cell proliferation and survival through regulation of apoptotic and cell-cycle genes^28^.

**Table 1:** The role of microRNAs (miRNAs) selected for castling in T cell function.

| miRNA | Function in T cells | References |
| --- | --- | --- |
| miR15/16 cluster | Restricts the proliferation of antigen specific T cells; restricts the survival and early differentiation of CD8+ memory T cells. | 28,85,86 |
| miR-92b-3p | Promotes proliferation | 27,87 |
| miR17~92 cluster | Promotes proliferation, IFN $\gamma$ production (critical for Th1 response and CD4+ T cell antitumor response); prevents activation induced death of CD4+ T cells | 26,88,89 |

### Development of TRIPLE: a robust genome editing strategy enabling miRNA *castling*

*Castling* requires coordinated knock-out (KO) of the dysfunction-associated miRNA (i.e., from profile D) and simultaneous knock-in (KI), within the same locus, of a protective miRNA counterpart (i.e., from profile C). To achieve this with high efficiency and precision, we developed a novel method to maximize the efficiency of homology-directed repair (HDR) and to allow *castling* to occur in a single, high-fidelity editing step.

We first optimized the KO step for the inactivation of the selected detrimental miR-15/16 cluster, present as two copies in the human genome (miR-15a/16-1 on chromosome 13 and miR-15b/16-2 on chromosome 3). We tested multiple targeting designs, prioritizing an approach that employed two flanking CRISPR-Cas9 complexes to remove the entire cluster regions from both genomic loci simultaneously (**Supplementary Fig. 3a-b; Supplementary Table 2**). For the chromosome 3 cluster, we found that deleting miR-16-2 alone (using gRNAs g8 and g9, **Supplementary Fig. 3b**) was sufficient to abolish expression of both miR-16-2 and miR-15b (**Supplementary Fig. 3c-e**), consistent with previously published observations.

We next optimized the conditions for the KI step of *castling*. We chose miR-92b as the initial KI candidate, reasoning that its shorter sequence, relative to the miR-17∼92 cluster, offered a more practical scenario for evaluating the genome-editing strategy. Transcriptomic analysis indicated that among the two miR-15/16 clusters, the one on chromosome 13 was the most transcriptionally active (**Supplementary Fig. 2**). Therefore, we targeted only this site for insertion of the beneficial miR-92b, while simultaneously inactivating the miR-15b/16-2 cluster on chromosome 3.

Using our established T cell genome editing workflow^17,29^, all CRISPR-Cas9 ribonucleoprotein (RNP) complexes were electroporated simultaneously three days after initiation of primary T cell activation, followed by transduction with an AAV6 repair template 30 minutes later (**Figure 2a**). In the absence of the AAV repair template, high editing efficiency at the target locus was observed with ∼97% of alleles harboring indel mutations as measured via small amplicon sequencing (**Supplementary Fig. 4a**). To measure the efficiency of precise HDR-mediated insertion, we used an optimized two-step amplification protocol to overcome the challenges posed by long homology arms that exceed the typical 500 bp limit of short-read next generation sequencing (NGS). To this end, a primary PCR using primers external to the homology arms was followed by a nested PCR generating a short, next generation sequencing-compatible amplicon spanning the editing junction (**Supplementary Table 3**). Using this method, we observed only a modest integration efficiency of miR-92b (∼7%; **Figure 2c, left panel**), prompting development of a novel approach designed to promote HDR-mediated DNA repair. Since after the initial two-RNP excision, we observed the formation of a predominant deletion at the target locus (**Figure 2b** and **Supplementary Fig. 4b**), we hypothesized that adding a third RNP directed to this predominantly deleted region would recurrently generate a double-stranded break, thereby keeping the locus accessible for a longer time and thus promoting HDR-mediated integration of the desired miRNA. We named this approach TRIPLE (for Targeted Replacement Induced by Persistent Locus Editing; **Figure 2b**). Application of TRIPLE led to a striking increase in the insertion efficiency, achieving up to ∼70% precise integration of miR-92b at the miR-15a/16-1 locus on chromosome 13, while preserving efficient simultaneous inactivation of the second miR-15/16 cluster on chromosome 3 (**Figure 2c, right panel**). This insertion rate represented an approximately 10-fold improvement over the canonical procedure and effectively eliminated the reads corresponding to the predominant deletion at the target locus on chromosome 13 (**Supplementary Fig. 4c**). We next attempted *castling* of the larger miR-17∼92 cluster, which encodes six miRNAs (**Supplementary Table 1**) within the same locus. While knockout of both miR-15/16 clusters remained highly efficient using the canonical procedure, integration efficiency of the miR-17∼92 cluster was initially below detection limit. TRIPLE, however, increased the knock-in efficiency to ∼16%, measured via long read amplicon sequencing, demonstrating the scalability of the method also for complex multi-miRNA replacements (**Figure 2c, right panel** and **Supplementary Figure 4d**).

**Figure 2.**
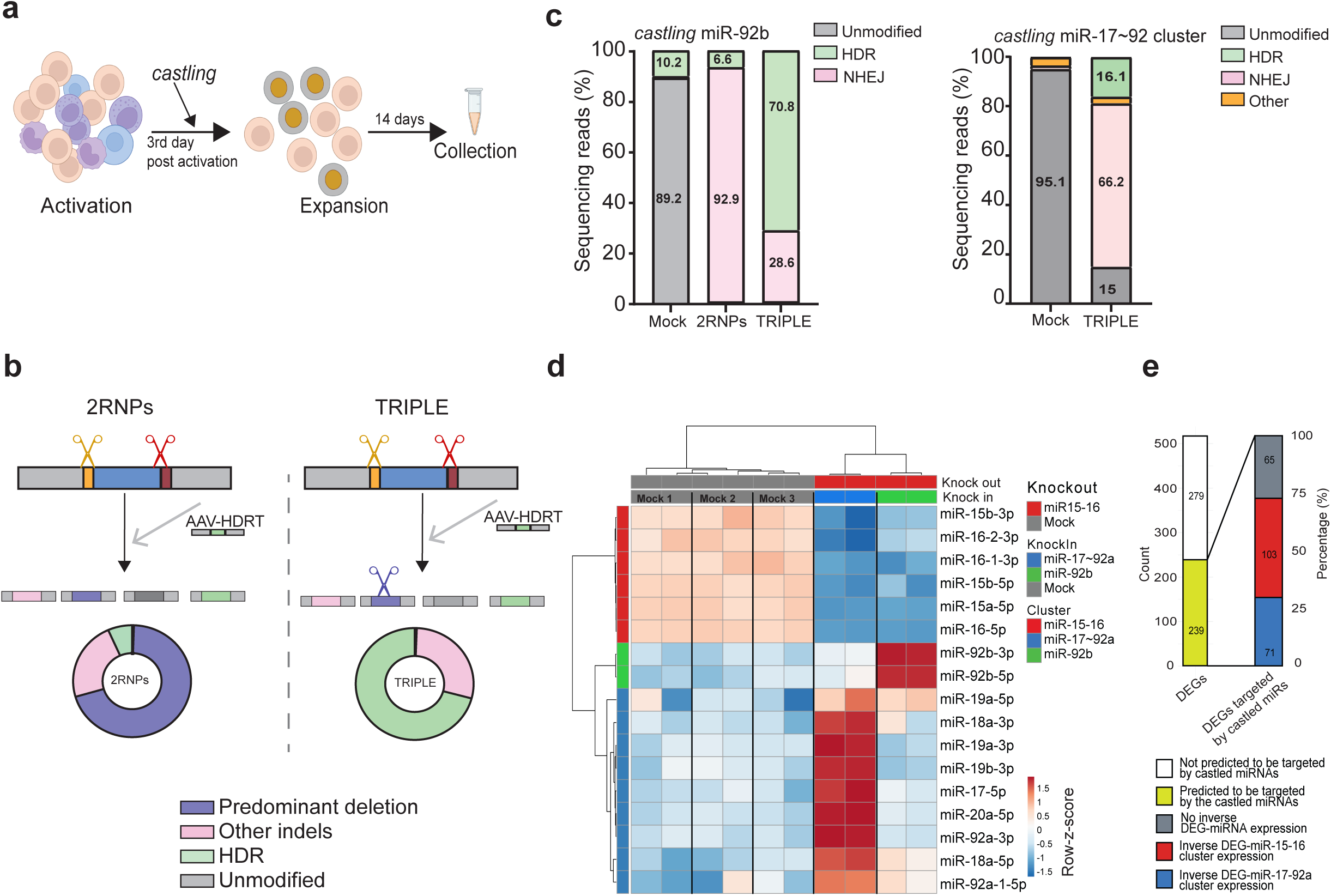
TRIPLE enables efficient miRNA *castling*. **a.** Schematic presentation of the experimental flow. PBMC are thawed and activated as indicated in figure 1a. Three days later, T cells are with the genome editing components for *castling*. Following 14 days of expansion, cells are harvested for subsequent analyisis. **b.** Schematic of the TRIPLE genome editing procedure. In conventional CRISPR–Cas9 editing, two ribonucleoprotein (RNP) complexes target sequences flanking the miRNA locus, resulting in excision of the intervening region and generation of a predominant deletion allele (violet). In the TRIPLE approach, a third RNP is simultaneously delivered to specifically target this predominant deletion junction, promoting recurrent cleavage of the edited locus and thereby increasing the frequency of homology-directed repair (HDR)–mediated insertion using the provided HDR template (HDRT). **c.** The bar graphs indicate the integration efficiency of the miR-92b (left) and of the miR-17∼92 cluster (right) at the miR-15a/16-1 locus on chromosome 13, measured via short or long read sequencing, respectively, in the indicated sample. The numbers within the bars indicate the percentage of sequencing reads of different types: unmodified reads match the normal DNA sequence (grey); reads harboring indel mutations (pink) or precise miRNA sequence integration (green) result from the activation of the non-homologous end-joining (NHEJ) or homology directed repair (HDR) DNA repair pathways, respectively; other (orange) indicate sequences that remained unclassified. **d.** Heatmap showing the scaled (unit variance) expression of the *castled* miRNAs following miR-15/16 KO combined with KI of either miR-17∼92 cluster or miR-92b. Rows and columns were hierarchically clustered using Pearson correlation distance and average linkage. Each pair of columns represents technical replicates of the indicated condition. All mock samples are electroporated without genome editing components and either left non-transduced (Mock 1) or transduced with the respective AAV-HDRT (Mock2: mir92b; Mock 3: miR-17∼92 cluster). **e.** Bar graph summarizing the number of differentially expressed genes (DEGs) identified via bulk RNA-seq in the indicated sample relative to mock controls (FDR < 0.05). DEGs are classified as either not predicted (white) or predicted (yellow) to be targeted by the *castled* miRNAs. The latter are further subdivided into transcripts with expression not inversely correlated with the *castled* miRNA expression (grey), or inversely correlated with miR-15/16 (red) or with miR-17∼92 cluster members (blue).

### Evaluation of coordinate miRNA and mRNA expression profiles in the *castled* T cells

To demonstrate that miRNA *castling* can effectively reprogram gene expression networks, resulting in the anticipated coordinated changes in expression levels of the edited miRNAs and their corresponding target mRNAs, we subjected the *castled* primary T cells described above to a combined bulk miRNA and mRNA sequence analysis. The analysis was performed at the time point typically used as day 0 (pre-antigen stimulation) in our *in vitro* CAR T chronic antigen stimulation assay (**Figure 1b**). Notably, at this pre-stimulation phase, miRNAs of profile D (e.g., miR-15/16 clusters) are already expressed at higher level as compared to the majority of those belonging to profile C (**Figure 1c** and **Supplementary Figure 2**).

We next focused on the analysis of expression levels of the specific miRNAs used for *castling*. In line with the expectations, edited T cells displayed strong downregulation of all miR-15/16 cluster members and a corresponding increase in expression of the inserted miRNAs, with miR-92b reaching a 4.4-fold increase, while the individual members of the miR-17∼92 cluster were upregulated between 1.6-fold and 2.6-fold relative to mock controls (**Supplementary Fig. 5, left panel**), consistent with the higher insertion efficiency of the single miRNA compared to the miRNA cluster (70.8% vs 16.1%, respectively; **Figure 2c)**. Importantly, all miRNA isoforms (isomiRs) derived from the targeted clusters followed the same regulatory direction, demonstrating precise modulation across both canonical and variant sequences (**Supplementary Fig. 5, right panel**).

To determine whether these changes were accompanied by coordinated mRNA responses, we analyzed global mRNA expression in the same samples in the cells where miR-17∼92 cluster was inserted into the miR-15a/16-1 locus on chromosome 13, as this configuration was expected to elicit broader downstream effects than castling of the single miR-92b. Among more than 500 statistically significant (FDR < 0.05) expressed genes identified in the *castled* samples as compared to the mock control sample, approximately half of them were predicted targets of the *castled* miRNAs. Of these, ∼75% exhibited inverse regulation relative to their corresponding miRNA partners, consistent with canonical miRNA-mediated mRNA repression (**Figure 2f**). The lists of genes identified as inversely regulated compared to either miR-15/16 or miR-17∼92 cluster members are displayed in **Supplementary Tables 4** and **5**, respectively. These findings confirm that *castling* of selected miRNA sequences rewires downstream gene expression, largely in the expected direction. Next, we investigated whether molecular changes elicited by *castling* in primary T cells translate into measurable functional benefits when applied to engineered CAR T cells exposed to chronic antigen stimulation.

### *Castling* of miR-17∼92 and miR-15a/16-1 elicits mRNA expression profile supporting CAR T effector function and delays the onset of CAR T cell dysfunction under chronic antigen stimulation

As indicated above, the functional impact of *castling*-mediated gene networks rewiring in CAR T cells was assessed by focusing on insertion of the miR-17∼92 cluster into the miR-15a/16-1 locus on chromosome 13, with simultaneous inactivation of the second miR-15/16 cluster on chromosomes 3. We hypothesized that *castling* would sustain CAR T cell effector function during chronic antigen stimulation by maintaining high miR-17∼92 expression through its integration at the miR-15a/16-1 locus, while simultaneously suppressing the dysfunction-promoting miR-15/16 program via genomic knockout.

The experimental design followed the outline shown in **Figure 1b** with *castling* performed using the established TRIPLE-based workflow two days after the CAR cassette transduction. Edited and control cells were expanded for 12 days prior to initiation of chronic antigen stimulation and were harvested for analysis before antigen exposure (day 0) and after 6 and 10 of stimulation. Genome editing efficiency was assessed via long-read sequencing prior to initiation of the chronic antigen stimulation, revealing that approximately 50% of reads showed evidence of indel mutations as a result of nuclease activity, indicative of efficient inactivation of the targeted miR-15a/16-1 locus. Additionally, ∼15% of the reads corresponded to precise knock-in events of the miR-17∼92 cluster at the chromosome 13 target site, consistent with previous results in primary T cells (**Figure 3a**).

**Figure 3.**
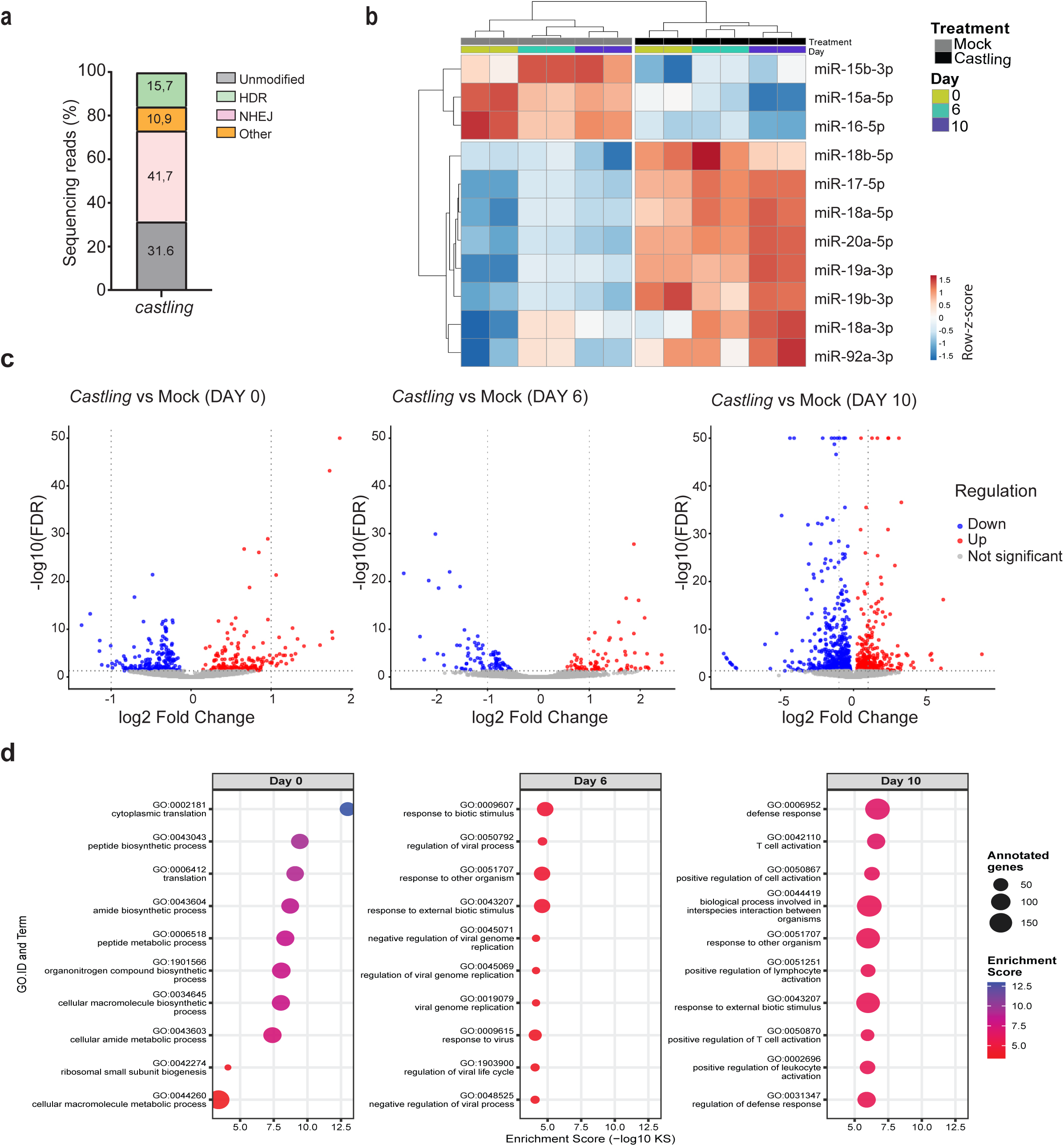
Effect of miR-17∼92/miR-15a/16-1 *castling* in CAR T cells subjected to chronic antigen stimulation. **a.** Bar graph showing genome editing efficiency for the *castling* of the miR-17∼92 cluster into the miR-15a/16-1 locus, measured via long read amplicon sequencing. Reads are classified as indicated in figure 2c. **b.** Heatmap showing the expression level of the *castled* miRNA measured via small RNA sequencing in the indicated samples. Each group of two columns corresponds to two technical replicates of the indicated sample. **c.** Volcano plots (upper) depicting differential gene expression between *castled* and mock-transfected CAR T cells at the indicated time points, shown as log2 fold change (x-axis) versus −log10 false discovery rate (FDR; y-axis). The top 10 enriched Gene Ontology (GO) biological process terms, identified among differentially expressed genes at each time point using a Kolmogorov–Smirnov (KS) enrichment test, are show in the lower panels.

To track miRNA expression dynamics, miRNA sequencing was performed at days 0, 6, and 10. Consistent with the *castling* strategy, edited CAR T cells displayed an efficient ablation of the endogenous miR-15/16 clusters expression, accompanied by a sustained upregulation of all miR-17∼92 cluster members (**Figure 3b**). Unlike their physiological transient expression in mock controls, which peaked at day 6 and declined thereafter, in the edited cells, the expression of miR-17∼92 cluster members remained persistently elevated while miR-15/16 clusters expression was suppressed throughout the assay (**Supplementary Fig. 6**), supporting the underlying hypothesis that *castling*-mediated rewiring of promoter control can maintain beneficial regulatory programs over time.

Functionally, these molecular changes translated into measurable performance gains of the *castled* CAR T cells. During repeated antigen exposure, *castled* CAR T cells maintained >90% tumor cell killing ability up to day 8 of antigen stimulation, as measured by counting the remaining NALM6 cells at the specific time points before replenishing them, while control CAR T cells showed an early functional decline, with cytotoxicity falling to ∼75% by that time (**Supplementary Fig. 7a**). In line with their enhanced cytolytic potential, flow cytometric analysis at the assay end point revealed a modest but consistent increase in the frequency of CD8⁺ effector T cells in the *castled* sample (**Supplementary Fig. 7b**).

To assess the impact of *castling* on CAR T cell gene expression, we performed bulk mRNA sequencing and visualized differentially expressed genes in volcano plots comparing *castled* cells to their respective mock control counterparts at each time point (**Figure 3c, upper panel**). Consistent with progressive cellular remodeling induced by persistent antigen stimulation, the number of significantly differentially expressed genes (FDR <0.05) increased over time, reflecting the dynamic reshaping of the transcriptional landscape as the assay progressed. This observation was also confirmed by the principal component analysis (PCA) demonstrating substantial gene expression differences between the two cell subsets on day 10 (**Supplementary Fig. 8**).

Gene Ontology (GO) analysis provided functional insight into these transcriptional changes (**Figure 3c, lower panel**). At day 0, the primary differences between *castled* and control cells were observed in metabolic and biosynthetic processes. By the intermediate day 6 time point, enriched GO terms predominantly reflected responses to antigenic stimuli, including viral, biotic, and other organisms (**Figure 3c, lower panel**) suggesting that *castled* cells were more reactive relative to controls. At the endpoint (day 10), numerous GO terms related to T cell activation were strongly enriched, in agreement with the increased proportion of CD8⁺ cells at this time point (**Supplementary Fig. 7b**) and consistent with the rewired cells adopting a more terminally differentiated cytotoxic phenotype.

At this latter time point, about 500 genes were significantly deregulated (FDR <0.05) with expression changes of at least 2-fold (**Supplementary Table 6**). Several genes known to enhance CAR T cell effector function were upregulated, including those associated with increased invasiveness (*CCL5 –* ∼5-fold upregulation)^30^, improved antigen recognition (*KLRC4*, *AOAH* – ∼3-4-fold upregulation)^31,32^, enhanced cytotoxicity (*CD8A/B*, *ITGAE*, *SYTL2*, *GNLY*, *NKG7*, *SLAMF7, IL2RB – ∼*2-4-fold upregulation)^33–39^, improved survival and persistence (*KIT, CD27, SLAMF6* - ∼2-9-fold upregulation)^40–42^, enhanced proliferation and heightened activation (*JAML, CRACR2A, CD63 –* ∼2-3 fold upregulation)^43–45^. Whereas upregulation of CD8A/B could be a reflection of a higher proportion of CD8+ cells among the *castled* ones (**Supplementary Fig. 7b**), the detected 4-fold and 7-fold significant upregulation of *TRGC1* and *TRGC2*, respectively, in the *castled* cells could be a reflection of a higher proportion of γδ T-cell lineage known for their contribution to rapid cytotoxicity, cytokine secretion and tissue surveillance^46^. Conversely, a notable set of genes linked to Type I interferon (IFN) response (*IFI44L*, *IFI44*, *IFIT3*, *MX2*, *RSAD2* and *OAS2*), known for hampering CAR T cell antitumor function^47,48^, were downregulated up to 20-fold. Additionally, expression of most of the IFN response genes mentioned above has been individually confirmed as detrimental for CAR T effector function^49–52^. Among other known genes whose damped expression has been previously associated with improved T cell effector function and persistence, *SOX4*^16^, *LGALS1*^53^*, SOCS1*^54^*, LGALS9*^55^*, ARG2*^56^*, PTGS2*^57^*, SESN3*^58^, and *CD38*^59^ were found downregulated up to 8-fold.

Together, these results provide proof-of-concept that *castling* of rationally selected miRNAs can functionally reprogram CAR T cells, sustaining effector capacity under chronic antigen stimulation.

## Discussion

Our results demonstrate that *castling* can rewire complex pathology-driven gene regulatory networks to sustain effector function of engineered CAR T cells under chronic antigen stimulation compared to non-modified CAR T cells. This entails the coordinated upregulation of numerous beneficial genes and downregulation of detrimental ones, in line with several literature reports correlating the expression of these genes with enhancement of specific aspects of CAR T or T cell function. By orchestrating these changes simultaneously, this approach achieves a holistic reprogramming that extends beyond conventional strategies for CAR T cells enhancement.

*Castling* involves a single genome editing step, in which miRNA sequence(s) that counteract the disease state and are insufficiently expressed under pathological conditions are inserted into the genomic locus of a disease-promoting miRNA. Because this locus is transcriptionally activated in the disease state, *castling* simultaneously suppresses the pathogenic program while inducing a beneficial one (**Figure 1a**). Hence, the potential therapeutic gain of *castling* is two-fold: (1) activation of the “therapeutic program” through a disease-triggered promoter, thus ensuring its execution under pathological conditions of interest via using the disease’s own transcriptional machinery against itself; and (2) inhibition of the “disease program” that is usually triggered by pathology from the unmodified locus into which insertion has been performed. Although not addressed in this study, a third advantage - an automatic winding down of the *castled* therapeutic program once the pathological trigger has gone - can be theoretically anticipated.

To enable efficient *castling* at the genomic level, we developed TRIPLE (Targeted Replacement Induced by Persistent Locus Editing). This novel genome editing procedure uses two RNPs to generate a defined excision at the target locus, creating a predominant and predictable deletion allele. A third RNP is then directed to this newly formed junction, enabling recurrent cleavage of the edited chromosome. Although the precise mechanism remains to be fully elucidated, this repeated targeting likely prolongs the window during which the locus remains accessible to homology-directed repair, thereby markedly improving knock-in efficiency. Importantly, the conceptual simplicity of TRIPLE, unlike previous double-tap or recursive editing strategies relying on microhomology^60,61^, suggests that it should be broadly applicable to diverse genomic contexts. Importantly, other emerging genome engineering technologies could, in principle, also be adapted to achieve targeted sequence insertion. These include e.g. prime editing^62^, CRISPR-associated transposases (CAST)^63^, recombinase- or integrase-based systems such as PASTE^64^, and homology-independent targeted integration (HITI)^65^. Each of these platforms offers distinct advantages with respect to cargo size, repair pathway usage, or genomic flexibility. However, they mostly require specialized molecular components, complex optimization, or non-canonical enzymes that may limit immediate applicability across diverse loci and cell types as well accessibility for a broad community of researchers. In contrast, TRIPLE builds on well-established CRISPR–Cas9 editing workflows and achieves substantial gains in knock-in efficiency through the simple addition of an extra guide RNA, making it straightforward to implement, broadly accessible, and readily compatible with existing *ex vivo* cell engineering pipelines.

Our data confirmed that *castling* could efficiently place the selected “therapeutic” miRNAs (miR-92b or the members of the miR-17∼92 cluster) under the regulatory elements of the target “pathogenic” miRNA (miR-15a/16-1) while disrupting the expression of the latter with concomitant inverse regulation of numerous mRNAs predicted to be targeted by the *castled* miRNAs. Importantly, *castling*-dependent rewiring can sustain beneficial regulatory programs over time, and, unlike their transient physiological expression in non-edited cells, the *castled* miR-17∼92 cluster members remained elevated in modified CAR T cells throughout chronic antigen stimulation. In our experiment, being performed in unstimulated T cells, *castling* also likely “primed” CAR T cells obtained from this cell population, so that upon the onset of chronic antigen-stimulation elicited dysfunction, the protective miRNAs could be rapidly and robustly upregulated from the disease(pathology)-activated promoter to levels far above normal, positioning the cells to sustain beneficial gene expression programs when they are mostly needed.

Notably, the model of CAR T cell dysfunction used in our study mimics a well-known phenomenon of CAR T cell exhaustion, the phenotype that they typically acquire upon entering the tumor microenvironment as shown in multiple instances and that limits their therapeutic potential^16–18^. However, we emphasize that the *castled* CAR T cells presented here are not yet optimized for clinical use; the primary aim of these experiments was to provide a technical proof of concept for the feasibility of this novel methodology. The cells have not yet been tested *in vivo,* which will be addressed in future studies. Furthermore, although the observed gene expression changes and functional assays support enhanced effector function of the *castled* cells, the optimal miRNA pairs or clusters for *castling* may remain to be identified. In addition, the efficiency and specificity of *castling* may be further improved beyond what was achieved in the present study, for example through optimization of repair templates^66^ or the use of genome-editing systems with enhanced homology-directed repair activity^67^.

In conclusion, *castlin*g provides a versatile framework for miRNA-guided network reprogramming, with TRIPLE enabling high-efficiency, precise genome editing not only for single miRNAs but also for miRNA clusters. By harnessing endogenous regulatory logic to modulate gene networks, *castling* enables single-event, multi-target tuning of cellular programs, offering a versatile blueprint for engineering resilient cell therapies. Beyond potential implications in improving CAR T cell therapy, miRNA *castling* may also be used *ex vivo* for improvement of other therapeutically relevant cell types, including macrophages, NK cells, Tregs and various types of stem cells. With appropriate delivery systems, this approach holds a transformative potential also for *in vivo* interventions against complex chronic diseases, such as neurodegeneration, cardiovascular disorders, autoimmune diseases and fibrosis, driven by complex regulatory gene expression imbalances. Finally, *castling* may be applicable not only for swapping “pathogenic” and “therapeutic” miRNAs but also of “pathogenic” and “therapeutic” protein coding mRNAs either between them or with miRNA if and when it deems therapeutically appropriate.

## Methods

### Cell lines and primary T cell culture conditions

Peripheral blood mononuclear cells (PBMCs) from healthy donors were obtained from leukoreduction system (LRS) chambers and isolated by density gradient centrifugation using Biocoll (Bio&SELL, Nuremberg, Germany). Primary T cells were obtained from PBMCs either by enrichment after activation or by negative selection using the EasySep™ Human T Cell Isolation Kit (Stemcell Technologies, Vancouver, Canada) to yield a pure CD3⁺ cell population. T cells or CAR T cells were cultured in complete RPMI medium consisting of RPMI 1640 GlutaMAX (Life Technologies, Carlsbad, CA), 10% FBS (PAN Biotech, Aidenbach, Germany), 1X penicillin–streptomycin, and 10 mM HEPES buffer (Sigma-Aldrich, St. Louis, MO). Unless stated otherwise, this medium was supplemented with 100 U/mL recombinant human IL-2 (rhIL-2; ImmunoTools, Friesoythe, Germany). Medium was replaced every 2–3 days depending on the rate of cell expansion. NALM6 cells naturally expressing the CD19 antigen were transduced with a lentivirus to express GFP (NALM6-GFP⁺), as described previously^17^, and cultured in the same medium as T cells but without rhIL-2 supplementation. All cells were maintained at 37 °C in a humidified atmosphere containing 5% CO₂.

### T cell manipulation

Primary T cells (from PBMCs or isolated CD3⁺ cells) were activated in activation medium consisting of complete RPMI supplemented with ImmunoCult™ Human CD3/CD28/CD2 T Cell Activator (Stemcell Technologies, Vancouver, Canada) at a final concentration of 10 µL/mL and rhIL-2 at 100 U/mL. Cell density was kept at 2×10⁶ cells/mL. Afterwards, T cells were either subjected to *castling* via genome editing or first transduced to express a CD19-specific CAR prior *castling*. For the genome editing procedure, activated T cells derived from PBMCs were electroporated on day 3 post-activation using the 4D-Nucleofector™ device (Lonza, Basel, Switzerland) following our established protocol^17^. Briefly, all ribonucleoprotein (RNP) complexes were pre-assembled by mixing 75 pmol of the guide RNA (GeneScript, Piscataway, NJ) with 18 pmol of Alt-R® S.p. Cas9 Nuclease V3 or Alt-R® S.p. HiFi Cas9 Nuclease V3 (Integrated DNA Technologies, Inc., Coralville, Iowa) and incubated for 10–12 minutes at room temperature. Each RNP complex targeted to each selected site (**Supplementary Table 2**) was prepared separately and combined only immediately before electroporation. After the electroporation pulse, cells were transferred into 96-well plates containing 180 µL of fresh complete RPMI supplemented with 1,000 U/mL rhIL-2 for optimal cell recovery. After 30 minutes, the electroporated cells were transduced with an adeno-associated virus (AAV) serotype 6 encoding a suitable repair template (**Supplementary Table 7**) at a genome copy number of 5×10⁴ GC/cell to achieve the selected *castling* event. Over the next four days, the rhIL-2 concentration was gradually reduced to 100 U/mL. The edited cells were expanded for a total of 14 days and then frozen in CryoStor® CS10 (STEMCELL Technologies, Cologne, Germany) for further use. For the generation of CD19-specific CAR T cells we used a VSV-G pseudotyped lentiviral vector encoding for a second-generation CD19-specific CAR driven by the phosphoglycerate kinase-1 promotor (PGK) promoter. The CAR expression plasmid was generated via Gibson assembly and included the CD19-specific scFV sequence, followed by a CD8 spacer, the 4-1BB/CD3 zeta stimulatory domain and linked via P2A element to a truncated LNGFR used to enrich for CAR+ cells after transduction. The lentiviral particles were purchased using the VSV-G Lentivirus Large-scale Packaging Service for custom vectors (VectorBuilder Inc., Chicago, Illinois). For transduction, 1×10⁶ CD3⁺ cells were activated as described above and plated in one well of a 48-well plate in activation medium with 1,000 U/mL rhIL-2. Lentiviral particles were added 24 hours later at 5 transducing units (TU)/cell, after incubation with Polybrene (final concentration 5 µg/mL; VectorBuilder Inc., Chicago, Illinois) for 5 minutes at room temperature to enhance transduction efficiency. The plate was then spinoculated at 1,600 RCF for 90 minutes to concentrate the virus onto the cell surface. Transduced cells were maintained in culture for two days prior the genome editing procedure conducted as described above.

### Cell surface staining and flow cytometry

Cell surface staining was performed using fluorochrome-conjugated primary antibodies for direct detection by flow cytometry. Between 5×10⁴ and 1×10⁵ cells were resuspended in the appropriate antibody mix following the manufacturers’ instructions. The following antibodies were used: anti-human CD271 (LNGFR)-APC (clone ME20.4-1.H4, Miltenyi Biotec, Bergisch Gladbach, Germany), anti-human CD25-PE (clone 4E3, Miltenyi Biotec). anti-human CD279-FITC (clone NAT105, BioLegend, San Diego, CA), anti-human CD62L-BV421 (clone DREG-56, BD Horizon, Franklin Lakes, NJ), anti-human CD45RA-FITC (clone HI100, BioLegend), anti-human CD39-BV711 (clone A1, BioLegend), anti-human CD127 (IL-7Rα)-PE/Cy5 (clone A019D5, BioLegend).

### Chronic antigen exposure (CAE) assay

Transduced or control T cells were expanded for 14 days in culture with 100 U/mL rhIL-2, with medium exchange every 2–3 days as needed. On the day of assay initiation, LNGFR⁺ cells (indicative of CAR+ cells as both proteins were co-expressed from the same lentiviral expression cassette) were enriched using the MACS® Cell Separation System and the CD271 (LNGFR) MicroBead Kit (Miltenyi Biotec, Bergisch Gladbach, Germany). This ensured initiation of the assay with a purified (i.e. >95%) CAR⁺ population. The CAE consisted of co-culturing CD19-specific CAR T cells with CD19⁺/GFP⁺ NALM6 cells at a 1:4 effector-to-target (E:T) ratio. As the co-culture progressed, the number of target cells was adjusted to maintain this ratio. Every two days, cells were counted, stained for LNGFR, and analyzed by flow cytometry to determine the number of CAR⁺ (effector) and GFP⁺ (target) cells. Based on these counts, the appropriate number of GFP⁺ NALM6 cells was added to sustain the 1:4 ratio. Effector and target cell counts were recorded throughout the assay to monitor CAR T cell proliferation and infer cytotoxic activity. To this end, the number of CAR T cells were quantified as live LNGFR+ cells and cytotoxic activity was calculated as the percentage of live target cells remaining at each time point, relative to the number of target cells initially seeded. The duration required to induce key T cell exhaustion features varied among donors (typically 10–14 days). For miRNA and mRNA sequencing, CAR T cells were separated from the fluorescent target cells by FACS as GFP⁻ cells (and cryopreserved on days 6, 10, 12, or 14 of the CAE assay). Day 0 samples were collected immediately before assay initiation. Endpoint analyses were performed at the time point when all samples exhibited decrease of the total CAR T cell population (i.e. days 12-14 of the CAE). These analyses included: surface staining for exhaustion and stemness markers (CD39 and IL-7R, respectively); assessment of cytotoxicity by quantifying via flow cytometry the amount of GFP⁺ cells remaining after an additional 24-hours co-culture at a 1:4 E:T ratio; measurement of cytokine release (IFN-γ and IL-2) using the BD™ Cytometric Bead Array (CBA) Human Soluble Protein Master Buffer Kit (BD Biosciences, Franklin Lakes, NJ), according to the manufacturer’s instructions.

### Sample preparation for the measurement of genome editing efficiency

To determine genome editing efficiency, the regions surrounding the nuclease target sites were PCR-amplified and subjected to next-generation sequencing analysis. In brief, genomic DNA was extracted from frozen pellets of approximately 500,000 cells using a Nucleospin Tissue Extraction Kit (Macherey-Nagel, Düren, Germany). Long or short PCR amplicons were obtained from 100 ng of genomic DNA using the primers indicated in **Supplementary Table 3**. For nested PCR, the long amplicon PCR product was separated on an agarose gel, and the desired band was purified via gel extraction (Qiagen, Germantown, MD 20874, USA). Approximately 20ng of the purified PCR product was used as the input for the nested PCR using the primers indicated in **Supplementary Table 3**. All PCRs were performed with Q5 HotStart polymerase (New England Biolabs GmbH, Frankfurt am Main, Germany), following the manufacturer’s instructions.

### Quantification of genome editing efficiency

Genome editing efficiency was evaluated by subjecting the PCR amplicons described above to either small or long amplicon sequencing. The small amplicons were used to determine the frequency of indel mutations at the miR-15/16 clusters target sites as well as the precise integration of a small insert (i.e. the mir92b sequence) at the miR-15a/16-1 target on chromosome 13. To this end, small amplicons (500ng of purified PCR product) were sequenced using Illumina paired-end sequencing at Genewiz (Leipzig, Germany) and data analyzed using Crispresso2 (v2.0.20b) with default settings. To quantify the extent of large deletions at the miR-15a/16-1 locus on chromosome 13 and the integration of the large miR-17∼92 cluster cassette, long read amplicon sequencing was performed using Oxford Nanopore Technology (Microsynth AG, Balgach, Switzerland). The raw reads were filtered using filtlong (version 0.2.1) (min_length 4500) to remove short reads. We then created a BLAST database using the reads of the sample with the “makeblast nucl” command. A series of 6 landmark sequences (the 2 PCR primers, 2gRNAs, the entire inserted sequence and the entire wild-type sequence; **Supplementary Table 8**) were aligned to the sample’s database using BLAST (version 2.16.0) using the following parameters: word_size=5, evalue=1, perc_identity=90, dust=no, gapopen=2, gapextend=10, num_alignements=10000000. We parsed the BLAST results of each read and determined the presence of a landmark sequence on the read if the read was successfully aligned to that landmark sequence. In that manner, we evaluated the presence or absence of each landmark sequence in each of the reads. To select complete amplicons, we filtered out all reads that did not have both PCR primer sequences. We lastly sought out to classify reads into 4 groups. The reads containing the wild-type sequence, both gRNA target sequences but not the insertion were classified as “unmodified” reads. The reads missing the gRNA target sequences but containing the inserted sequence were classified as “edited” or “HDR” to indicate that presumably these alleles result from the activation of the HDR-repair pathway leading to precise knock-in events. The reads missing the insertion sequence and at least 1 gRNA target sequence were classified as “cut” or “NHEJ” to indicate that presumably these alleles result from the activation of the NHEJ-repair pathway leading to the formation of variable indel mutations. Lastly, the remaining reads were classified as “other”.

### Total RNA extraction

Total RNA was extracted from frozen cell pellets containing 10,000–50,000 cells using the miRNeasy Mini Kit (Qiagen, Germany) according to the manufacturer’s instructions. Cells were lysed in 1,000 µl QIAzol reagent by mixing on a vortex for 10 seconds. Phase separation was achieved by adding 200 µl chloroform, followed by centrifugation. RNA was precipitated with ethanol in the presence of 50 µg/ml glycogen to enhance the RNA yield. Washing steps were performed automatically on a QIAcube instrument (Qiagen, Germany). RNA was eluted in 30 µl nuclease-free water. RNA concentration and integrity were assessed by spectrophotometry (NanoDrop, absorbance at 260 nm) and microcapillary electrophoresis using the High Sensitivity RNA Kit together with the Fragment Analyzer (Agilent, United States). Purified total RNA samples were stored at –80 °C until further processing.

### Measurement of small RNA expression levels

The expression levels of mature miRNA shown in **Supplementary Figure 3c-e** were measured using the TaqMan™ Small RNA Assays” (Applied Biosystems) followed by real time quantitative PCR using TaqMan® Fast Advanced Master Mix (Applied biosystems). In brief, total miRNA was isolated from cells using the miRNeasy Mini Kit (Qiagen) according to the manufacturer’s instructions. The selected miRNAs were reverse transcribed using TaqMan™ MicroRNA Reverse Transcription Kit (Applied Biosystems) and quantified using the specific assay IDs: 000389 (hsa-miR-15a-5p), 000391, (has-miR-16-5p), 000390 (hsa-miR-15b-5p) and 002277 (hsa-miR-320a). To obtain relative expression levels, normalization against the reference miRNA 320a was chosen using the 2^-ΔΔCt method^68^. For next generation small RNA sequencing, miRNA libraries were prepared from 100 ng total RNA using the RealSeq® Small RNA Library Preparation Kit (RealSeq Biosciences) as previously described^69^. Adapter-ligated and reverse-transcribed libraries were quantified by qPCR to determine the optimal number of PCR amplification cycles (17–19 cycles). Amplification was performed using barcoded Illumina reverse primers in combination with the Illumina forward primer. Library quality was assessed using either the DNA 1000 Chip or DNA Fragment Analyzer reagents (Agilent, USA). Equimolar pooling of individually barcoded libraries was performed prior to sequencing on an Illumina NextSeq 2000 instrument with 100 bp single-end reads. Sequencing data were processed using the miND analysis pipeline previously reported^70^. Briefly, reads were demultiplexed, and quality was assessed using FastQC and MultiQC. Adapter trimming and quality filtering were performed with Cutadapt, retaining reads ≥17 nt in length. Reads were mapped to the human genome (GRCh38.p14, Ensembl) using Bowtie v1.2.2 allowing up to two mismatches, followed by mapping to miRBase v22.1 (https://www.mirbase.org/) allowing one mismatch, via miRDeep2 v2.0.1.2. Statistical analyses were performed in R v3.6 using the packages pheatmap v1.0.12, pcaMethods v1.78, and genefilter v1.68. Differential expression analysis was conducted with edgeR v3.28, employing quasi-likelihood negative binomial generalized log-linear models. To enhance false discovery rate (FDR) control, the independent filtering approach of DESeq2 was adapted for edgeR to remove low-abundance miRNAs prior to testing.

### mRNA sequencing

mRNA-seq libraries were prepared from 100 ng of total RNA using the QuantSeq 3′ mRNA-Seq Library Prep Kit FWD for Illumina (Lexogen, Austria) according to the manufacturer’s instructions. After library purification, yields were quantified using the Fragment Analyzer (NGS Fragment Kit, Agilent Technologies). Individually barcoded libraries were pooled at equimolar concentrations and sequenced on an Illumina NextSeq 2000 instrument with 100 bp single-end reads. Sequencing data quality was evaluated using FastQC and MultiQC. Adapter trimming and quality filtering were performed using the BBDuk tool (part of the BBMap package, v38.69), retaining reads with a minimum length of 17 nt and Phred quality ≥30. Reads were aligned to the human reference genome (GRCh38.p14, Ensembl) using STAR v2.7, and indexed with SAMtools v1.9. Gene-level read counts were obtained using HTSeq-count v0.13.

### Visualization of microRNA and mRNA expression analysis

Heatmaps and principal component analysis (PCA) plots were produced on the basis of ln-transformed RPM data using clustvis^71^. For heatmaps unit variance scaling was applied to RPM values and average linkage together with Pearson correlation clustering were used as distance measures. MA plots were produced in Microsoft Excel using log2 fold changes and logCPM values derived from EdgeR differential expression analysis. Volcano plots were generated in R (version ≥4.2) using the *ggplot2* package. Differential expression results were visualized by plotting log2 fold change against −log10 false discovery rate (FDR). Genes were classified as significantly up-regulated or down-regulated using thresholds of FDR < 0.05 and were colored accordingly.

### microRNA target prediction and Gene Ontology (GO) enrichment analysis

Target genes for differentially expressed microRNAs were identified using multimiR v1.30.0^72^. This tool integrates eight miRNA target prediction databases: DIANA^73^, ElMMo^74^, MicroCosm^75^, PITA^76^, miRanda^77^, PicTar^78^, TargetScan^79^ and miRDB^80^, as well as three databases with experimentally validated targets: miRTarBase^81^, miRecords^82^ and TarBase^83^. Only target genes found in at least three of the 11 databases were retained for downstream analysis.

To identify potential functional miRNA-mRNA interactions, the miRNA results were integrated with differential expression data from mRNA sequencing. The analysis focused exclusively on inverse regulatory relationships, pairing upregulated miRNAs with downregulated mRNAs and downregulated miRNAs with upregulated mRNAs. Differentially expressed mRNAs (FDR ≤ 0.05) between the castled and mock transfected cells at day 0, day 6 or day 10 were used in a GO-term enrichment analysis. Enriched biological processes (BP) were identified by using the Kolmogorov Smirnov (KS) test with the unadjusted p-values from the differential expression analysis as rank information with the tool topGO^84^. Gene Ontology (GO) enrichment results were visualized using dot plots generated in R (version ≥4.2) with the ggplot2 package. For each analysis timepoint, the ten most significantly enriched biological process GO terms were selected based on their enrichment score, defined as the negative logarithm (base 10) of the Kolmogorov–Smirnov (KS) test p-value. Dot plots display GO terms on the y-axis and enrichment scores on the x-axis. Dot size corresponds to the number of annotated genes associated with each GO term, while dot color represents the enrichment score using a continuous red-to-blue gradient. GO terms were ordered within each timepoint from highest to lowest enrichment score and displayed side-by-side to enable comparison across timepoints.

### Data availability

The sequencing data generated in this study are deposited in the gene expression omnibus (GEO) database with the accession number GSE316480.

## Supporting information

Antony_et_al_SuppleentaryData

## Acknowledgments

We would like to thank all the members of the Institute for Transfusion Medicine and Gene Therapy’s R&D department, Juliana Friedman of Lepton Pharmaceuticals and Dr. Elena Sotillo, Senior Research Scientist at Stanford Cancer Institute (Stanford University, Palo Alto, USA) for productive discussions. Special thanks to Antonio Carusillo, Jamal Alzubi, Sibtain Haider, Michalis Papacharalampous and Melissa Whitehead for scientific advice. We are grateful to the FREEZE Biobank, the Blood Donation Center, and the Lighthouse Core facility, all Medical Center-University of Freiburg, for provision of the LRS chambers containing the PBMCs of healthy donors, or help with flow cytometry, respectively.

## Author contributions

E.F and C.M. conceived the ideas of *castling* and TRIPLE, respectively. D.A, M.S.R.A, H.K., S.A, S.A.N., E.S., D.Z. and C.M. designed the study. D.A, M.S.R.A, M.P., M.R., A.T. and A.D. performed experiments. D.A, M.S.R.A, H.K., S.A., M.P., A.T., A.D., G.A., M.H., D.Z., E.F. and C.M. analyzed data. M.B., T.I.C., D.Z., T.C. and C.M. acquired funding. D.Z., E.F. and C.M. wrote the paper with input from all authors.

## References

1 Lee, T. I. & Young, R. A. Transcriptional regulation and its misregulation in disease. Cell 152, 1237–1251 (2013). 10.1016/j.cell.2013.02.014

2 Wang, J., Liu, Q. & Shyr, Y. Dysregulated transcription across diverse cancer types reveals the importance of RNA-binding protein in carcinogenesis. BMC genomics 16 Suppl 7, S5 (2015). 10.1186/1471-2164-16-S7-S5

3 Hu, Z. et al. Hepatocellular carcinoma cell-derived exosomal miR-21-5p promotes the polarization of tumor-related macrophages (TAMs) through SP1/XBP1 and affects the progression of hepatocellular carcinoma. Int Immunopharmacol 126, 111149 (2024). 10.1016/j.intimp.2023.111149

4 Kim, J. et al. TLR7 activation by miR-21 promotes renal fibrosis by activating the pro-inflammatory signaling pathway in tubule epithelial cells. Cell Commun Signal 21, 215 (2023). 10.1186/s12964-023-01234-w

5 Kurowska-Stolarska, M. et al. MicroRNA-155 as a proinflammatory regulator in clinical and experimental arthritis. Proc Natl Acad Sci U S A 108, 11193–11198 (2011). 10.1073/pnas.1019536108

6 Pasculli, B. et al. Hsa-miR-155-5p Up-Regulation in Breast Cancer and Its Relevance for Treatment With Poly[ADP-Ribose] Polymerase 1 (PARP-1) Inhibitors. Frontiers in oncology 10, 1415 (2020). 10.3389/fonc.2020.01415

7 Hsu, C. H. et al. miR-29a-3p/THBS2 Axis Regulates PAH-Induced Cardiac Fibrosis. International journal of molecular sciences 22 (2021). 10.3390/ijms221910574

8 Li, L., He, X., Liu, M., Yun, L. & Cong, B. Diagnostic value of cardiac miR-126-5p, miR-134-5p, and miR-499a-5p in coronary artery disease-induced sudden cardiac death. Front Cardiovasc Med 9, 944317 (2022). 10.3389/fcvm.2022.944317

9 Beg, M. S. et al. Phase I study of MRX34, a liposomal miR-34a mimic, administered twice weekly in patients with advanced solid tumors. Invest New Drugs 35, 180–188 (2017). 10.1007/s10637-016-0407-y

10 Diener, C., Keller, A. & Meese, E. Emerging concepts of miRNA therapeutics: from cells to clinic. Trends Genet 38, 613–626 (2022). 10.1016/j.tig.2022.02.006

11 Viteri, S. & Rosell, R. An innovative mesothelioma treatment based on miR-16 mimic loaded EGFR targeted minicells (TargomiRs). Transl Lung Cancer Res 7, S1–S4 (2018). 10.21037/tlcr.2017.12.01

12 Grimm, D. et al. Fatality in mice due to oversaturation of cellular microRNA/short hairpin RNA pathways. Nature 441, 537–541 (2006). 10.1038/nature04791

13 Segal, M. & Slack, F. J. Challenges identifying efficacious miRNA therapeutics for cancer. Expert Opin Drug Discov 15, 987–992 (2020). 10.1080/17460441.2020.1765770

14 Hong, D. S. et al. Phase 1 study of MRX34, a liposomal miR-34a mimic, in patients with advanced solid tumours. British journal of cancer 122, 1630–1637 (2020). 10.1038/s41416-020-0802-1

15 Liu, Y. P. & Berkhout, B. miRNA cassettes in viral vectors: problems and solutions. Biochim Biophys Acta 1809, 732–745 (2011). 10.1016/j.bbagrm.2011.05.014

16 Good, C. R. et al. An NK-like CAR T cell transition in CAR T cell dysfunction. Cell 184, 6081–6100 e6026 (2021). 10.1016/j.cell.2021.11.016

17 Roman Azcona, M. S. et al. Sustained and specific multiplexed immune checkpoint modulation in CAR T cells induced by targeted epigenome editing. Molecular therapy. Nucleic acids 36, 102618 (2025). 10.1016/j.omtn.2025.102618

18 Locke, F. L. et al. Impact of tumor microenvironment on efficacy of anti-CD19 CAR T cell therapy or chemotherapy and transplant in large B cell lymphoma. Nat Med 30, 507–518 (2024). 10.1038/s41591-023-02754-1

19 Yan, Z. X. et al. Clinical Efficacy and Tumor Microenvironment Influence in a Dose-Escalation Study of Anti-CD19 Chimeric Antigen Receptor T Cells in Refractory B-Cell Non-Hodgkin’s Lymphoma. Clinical cancer research : an official journal of the American Association for Cancer Research 25, 6995–7003 (2019). 10.1158/1078-0432.CCR-19-0101

20 Liu, X. et al. A novel dominant-negative PD-1 armored anti-CD19 CAR T cell is safe and effective against refractory/relapsed B cell lymphoma. Transl Oncol 14, 101085 (2021). 10.1016/j.tranon.2021.101085

21 John, L. B. et al. Anti-PD-1 antibody therapy potently enhances the eradication of established tumors by gene-modified T cells. Clinical cancer research : an official journal of the American Association for Cancer Research 19, 5636–5646 (2013). 10.1158/1078-0432.CCR-13-0458

22 Dai, X. et al. Massively parallel knock-in engineering of human T cells. Nat Biotechnol 41, 1239–1255 (2023). 10.1038/s41587-022-01639-x

23 Shao, L., Zheng, Y., Somerville, R. P., Stroncek, D. F. & Jin, P. New insights on potency assays from recent advances and discoveries in CAR T-cell therapy. Frontiers in immunology 16, 1597888 (2025). 10.3389/fimmu.2025.1597888

24 Guo, L., Zhao, Y., Zhang, H., Yang, S. & Chen, F. Integrated evolutionary analysis of human miRNA gene clusters and families implicates evolutionary relationships. Gene 534, 24–32 (2014). 10.1016/j.gene.2013.10.037

25 Wang, Y., Luo, J., Zhang, H. & Lu, J. microRNAs in the Same Clusters Evolve to Coordinately Regulate Functionally Related Genes. Molecular biology and evolution 33, 2232–2247 (2016). 10.1093/molbev/msw089

26 Jiang, S. et al. Molecular dissection of the miR-17-92 cluster’s critical dual roles in promoting Th1 responses and preventing inducible Treg differentiation. Blood 118, 5487–5497 (2011). 10.1182/blood-2011-05-355644

27 Li, M. et al. Exosomal miR-92b-3p Promotes Chemoresistance of Small Cell Lung Cancer Through the PTEN/AKT Pathway. Front Cell Dev Biol 9, 661602 (2021). 10.3389/fcell.2021.661602

28 Gagnon, J. D. et al. miR-15/16 Restrain Memory T Cell Differentiation, Cell Cycle, and Survival. Cell reports 28, 2169–2181 e2164 (2019). 10.1016/j.celrep.2019.07.064

29 Dibas, A. et al. Cell-Based Models of ‘Cytokine Release Syndrome’ Endorse CD40L and Granulocyte-Macrophage Colony-Stimulating Factor Knockout in Chimeric Antigen Receptor T Cells as Mitigation Strategy. Cells 12 (2023). 10.3390/cells12212581

30 Tian, Y. et al. CCR5 and IL-12 co-expression in CAR T cells improves antitumor efficacy by reprogramming tumor microenvironment in solid tumors. Cancer immunology, immunotherapy : CII 74, 55 (2025). 10.1007/s00262-024-03909-w

31 Gong, L. et al. Cancer immunology data engine reveals secreted AOAH as a potential immunotherapy. Cell 188, 5062–5080 e5032 (2025). 10.1016/j.cell.2025.07.004

32 Tan, W. et al. Novel immune-related genes in the tumor microenvironment with prognostic value in breast cancer. BMC cancer 21, 126 (2021). 10.1186/s12885-021-07837-1

33 Kagoya, Y. et al. A novel chimeric antigen receptor containing a JAK-STAT signaling domain mediates superior antitumor effects. Nat Med 24, 352–359 (2018). 10.1038/nm.4478

34 Laugel, B. et al. The multiple roles of the CD8 coreceptor in T cell biology: opportunities for the selective modulation of self-reactive cytotoxic T cells. J Leukoc Biol 90, 1089–1099 (2011). 10.1189/jlb.0611316

35 Li, C. et al. Integrin CD103 expression in naive CD8(+) T cells promotes cytokine-driven acquisition of memory phenotype and effector function. Immunity 58, 2734–2752 e2739 (2025). 10.1016/j.immuni.2025.08.014

36 Lingel, H. et al. SLAMF7 (CD319) on activated CD8(+) T cells transduces environmental cues to initiate cytotoxic effector cell responses. Cell Death Differ 32, 561–572 (2025). 10.1038/s41418-024-01399-y

37 Menasche, G. et al. A newly identified isoform of Slp2a associates with Rab27a in cytotoxic T cells and participates to cytotoxic granule secretion. Blood 112, 5052–5062 (2008). 10.1182/blood-2008-02-141069

38 Saini, R. V. et al. Granulysin delivered by cytotoxic cells damages endoplasmic reticulum and activates caspase-7 in target cells. Journal of immunology 186, 3497–3504 (2011). 10.4049/jimmunol.1003409

39 Dong, H. et al. A Dual Role for NKG7 in T-cell Cytotoxicity and Longevity. Cancer immunology research 13, 1510–1515 (2025). 10.1158/2326-6066.CIR-25-0384

40 Hendriks, J. et al. CD27 is required for generation and long-term maintenance of T cell immunity. Nature immunology 1, 433–440 (2000). 10.1038/80877

41 Oba, T., Long, M. D., Ito, K. I. & Ito, F. Clinical and immunological relevance of SLAMF6 expression in the tumor microenvironment of breast cancer and melanoma. Scientific reports 14, 2394 (2024). 10.1038/s41598-023-50062-y

42 Xiong, Y. et al. c-Kit signaling potentiates CAR T cell efficacy in solid tumors by CD28- and IL-2-independent co-stimulation. Nat Cancer 4, 1001–1015 (2023). 10.1038/s43018-023-00573-4

43 Hao, Z. et al. JAML promotes the antitumor role of tumor-resident CD8(+) T cells by facilitating their innate-like function in human lung cancer. Cancer letters 590, 216839 (2024). 10.1016/j.canlet.2024.216839

44 Pfistershammer, K. et al. CD63 as an activation-linked T cell costimulatory element. Journal of immunology 173, 6000–6008 (2004). 10.4049/jimmunol.173.10.6000

45 Woo, J. S. et al. CRACR2A-Mediated TCR Signaling Promotes Local Effector Th1 and Th17 Responses. Journal of immunology 201, 1174–1185 (2018). 10.4049/jimmunol.1800659

46 Di Simone, M. et al. Tumor-infiltrating gammadelta T cells as targets of immune checkpoint blockade in melanoma. J Leukoc Biol 115, 760–770 (2024). 10.1093/jleuko/qiae023

47 Jung, I. Y. et al. Type I Interferon Signaling via the EGR2 Transcriptional Regulator Potentiates CAR T Cell-Intrinsic Dysfunction. Cancer Discov 13, 1636–1655 (2023). 10.1158/2159-8290.CD-22-1175

48 Sun, S., Zhi, Z., Su, Y., Sun, J. & Li, Q. A CD8(+) T cell-associated immune gene panel for prediction of the prognosis and immunotherapeutic effect of melanoma. Frontiers in immunology 13, 1039565 (2022). 10.3389/fimmu.2022.1039565

49 Chen, G. M. et al. Integrative Bulk and Single-Cell Profiling of Premanufacture T-cell Populations Reveals Factors Mediating Long-Term Persistence of CAR T-cell Therapy. Cancer Discov 11, 2186–2199 (2021). 10.1158/2159-8290.CD-20-1677

50 Dar, A. A. et al. Extracellular 2’5’-oligoadenylate synthetase 2 mediates T-cell receptor CD3-zeta chain down-regulation via caspase-3 activation in oral cancer. Immunology 147, 251–264 (2016). 10.1111/imm.12560

51 DeDiego, M. L., Nogales, A., Martinez-Sobrido, L. & Topham, D. J. Interferon-Induced Protein 44 Interacts with Cellular FK506-Binding Protein 5, Negatively Regulates Host Antiviral Responses, and Supports Virus Replication. mBio 10 (2019). 10.1128/mBio.01839-19

52 Zeng, Y. et al. IFI44L as a novel epigenetic silencing tumor suppressor promotes apoptosis through JAK/STAT1 pathway during lung carcinogenesis. Environmental pollution 319, 120943 (2023). 10.1016/j.envpol.2022.120943

53 Cagnoni, A. J. et al. Galectin-1 fosters an immunosuppressive microenvironment in colorectal cancer by reprogramming CD8(+) regulatory T cells. Proc Natl Acad Sci U S A 118 (2021). 10.1073/pnas.2102950118

54 Schlabach, M. R. et al. Rational design of a SOCS1-edited tumor-infiltrating lymphocyte therapy using CRISPR/Cas9 screens. J Clin Invest 133 (2023). 10.1172/JCI163096

55 Zhu, C. et al. The Tim-3 ligand galectin-9 negatively regulates T helper type 1 immunity. Nature immunology 6, 1245–1252 (2005). 10.1038/ni1271

56 Marti i Lindez, A. A. et al. Mitochondrial arginase-2 is a cell-autonomous regulator of CD8+ T cell function and antitumor efficacy. JCI insight 4 (2019). 10.1172/jci.insight.132975

57 Lacher, S. B. et al. PGE(2) limits effector expansion of tumour-infiltrating stem-like CD8(+) T cells. Nature 629, 417–425 (2024). 10.1038/s41586-024-07254-x

58 Chen, Q. et al. The multifaceted role of Sestrin 3 (SESN3) in oxidative stress, inflammation and tumorigenesis. Biochim Biophys Acta Mol Cell Res 1872, 119938 (2025). 10.1016/j.bbamcr.2025.119938

59 Veliz, K. et al. Deletion of CD38 enhances CD19 chimeric antigen receptor T cell function. Mol Ther Oncol 32, 200819 (2024). 10.1016/j.omton.2024.200819

60 Bishop, A. L. et al. Double-tap gene drive uses iterative genome targeting to help overcome resistance alleles. Nat Commun 13, 2595 (2022). 10.1038/s41467-022-29868-3

61 Moller, L. et al. Recursive Editing improves homology-directed repair through retargeting of undesired outcomes. Nat Commun 13, 4550 (2022). 10.1038/s41467-022-31944-7

62 Anzalone, A. V. et al. Search-and-replace genome editing without double-strand breaks or donor DNA. Nature 576, 149–157 (2019). 10.1038/s41586-019-1711-4

63 Witte, I. P. et al. Programmable gene insertion in human cells with a laboratory-evolved CRISPR-associated transposase. Science 388, eadt5199 (2025). 10.1126/science.adt5199

64 Yarnall, M. T. N. et al. Drag-and-drop genome insertion of large sequences without double-strand DNA cleavage using CRISPR-directed integrases. Nat Biotechnol 41, 500–512 (2023). 10.1038/s41587-022-01527-4

65 Suzuki, K. et al. In vivo genome editing via CRISPR/Cas9 mediated homology-independent targeted integration. Nature 540, 144–149 (2016). 10.1038/nature20565

66 Haider, S. & Mussolino, C. Fine-Tuning Homology-Directed Repair (HDR) for Precision Genome Editing: Current Strategies and Future Directions. International journal of molecular sciences 26 (2025). 10.3390/ijms26094067

67 Carusillo, A. et al. A novel Cas9 fusion protein promotes targeted genome editing with reduced mutational burden in primary human cells. Nucleic Acids Res 51, 4660–4673 (2023). 10.1093/nar/gkad255

68 Livak, K. J. & Schmittgen, T. D. Analysis of relative gene expression data using real-time quantitative PCR and the 2(-Delta Delta C(T)) Method. Methods 25, 402–408 (2001). 10.1006/meth.2001.1262

69 Khamina, K. et al. A MicroRNA Next-Generation-Sequencing Discovery Assay (miND) for Genome-Scale Analysis and Absolute Quantitation of Circulating MicroRNA Biomarkers. International journal of molecular sciences 23 (2022). 10.3390/ijms23031226

70 Diendorfer, A., Khamina, K., Pultar, M. & Hackl, M. miND (miRNA NGS Discovery pipeline): a small RNA-seq analysis pipeline and report generator for microRNA biomarker discovery studies. F1000Research 11 (2022). 10.12688/f1000research.94159.1

71 Metsalu, T. & Vilo, J. ClustVis: a web tool for visualizing clustering of multivariate data using Principal Component Analysis and heatmap. Nucleic Acids Res 43, W566–570 (2015). 10.1093/nar/gkv468

72 Ru, Y. et al. The multiMiR R package and database: integration of microRNA-target interactions along with their disease and drug associations. Nucleic Acids Res 42, e133 (2014). 10.1093/nar/gku631

73 Maragkakis, M. et al. DIANA-microT web server: elucidating microRNA functions through target prediction. Nucleic Acids Res 37, W273–276 (2009). 10.1093/nar/gkp292

74 Gaidatzis, D., van Nimwegen, E., Hausser, J. & Zavolan, M. Inference of miRNA targets using evolutionary conservation and pathway analysis. BMC Bioinformatics 8, 69 (2007). 10.1186/1471-2105-8-69

75 Griffiths-Jones, S., Saini, H. K., van Dongen, S. & Enright, A. J. miRBase: tools for microRNA genomics. Nucleic Acids Res 36, D154–158 (2008). 10.1093/nar/gkm952

76 Kertesz, M., Iovino, N., Unnerstall, U., Gaul, U. & Segal, E. The role of site accessibility in microRNA target recognition. Nat Genet 39, 1278–1284 (2007). 10.1038/ng2135

77 Betel, D., Wilson, M., Gabow, A., Marks, D. S. & Sander, C. The microRNA.org resource: targets and expression. Nucleic Acids Res 36, D149–153 (2008). 10.1093/nar/gkm995

78 Anders, G. et al. doRiNA: a database of RNA interactions in post-transcriptional regulation. Nucleic Acids Res 40, D180–186 (2012). 10.1093/nar/gkr1007

79 Friedman, R. C., Farh, K. K., Burge, C. B. & Bartel, D. P. Most mammalian mRNAs are conserved targets of microRNAs. Genome Res 19, 92–105 (2009). 10.1101/gr.082701.108

80 Wang, X. miRDB: a microRNA target prediction and functional annotation database with a wiki interface. Rna 14, 1012–1017 (2008). 10.1261/rna.965408

81 Hsu, S. D. et al. miRTarBase: a database curates experimentally validated microRNA-target interactions. Nucleic Acids Res 39, D163–169 (2011). 10.1093/nar/gkq1107

82 Xiao, F. et al. miRecords: an integrated resource for microRNA-target interactions. Nucleic Acids Res 37, D105–110 (2009). 10.1093/nar/gkn851

83 Vergoulis, T. et al. TarBase 6.0: capturing the exponential growth of miRNA targets with experimental support. Nucleic Acids Res 40, D222–229 (2012). 10.1093/nar/gkr1161

84 Alexa, A. & Rahnenfuhrer, J. topGO: Enrichment analysis for Gene Ontology. R package version 2.28. 0. Cranio 2, 11 (2016).

85 Gagnon, J. D. & Ansel, K. M. MicroRNA regulation of CD8(+) T cell responses. Noncoding RNA Investig 3 (2019). 10.21037/ncri.2019.07.02

86 Wheeler, B. D. et al. The lncRNA Malat1 inhibits miR-15/16 to enhance cytotoxic T cell activation and memory cell formation. eLife 12 (2023). 10.7554/eLife.87900

87 Zhao, F. et al. miR-92b-3p Regulates Cell Cycle and Apoptosis by Targeting CDKN1C, Thereby Affecting the Sensitivity of Colorectal Cancer Cells to Chemotherapeutic Drugs. Cancers (Basel) 13 (2021). 10.3390/cancers13133323

88 Sasaki, K. et al. miR-17-92 expression in differentiated T cells - implications for cancer immunotherapy. Journal of translational medicine 8, 17 (2010). 10.1186/1479-5876-8-17

89 Steiner, D. F. et al. MicroRNA-29 regulates T-box transcription factors and interferon-gamma production in helper T cells. Immunity 35, 169–181 (2011). 10.1016/j.immuni.2011.07.009

