## Supplementary material for "*Castling*, a novel therapeutic concept for rewiring pathological gene-expression networks, enabled by the TRIPLE technology": Antony_et_al_SuppleentaryData

| | |
|---|---|
| Supplementary Figure 1 | Kinetic and endpoint analysis of CAR T cells during the chronic antigen stimulation |
| Supplementary Figure 2 | Expression level dynamics of miRNA selected for *castling* |
| Supplementary Figure 3 | Optimization of experimental conditions for deleting the miR-15/16 clusters |
| Supplementary Figure 4 | Effect of TRIPLE genome editing procedure on DNA repair |
| Supplementary Figure 5 | *Castling* affects expression levels of all isoforms (isomiRs) of the target miRNAs. |
| Supplementary Figure 6 | Expression level dynamics of the *castled* miRNAs in CAR T cells during the chronic antigen stimulation |
| Supplementary Figure 7 | Functional analysis of *castled* CAR T cells |
| Supplementary Figure 8 | Principal component analysis (PCA) of transcriptomic changes during chronic antigen stimulation |
| Supplementary Table 1 | Top candidate miRNAs, enriched for clustered regulators |
| Supplementary Table 2 | Spacer sequences of the guide RNAs (gRNAs) used in this study |
| Supplementary Table 3 | Oligonucleotides used in this study and their purpose |
| Supplementary Table 4 | List of genes upregulated (FDR < 0.05) upon miR-15/16 cluster KO |
| Supplementary Table 5 | Supplementary Table 5. List of genes downregulated (FDR < 0.05) upon miR-17~92 cluster KI |
| Supplementary Table 6 | List of significantly deregulated genes upon castling of miR-17~92 cluster into the miR-15a/16-1 locus |
| Supplementary Table 7 | Sequence of the repair templates used in this study |
| Supplementary Table 8 | Landmark sequences used for the long read amplicon sequencing analysis |

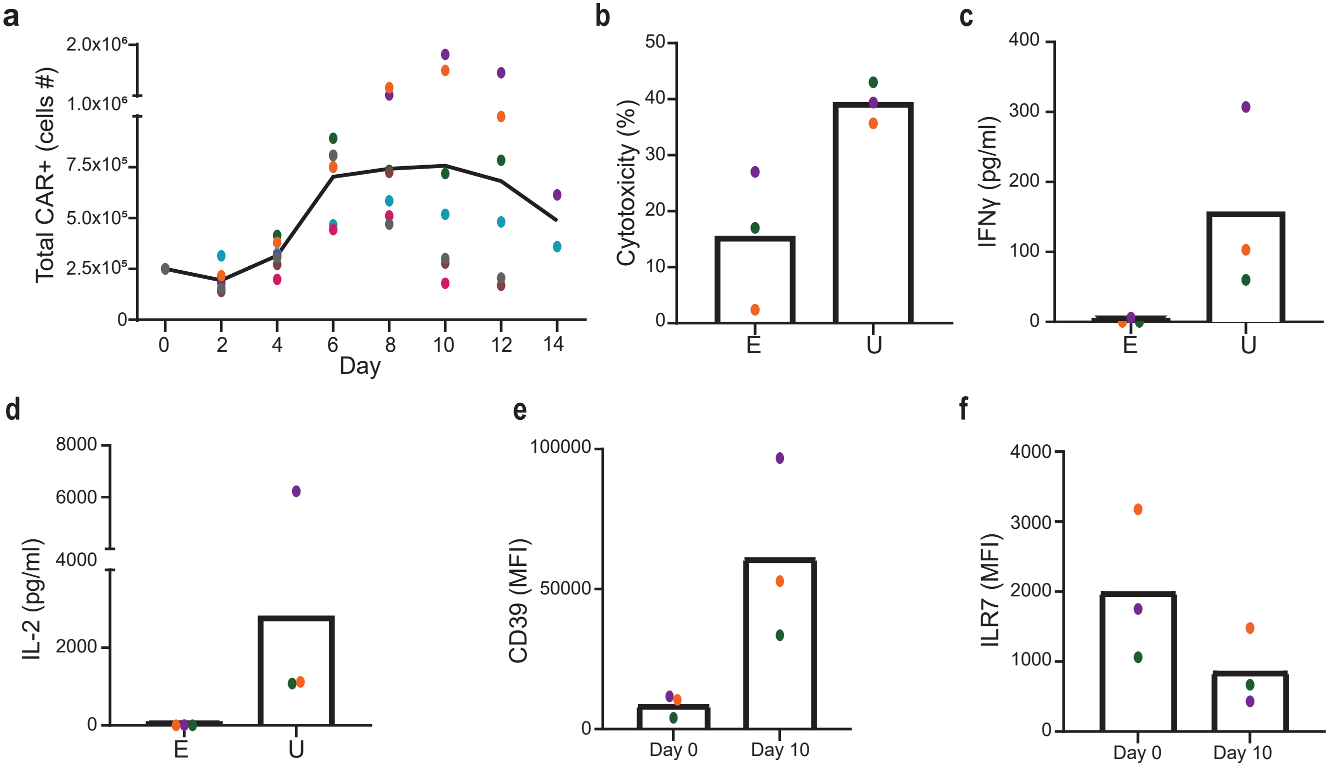

Antony et al. Supplementary Figure 1

**Supplementary Fig. 1. Kinetic and endpoint analysis of CAR T cells during the chronic antigen stimulation.** (**a**) Proliferation of CAR T cells over time. The graph indicates the total absolute number of CAR+ T cells at the indicated time points. Each dot represents the number of CAR T cells derived from a different T cell donor.  Donor origin is color coded and the colors are consistently used across all other panels in this figure to enable direct comparison. This panel summarizes proliferation data obtained from multiple T cell donors, whereas subsequent panels display detailed molecular and functional analyses from a subset of three donors that underwent full characterization (**b-f**).  Functional comparison between end-point samples at day 12/14 (E) and untreated CAR T cells (U) for three independent T cell donors. The bar graphs indicate the cytotoxicity capacity (**b**) and the amount of secreted IFNg (**c**) or IL-2 (**d**) cytokines (pg/ml). (**e, f**) The graph indicates the expression levels of exhaustion (CD39, **e**) and T cell memory (ILR7, **f**) markers, respectively, at the indicated time point within the assay for the T cell donors depicted measured via flow cytometry. MFI: mean fluorescence intensity.

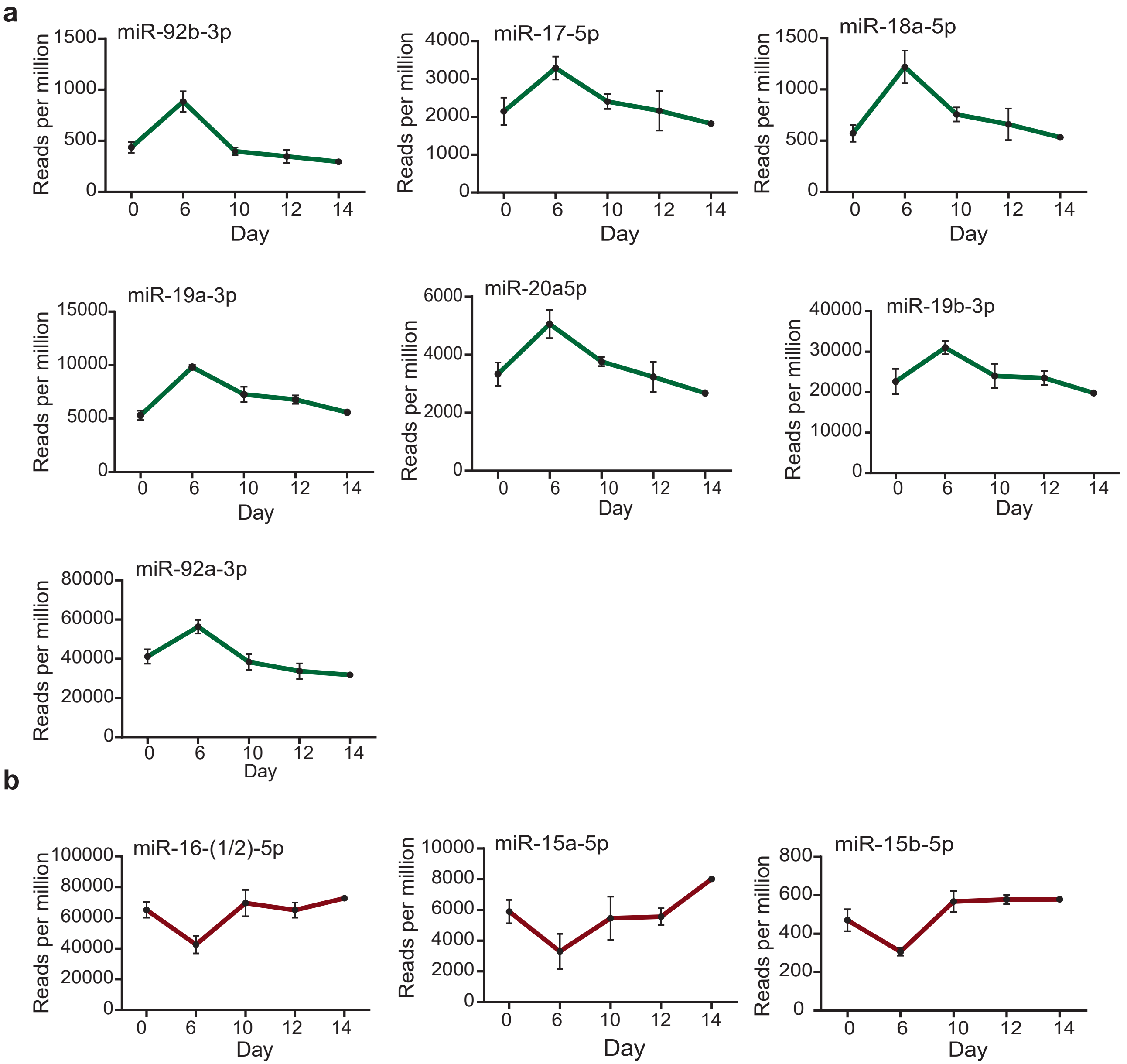

Antony et al. Supplementary Figure 2

**Supplementary Fig. 2. Expression level dynamics of miRNA selected for castling.** (**a, b**) The graphs show the small RNA sequencing results (reads per million, RPM) at the indicated time points, for selected miRNAs from either profile C (**a**) or D (**b**) as shown in Figure 1c. The miRNAs identities are indicated on top of each graph.

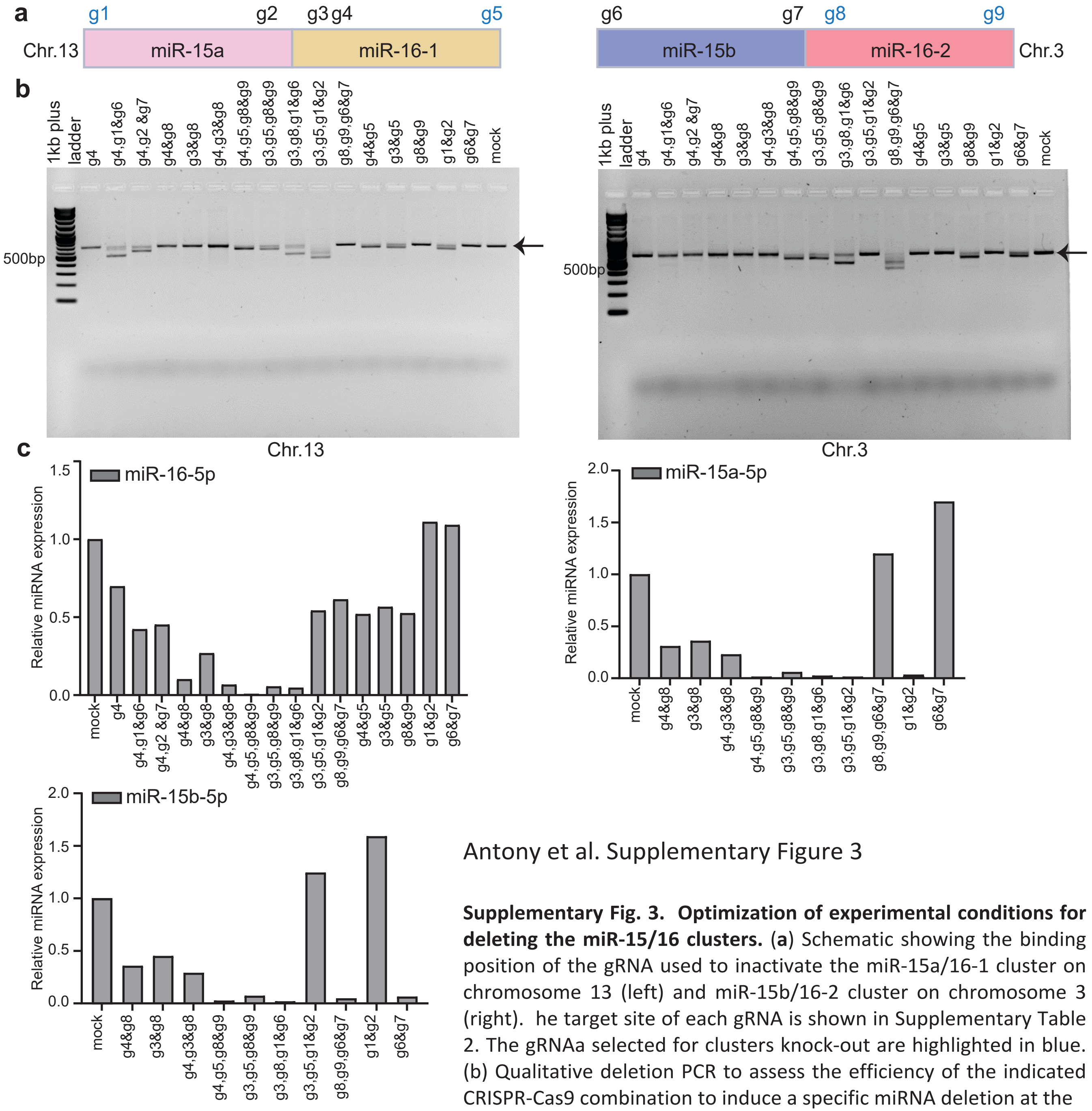

Antony et al. Supplementary Figure 3

**Supplementary Fig. 3. Optimization of experimental conditions for deleting the miR-15/16 clusters.** (a) Schematic showing the binding position of the gRNA used to inactivate the miR-15a/16-1 cluster on chromosome 13 (left) and miR-15b/16-2 cluster on chromosome 3 (right). he target site of each gRNA is shown in Supplementary Table 2. The gRNAa selected for clusters knock-out are highlighted in blue. (b) Qualitative deletion PCR to assess the efficiency of the indicated CRISPR-Cas9 combination to induce a specific miRNA deletion at the indicated locus (miR-15a/16-1 on chromosome 13, left) and (miR-15b/16-2 on chromosome 3, right). The arrows indicate the amplification band resulting from the intact alleles (~660bp). All bands below are indicative of effective miRNA deletion. (c) Bar graphs showing the expression levels of the indicated miRNA measured via real-time quantitative RT-PCR. gRNAs used are indicated below. Note that expression of miR-16-1 and miR-16-2 were measured using the same TaqMan probe that could not discriminate between the two and, therefore, their cumulative expression level is indicated.

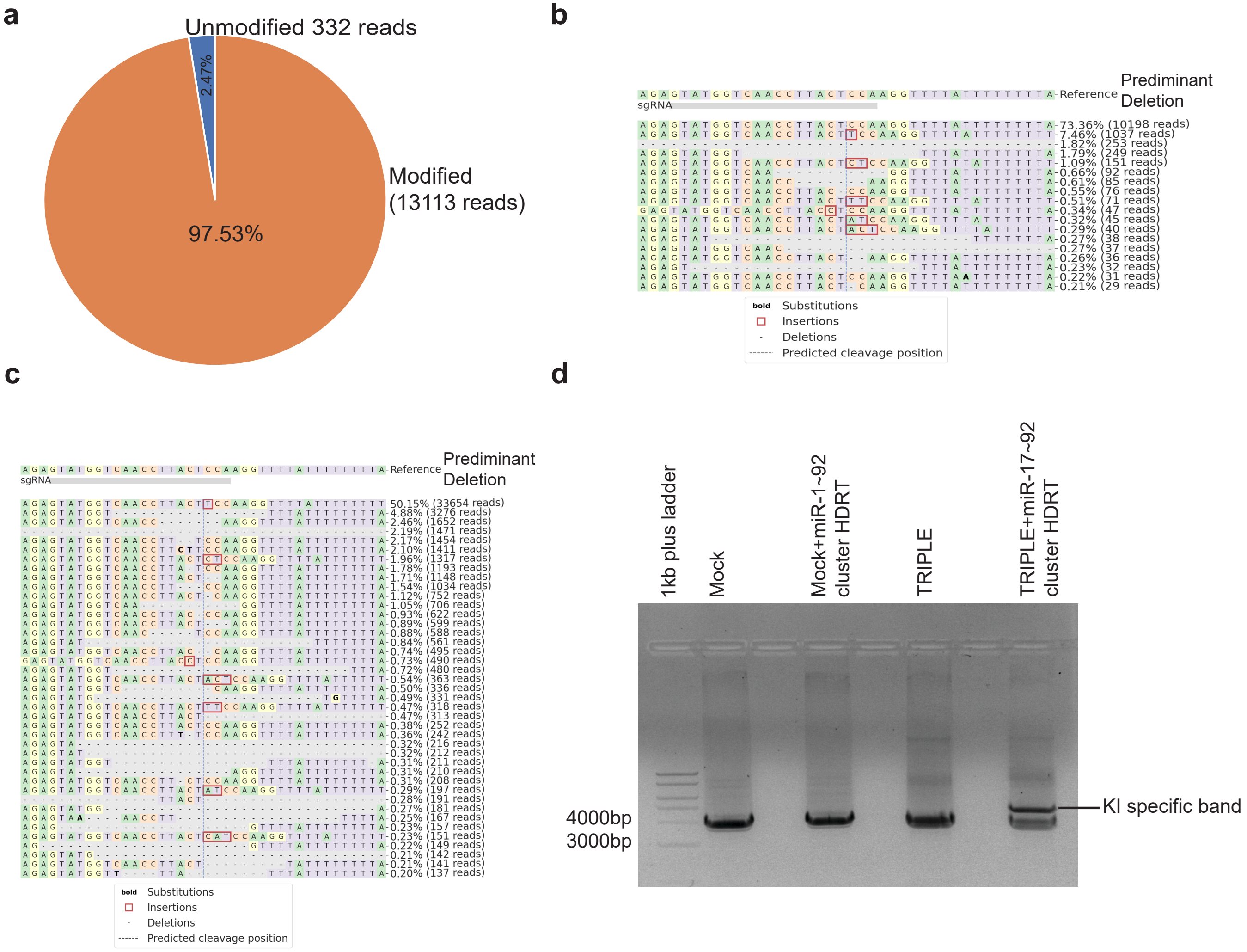

Antony et al. Supplementary Figure 4

**Supplementary Fig. 4. Effect of TRIPLE genome editing procedure on DNA repair.** (**a**) The pie chart shows the frequency of alleles harboring indel mutations resulting from the activation of the non-homologous end-joining (NHEJ) DNA repair pathway and measured via small amplicon sequencing at the target locus (miR-15a/16-1 cluster on chromosome 13). Unmodified reads matching the wild-type DNA sequence (blue) and reads harboring mutations (modified, orange) are indicated. (**b**) Frequency of the deleted allele at the miR-15a/16-1 locus on chromosome 13, resulting from the activity of the g1 and g5 RNPs (two RNP procedure), determined using Illumina paired-end amplicon sequencing and analyzed using the CRISPReesso2 pipeline. The given reference sequence indicates the predominant deletion allele, which represents about 73% of the total reads. (**c**) Frequency of the predominant deletion allele as in (b) after TRIPLE. The addition of the third RNP comprising a gRNA targeting the site of predominant deletion (indicated as sgRNA within the image) leads to its disappearance and results in a novel predominant +1nt insertion, typical of CRISPR-Cas9 editing. (**d**) Qualitative PCR analysis indicated the presence of a band specific for miR-17~92 cluster integration (~4732bp) upon castling at the miR-15a/16-1 locus on chromosome 13..

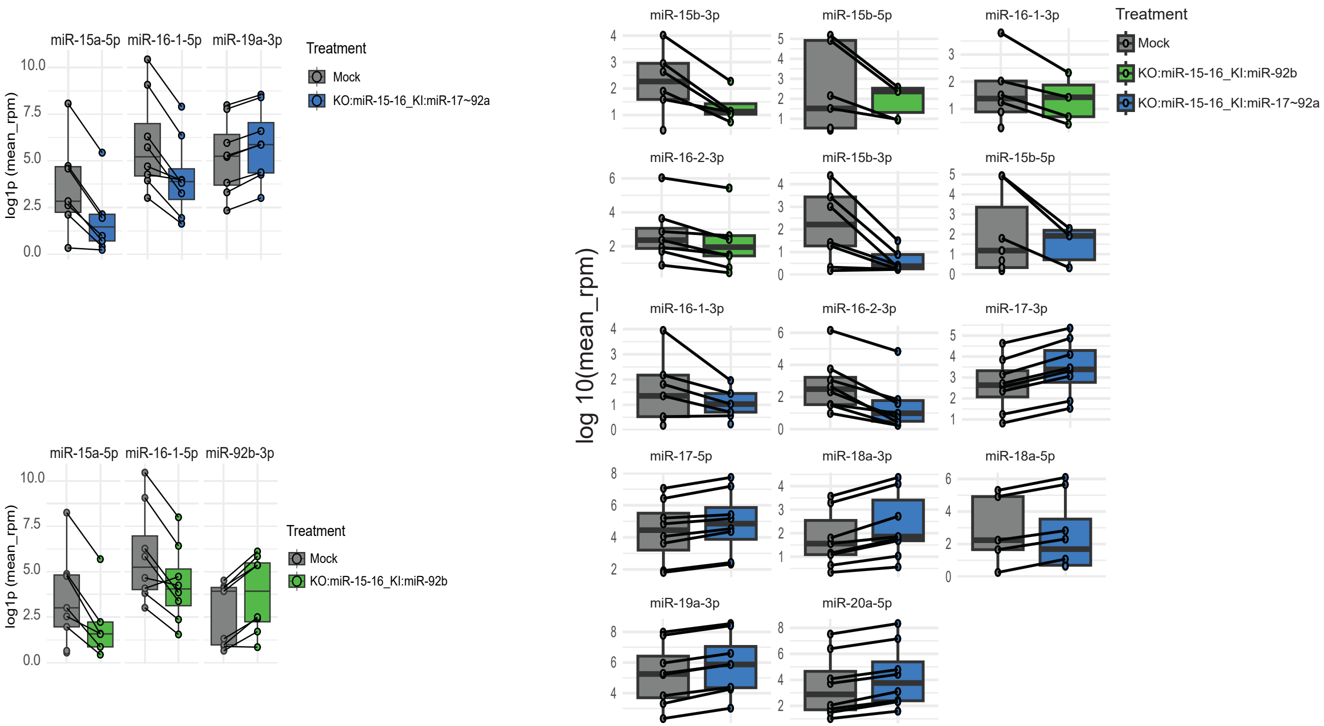

Antony et al. Supplementary Figure 5

**Supplementary Fig. 5. *Castling* affects expression levels of all isoforms (isomiRs) of the target miRNAs.** The box plots (left panels) show the expression levels of the castled miRNA measured via small RNA sequencing (log1p(mean RPM)) in the indicated samples at the pre-antigen stimulation time point. The expression levels of individual detectable isoforms of mature microRNAs (right panel) in the respective *castled* samples are shown as small circles. In both panels the castling of the miR-17~92 cluster or miR-92b into the miR-15a/16-1 cluster on chromosome 13 are indicated in blue or green, respectively, while the mock samples are indicated in grey.

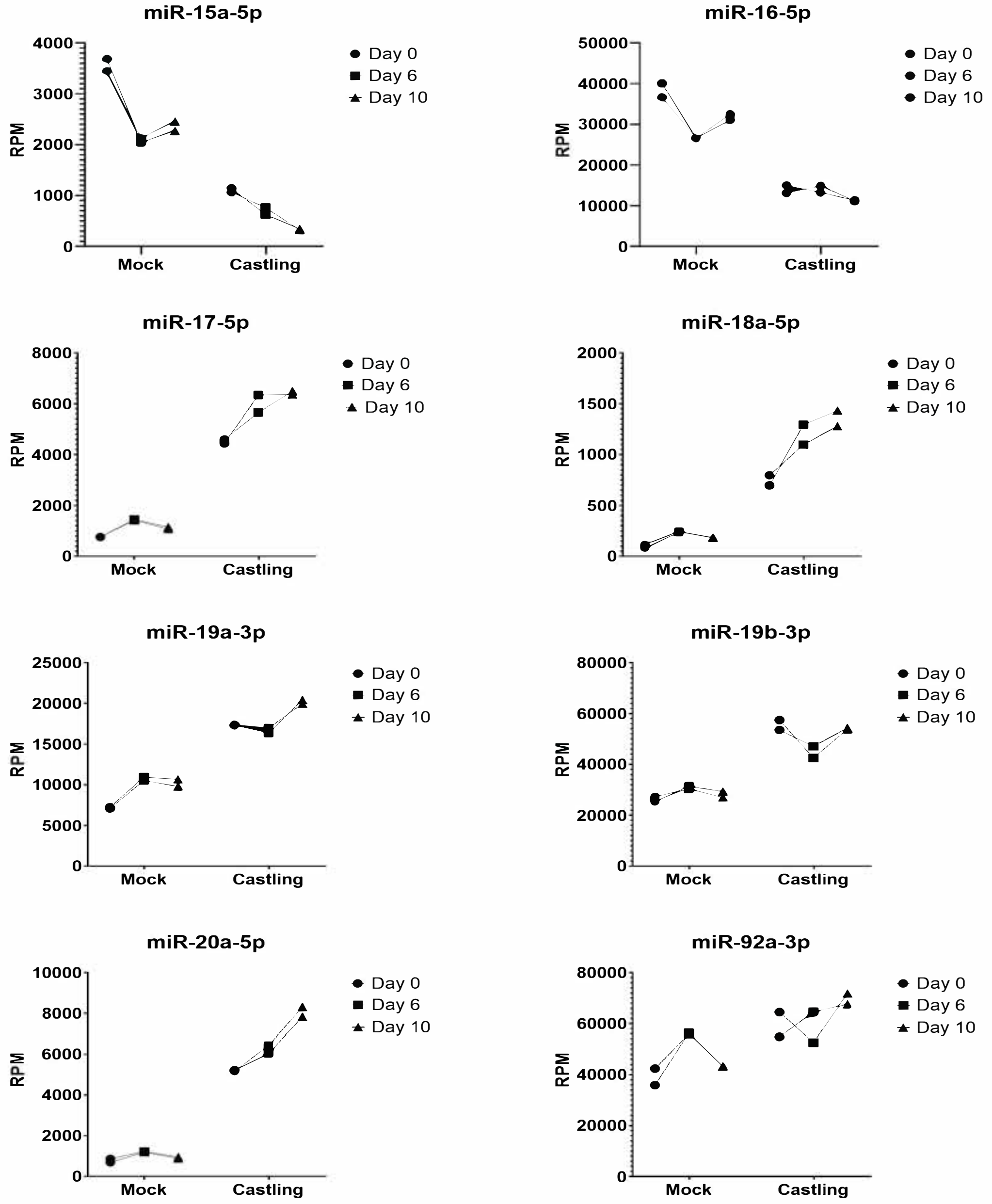

Antony et al. Supplementary Figure 6

**Supplementary Fig. 6. Expression level dynamics of the castled miRNAs in CAR T cells during the chronic antigen stimulation.** The graphs display the expression levels of the *castled* miRNA measured via small RNA sequencing (reads per million, RPM), at the indicated time point of the chronic antigen stimulation. *Castling* = CAR T cells edited with the indicated *castling* combination; mock = control non-edited CAR-T cells. The specific miRNA measured is indicated on top of each graph.

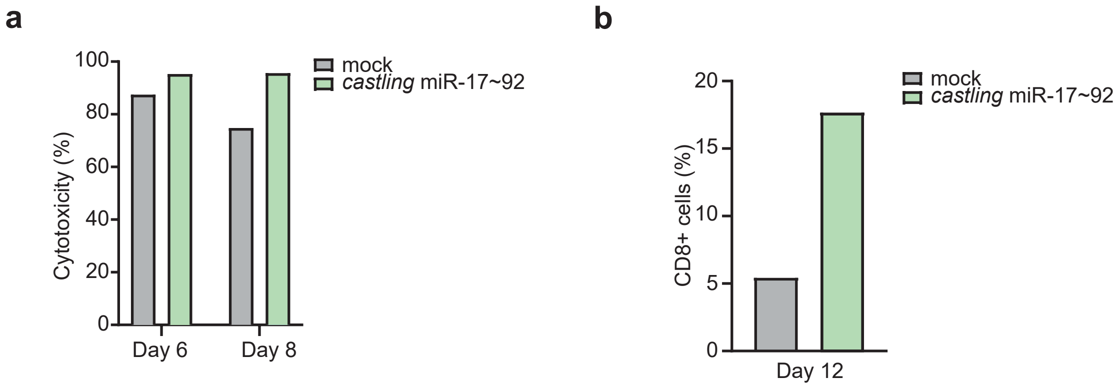

Antony et al. Supplementary Figure 7

**Supplementary Fig. 7. Functional analysis of castled CAR T cells.** The bar graphs indicate the cytotoxic activity (**a**) and the frequency of CD8+ T cells (**b**) in CAR T cell populations upon *castling* of miR-17~92 cluster into miR-15a/16-1 locus on chromosome 13, as compared to mock controls at each indicated time point.

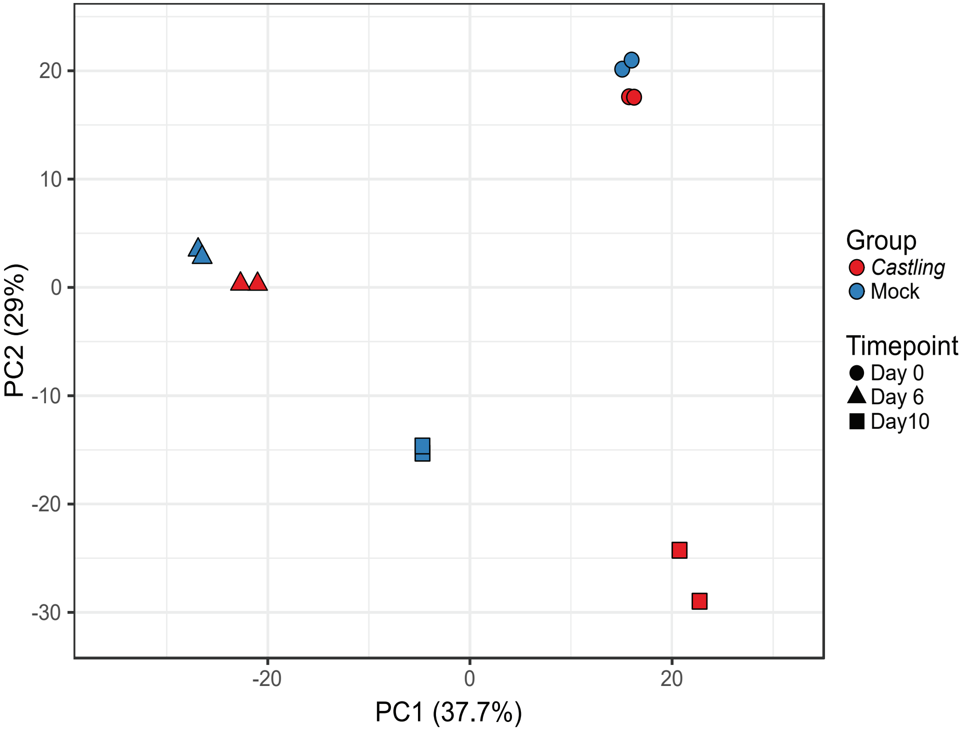

Antony et al. Supplementary Figure 8

**Supplementary Fig. 8. Principal component analysis (PCA) of transcriptomic changes during chronic antigen stimulation.** PCA plot based on ln-transformed RPM values, showing separation of *castled* and mock control CAR T cells across the indicated time points during chronic antigen stimulation. The analysis was performed using highly variable mRNAs (n = 1’054), corresponding to the top 10% ranked by coefficient of variation (CV).

**Supplementary Table 1. Top candidate miRNAs, enriched for clustered regulators**

| miRNA ID | Number and ID of hairpin precursors | Clustered | Cluster name | Cluster members |
|---|---|---|---|---|
| miR-23a | 1 | Yes | miR-23a~27a~24-2 | miR-23a, miR-27a and miR24-2 |
| miR-27a | 1 | Yes | miR-23a~27a~24-2 | miR-23a, miR-27a and miR24-2 |
| miR-92a | 2; miR-92a-1, miR-92a-2 | Yes | miR-17~92 | miR-17, miR-18a, miR-19a, miR-20a, miR-19b-1 and miR-92a-1 |
| miR-92b | 1 | No | - | - |
| miR-19b | 2; miR-19b-1, miR-19b-2 | Yes | miR-17~92 | miR-17, miR-18a, miR-19a, miR-20a, miR-19b-1 and miR-92a-1 |
| miR-18a | 1 | Yes | miR-17~92 | miR-17, miR-18a, miR-19a, miR-20a, miR-19b-1 and miR-92a-1 |
| miR-17 | 1 | Yes | miR-17~92 | miR-17, miR-18a, miR-19a, miR-20a, miR-19b-1 and miR-92a-1 |
| miR-20a | 1 | Yes | miR-17~92 | miR-17, miR-18a, miR-19a, miR-20a, miR-19b-1 and miR-92a-1 |
| miR-18a | 1 | Yes | miR-17~92 | miR-17, miR-18a, miR-19a, miR-20a, miR-19b-1 and miR-92a-1 |
| miR-155 | 1 | No | - | - |
| miR-19a | 1 | Yes | miR-17~92 | miR-17, miR-18a, miR-19a, miR-20a, miR-19b-1 and miR-92a-1 |
| miR-20b | 1 | Yes | miR-106a~363 | miR-106a, miR-18b, miR-20b, miR-19b-2, miR-92a-2, and miR-363 |
| miR-363 | 1 | Yes | miR-106a~363 | miR-106a, miR-18b, miR-20b, miR-19b-2, miR-92a-2, and miR-363 |
| miR-342 | 1 | Yes | miR-151b ~ 342 | miR-151b and miR-342 |
| miR-150 | 1 | No | - | - |
| miR-16-2 | 1 | Yes | miR-15b~16-2 | miR-15b and miR-16-2 |
| miR-29c | 1 | Yes | miR-29c~miR-29b-2 | miR-29c and miR-29b-2 |
| miR-142 | 1 | Yes | miR-142~ 4736 | miR-142 and miR-4736 |
| miR-181a | 2; miR-181a-1, miR-181a-2 | Yes | miR-181a-1 ~ 181b-1 | miR-181a-1 and miR-181b-1 |

| | | | | |
|---|---|---|---|---|
| miR-181b | 2; miR-181b-1, miR-181b-2 | Yes | miR-181a-1 ~ 181b-1 | miR-181a-1 and miR-181b-1 |
| miR-30b | 1 | Yes | miR-30b~ 30d | miR-30b and miR-30d |
| miR-140 | 1 | No | - | - |
| let-7b | 1 | Yes | let-7a~let-7b~miR-4763 | let-7a, let-7b, and miR-4763 |
| let-7g | 1 | No | - | - |
| miR-28 | 1 | No | - | - |
| miR-33a | 1 | No | - | - |
| let-7i | 1 | No | - | - |
| miR-140 | 1 | No | - | - |
| miR-22 | 1 | No | - | - |
| miR-30d | 1 | Yes | miR-30b~ 30d | miR-30b and miR-30d |
| miR-454 | 1 | No | - | - |
| miR-15a | 1 | Yes | miR-15a~16-1 | miR-15a and miR-16-1 |
| miR-16-1 | 1 | Yes | miR-15a~16-1 | miR-15a and miR-16-1 |
| let-7a | 3; let-7a-1, let-7a-2, let-7a-3 | Yes | let-7a-1~let-7d | let-7a-1, let-7f-1, and let-7d |
| miR-15b | 1 | Yes | miR-15b~16-2 | miR-15b and miR-16-2 |
| miR-30e | 1 | Yes | miR-30c-1~30e | miR-30c-1 and miR-30e |
| miR-26a | 2; miR-26a-1, miR-26a-2 | No | - | - |
| miR-26b | 1 | No | - | - |

**Supplementary Table 2. Spacer sequences of the guide RNAs (gRNAs) used in this study**

| gRNA ID | Sequence (5'--> 3') |
|---|---|
| g1 | tgtgctgctactttactcca |
| g2 | tgtgctgcctcaaaaataca |
| g3 | ccaatatttacgtgctgcta |
| g4 | ccttagcagcacgtaaatat |
| g5 | ttaactgtgctgctgaagta |
| g6 | agtactgtagcagcacatca |
| g7 | tttgctgctctagaaattta |
| g8 | atatttacgtgctgctagag |
| g9 | tgtgctgctttagtgtgaca |
| TRIPLE | gtatggtcaaccttactcca |

**Supplementary Table 3. Oligonucleotides used in this study and their purpose**

| Oligo ID | Sequence (5'--> 3') | Purpose |
|---|---|---|
| Lp 87 | aatgtatgtgatgagggggacc | miR-92b *castling* gene editing asssesment (Figure 2b; Supplementary Fig. 4c) |
| Lp 84 | agagacagggtttcaccacattg | |
| Lp 80 | gcatattacatcaatgttat | |
| Lp 81 | caagattatcaataatactg | |
| Lp 82 | gtatgtgatgagggggaccctg | miR-17~92 *castling* gene editing asssesment (Figure 2c) |
| Lp 84 | agagacagggtttcaccacattg | |
| Lp39 | gggcacagaatggacttcag | Qualitative PCR assessment to assess gRNA efficency and amplicon sequencing to design TRIPLE (Supplementary Fig.3b, 4a, 4b) |
| Lp15 | tcctctaatgctgcataagc | |
| Lp 41 | cagaacggcctgcagagataa | Qualitative PCR assessment to assess gRNA efficency (Supplementary Fig.3c) |
| Lp 42 | tgcttaggtaaatcaaacaccaagt | |

**Supplementary Table 4. List of genes upregulated (FDR < 0.05) upon miR-15/16 cluster KO**

| microRNA | Target Gene Symbol | ENSG | logFC_miRNA | logFC_mRNA |
|---|---|---|---|---|
| hsa-miR-15a-5p | *CCNE1* | ENSG00000105173 | -3,78 | 1,25 |
| hsa-miR-15a-5p | *ARL2* | ENSG00000213465 | -3,78 | 1,17 |
| hsa-miR-15a-5p | *KIF23* | ENSG00000137807 | -3,78 | 1,01 |
| hsa-miR-15a-5p | *CASK* | ENSG00000147044 | -3,78 | 0,92 |
| hsa-miR-15a-5p | *TSPAN5* | ENSG00000168785 | -3,78 | 0,92 |
| hsa-miR-15a-5p | *FNIP1* | ENSG00000217128 | -3,78 | 0,86 |
| hsa-miR-15a-5p | *ANAPC13* | ENSG00000129055 | -3,78 | 0,81 |
| hsa-miR-15a-5p | *ZYG11B* | ENSG00000162378 | -3,78 | 0,79 |
| hsa-miR-15a-5p | *MGAT5* | ENSG00000152127 | -3,78 | 0,75 |
| hsa-miR-15a-5p | *SPRYD3* | ENSG00000167778 | -3,78 | 0,74 |
| hsa-miR-15a-5p | *C12orf76* | ENSG00000174456 | -3,78 | 0,73 |
| hsa-miR-15a-5p | *ERCC6L2* | ENSG00000182150 | -3,78 | 0,71 |
| hsa-miR-15a-5p | *ATAD5* | ENSG00000176208 | -3,78 | 0,71 |
| hsa-miR-15a-5p | *SMPD1* | ENSG00000166311 | -3,78 | 0,67 |
| hsa-miR-15a-5p | *TMEM245* | ENSG00000106771 | -3,78 | 0,66 |
| hsa-miR-15a-5p | *CBX5* | ENSG00000094916 | -3,78 | 0,66 |
| hsa-miR-15a-5p | *KIF1C* | ENSG00000129250 | -3,78 | 0,66 |
| hsa-miR-15a-5p | *WEE1* | ENSG00000166483 | -3,78 | 0,66 |
| hsa-miR-15a-5p | *PAGR1* | ENSG00000280789 | -3,78 | 0,65 |
| hsa-miR-15a-5p | *TPRG1L* | ENSG00000158109 | -3,78 | 0,63 |
| hsa-miR-15a-5p | *SLC5A3* | ENSG00000198743 | -3,78 | 0,62 |
| hsa-miR-15a-5p | *TMEM55B* | ENSG00000165782 | -3,78 | 0,61 |
| hsa-miR-15a-5p | *HECTD1* | ENSG00000092148 | -3,78 | 0,58 |
| hsa-miR-15a-5p | *GCC2* | ENSG00000135968 | -3,78 | 0,56 |
| hsa-miR-15a-5p | *RBMS1* | ENSG00000153250 | -3,78 | 0,56 |
| hsa-miR-15a-5p | *NUCKS1* | ENSG00000069275 | -3,78 | 0,55 |
| hsa-miR-15a-5p | *TLE4* | ENSG00000106829 | -3,78 | 0,55 |
| hsa-miR-15a-5p | *CEP55* | ENSG00000138180 | -3,78 | 0,55 |
| hsa-miR-15a-5p | *BCL2L12* | ENSG00000126453 | -3,78 | 0,55 |
| hsa-miR-15a-5p | *MXD3* | ENSG00000213347 | -3,78 | 0,54 |
| hsa-miR-15a-5p | *INCENP* | ENSG00000149503 | -3,78 | 0,54 |
| hsa-miR-15a-5p | *KPNA3* | ENSG00000102753 | -3,78 | 0,54 |
| hsa-miR-15a-5p | *SPTBN1* | ENSG00000115306 | -3,78 | 0,52 |
| hsa-miR-15a-5p | *STK38* | ENSG00000112079 | -3,78 | 0,52 |
| hsa-miR-15a-5p | *RORA* | ENSG00000069667 | -3,78 | 0,50 |
| hsa-miR-15a-5p | *UTRN* | ENSG00000152818 | -3,78 | 0,49 |
| hsa-miR-15a-5p | *PHACTR2* | ENSG00000112419 | -3,78 | 0,49 |
| hsa-miR-15a-5p | *WIPI2* | ENSG00000157954 | -3,78 | 0,49 |
| hsa-miR-15a-5p | *MGAT4A* | ENSG00000071073 | -3,78 | 0,48 |
| hsa-miR-15a-5p | *BZW1* | ENSG00000082153 | -3,78 | 0,48 |
| hsa-miR-15a-5p | *BPTF* | ENSG00000171634 | -3,78 | 0,46 |
| hsa-miR-15a-5p | *TUBA1A* | ENSG00000167552 | -3,78 | 0,45 |
| hsa-miR-15a-5p | *TNRC6C* | ENSG00000078687 | -3,78 | 0,45 |
| hsa-miR-15a-5p | *CNOT6L* | ENSG00000138767 | -3,78 | 0,44 |
| hsa-miR-15a-5p | *RYBP* | ENSG00000163602 | -3,78 | 0,44 |
| hsa-miR-15a-5p | *PDE3B* | ENSG00000152270 | -3,78 | 0,44 |

| | | | | |
|---|---|---|---|---|
| hsa-miR-15a-5p | *PTCD3* | ENSG00000132300 | -3,78 | 0,43 |
| hsa-miR-15a-5p | *PRR11* | ENSG00000068489 | -3,78 | 0,43 |
| hsa-miR-15a-5p | *PIK3IP1* | ENSG00000100100 | -3,78 | 0,43 |
| hsa-miR-15a-5p | *TNPO1* | ENSG00000083312 | -3,78 | 0,43 |
| hsa-miR-15a-5p | *PIK3R1* | ENSG00000145675 | -3,78 | 0,42 |
| hsa-miR-15a-5p | *CDK1* | ENSG00000170312 | -3,78 | 0,42 |
| hsa-miR-15a-5p | *TSC22D3* | ENSG00000157514 | -3,78 | 0,41 |
| hsa-miR-15a-5p | *YWHAH* | ENSG00000128245 | -3,78 | 0,40 |
| hsa-miR-15a-5p | *CLSPN* | ENSG00000092853 | -3,78 | 0,39 |
| hsa-miR-15a-5p | *IPO9* | ENSG00000198700 | -3,78 | 0,39 |
| hsa-miR-15a-5p | *TPM3* | ENSG00000143549 | -3,78 | 0,39 |
| hsa-miR-15a-5p | *HIPK2* | ENSG00000064393 | -3,78 | 0,38 |
| hsa-miR-15a-5p | *MORF4L1* | ENSG00000185787 | -3,78 | 0,37 |
| hsa-miR-15a-5p | *YTHDC1* | ENSG00000083896 | -3,78 | 0,37 |
| hsa-miR-15a-5p | *PDCD4* | ENSG00000150593 | -3,78 | 0,37 |
| hsa-miR-15a-5p | *PHF19* | ENSG00000119403 | -3,78 | 0,35 |
| hsa-miR-15a-5p | *TNRC6B* | ENSG00000100354 | -3,78 | 0,34 |
| hsa-miR-15a-5p | *KIF2A* | ENSG00000068796 | -3,78 | 0,34 |
| hsa-miR-15a-5p | *WHSC1* | ENSG00000109685 | -3,78 | 0,34 |
| hsa-miR-15a-5p | *ABCF1* | ENSG00000204574 | -3,78 | 0,34 |
| hsa-miR-15a-5p | *SERBP1* | ENSG00000142864 | -3,78 | 0,33 |
| hsa-miR-15a-5p | *CEP350* | ENSG00000135837 | -3,78 | 0,32 |
| hsa-miR-15a-5p | *BCL11B* | ENSG00000127152 | -3,78 | 0,31 |
| hsa-miR-15a-5p | *LITAF* | ENSG00000189067 | -3,78 | 0,30 |
| hsa-miR-15a-5p | *CCND3* | ENSG00000112576 | -3,78 | 0,30 |
| hsa-miR-15a-5p | *TUBB* | ENSG00000196230 | -3,78 | 0,29 |
| hsa-miR-15a-5p | *DCAF7* | ENSG00000136485 | -3,78 | 0,28 |
| hsa-miR-15a-5p | *CLEC2D* | ENSG00000069493 | -3,78 | 0,26 |
| hsa-miR-15a-5p | *CDC42SE2* | ENSG00000158985 | -3,78 | 0,25 |
| hsa-miR-15b-3p | *CASK* | ENSG00000147044 | -4,45 | 0,92 |
| hsa-miR-15b-3p | *WEE1* | ENSG00000166483 | -4,45 | 0,66 |
| hsa-miR-15b-3p | *DCP2* | ENSG00000172795 | -4,45 | 0,63 |
| hsa-miR-15b-3p | *RBMS1* | ENSG00000153250 | -4,45 | 0,56 |
| hsa-miR-15b-3p | *NUCKS1* | ENSG00000069275 | -4,45 | 0,55 |
| hsa-miR-15b-3p | *SP1* | ENSG00000185591 | -4,45 | 0,55 |
| hsa-miR-15b-3p | *KPNA3* | ENSG00000102753 | -4,45 | 0,54 |
| hsa-miR-15b-3p | *USP1* | ENSG00000162607 | -4,45 | 0,50 |
| hsa-miR-15b-3p | *UTRN* | ENSG00000152818 | -4,45 | 0,49 |
| hsa-miR-15b-3p | *MGAT4A* | ENSG00000071073 | -4,45 | 0,48 |
| hsa-miR-15b-3p | *BPTF* | ENSG00000171634 | -4,45 | 0,46 |
| hsa-miR-15b-3p | *TRIM38* | ENSG00000112343 | -4,45 | 0,42 |
| hsa-miR-15b-3p | *FAM111A* | ENSG00000166801 | -4,45 | 0,42 |
| hsa-miR-15b-3p | *SMC3* | ENSG00000108055 | -4,45 | 0,41 |
| hsa-miR-15b-3p | *SMC1A* | ENSG00000072501 | -4,45 | 0,39 |
| hsa-miR-15b-3p | *NCAPG2* | ENSG00000146918 | -4,45 | 0,32 |
| hsa-miR-15b-3p | *RAD21* | ENSG00000164754 | -4,45 | 0,28 |
| hsa-miR-15b-3p | *CDC42SE2* | ENSG00000158985 | -4,45 | 0,25 |
| hsa-miR-15b-5p | *CCNE1* | ENSG00000105173 | -4,19 | 1,25 |
| hsa-miR-15b-5p | *ARL2* | ENSG00000213465 | -4,19 | 1,17 |
| hsa-miR-15b-5p | *KIF23* | ENSG00000137807 | -4,19 | 1,01 |

| | | | | |
|---|---|---|---|---|
| hsa-miR-15b-5p | *CASK* | ENSG00000147044 | -4,19 | 0,92 |
| hsa-miR-15b-5p | *TSPAN5* | ENSG00000168785 | -4,19 | 0,92 |
| hsa-miR-15b-5p | *FNIP1* | ENSG00000217128 | -4,19 | 0,86 |
| hsa-miR-15b-5p | *ANAPC13* | ENSG00000129055 | -4,19 | 0,81 |
| hsa-miR-15b-5p | *ZYG11B* | ENSG00000162378 | -4,19 | 0,79 |
| hsa-miR-15b-5p | *MGAT5* | ENSG00000152127 | -4,19 | 0,75 |
| hsa-miR-15b-5p | *SPRYD3* | ENSG00000167778 | -4,19 | 0,74 |
| hsa-miR-15b-5p | *C12orf76* | ENSG00000174456 | -4,19 | 0,73 |
| hsa-miR-15b-5p | *ATAD5* | ENSG00000176208 | -4,19 | 0,71 |
| hsa-miR-15b-5p | *SMPD1* | ENSG00000166311 | -4,19 | 0,67 |
| hsa-miR-15b-5p | *TMEM245* | ENSG00000106771 | -4,19 | 0,66 |
| hsa-miR-15b-5p | *CBX5* | ENSG00000094916 | -4,19 | 0,66 |
| hsa-miR-15b-5p | *KIF1C* | ENSG00000129250 | -4,19 | 0,66 |
| hsa-miR-15b-5p | *WEE1* | ENSG00000166483 | -4,19 | 0,66 |
| hsa-miR-15b-5p | *PAGR1* | ENSG00000280789 | -4,19 | 0,65 |
| hsa-miR-15b-5p | *TPRG1L* | ENSG00000158109 | -4,19 | 0,63 |
| hsa-miR-15b-5p | *SLC5A3* | ENSG00000198743 | -4,19 | 0,62 |
| hsa-miR-15b-5p | *TMEM55B* | ENSG00000165782 | -4,19 | 0,61 |
| hsa-miR-15b-5p | *SNRNP48* | ENSG00000168566 | -4,19 | 0,60 |
| hsa-miR-15b-5p | *HECTD1* | ENSG00000092148 | -4,19 | 0,58 |
| hsa-miR-15b-5p | *GCC2* | ENSG00000135968 | -4,19 | 0,56 |
| hsa-miR-15b-5p | *RBMS1* | ENSG00000153250 | -4,19 | 0,56 |
| hsa-miR-15b-5p | *NUCKS1* | ENSG00000069275 | -4,19 | 0,55 |
| hsa-miR-15b-5p | *TLE4* | ENSG00000106829 | -4,19 | 0,55 |
| hsa-miR-15b-5p | *CEP55* | ENSG00000138180 | -4,19 | 0,55 |
| hsa-miR-15b-5p | *BCL2L12* | ENSG00000126453 | -4,19 | 0,55 |
| hsa-miR-15b-5p | *MXD3* | ENSG00000213347 | -4,19 | 0,54 |
| hsa-miR-15b-5p | *INCENP* | ENSG00000149503 | -4,19 | 0,54 |
| hsa-miR-15b-5p | *KPNA3* | ENSG00000102753 | -4,19 | 0,54 |
| hsa-miR-15b-5p | *SPTBN1* | ENSG00000115306 | -4,19 | 0,52 |
| hsa-miR-15b-5p | *STK38* | ENSG00000112079 | -4,19 | 0,52 |
| hsa-miR-15b-5p | *RORA* | ENSG00000069667 | -4,19 | 0,50 |
| hsa-miR-15b-5p | *PALM2-AKAP2* | ENSG00000157654 | -4,19 | 0,50 |
| hsa-miR-15b-5p | *UTRN* | ENSG00000152818 | -4,19 | 0,49 |
| hsa-miR-15b-5p | *PHACTR2* | ENSG00000112419 | -4,19 | 0,49 |
| hsa-miR-15b-5p | *WIPI2* | ENSG00000157954 | -4,19 | 0,49 |
| hsa-miR-15b-5p | *MGAT4A* | ENSG00000071073 | -4,19 | 0,48 |
| hsa-miR-15b-5p | *BZW1* | ENSG00000082153 | -4,19 | 0,48 |
| hsa-miR-15b-5p | *BPTF* | ENSG00000171634 | -4,19 | 0,46 |
| hsa-miR-15b-5p | *TUBA1A* | ENSG00000167552 | -4,19 | 0,45 |
| hsa-miR-15b-5p | *TNRC6C* | ENSG00000078687 | -4,19 | 0,45 |
| hsa-miR-15b-5p | *CNOT6L* | ENSG00000138767 | -4,19 | 0,44 |
| hsa-miR-15b-5p | *RYBP* | ENSG00000163602 | -4,19 | 0,44 |
| hsa-miR-15b-5p | *PDE3B* | ENSG00000152270 | -4,19 | 0,44 |
| hsa-miR-15b-5p | *PTCD3* | ENSG00000132300 | -4,19 | 0,43 |
| hsa-miR-15b-5p | *PRR11* | ENSG00000068489 | -4,19 | 0,43 |
| hsa-miR-15b-5p | *PIK3IP1* | ENSG00000100100 | -4,19 | 0,43 |
| hsa-miR-15b-5p | *TNPO1* | ENSG00000083312 | -4,19 | 0,43 |
| hsa-miR-15b-5p | *PIK3R1* | ENSG00000145675 | -4,19 | 0,42 |
| hsa-miR-15b-5p | *CDK1* | ENSG00000170312 | -4,19 | 0,42 |

| | | | | |
|---|---|---|---|---|
| hsa-miR-15b-5p | *TSC22D3* | ENSG00000157514 | -4,19 | 0,41 |
| hsa-miR-15b-5p | *YWHAH* | ENSG00000128245 | -4,19 | 0,40 |
| hsa-miR-15b-5p | *CLSPN* | ENSG00000092853 | -4,19 | 0,39 |
| hsa-miR-15b-5p | *IPO9* | ENSG00000198700 | -4,19 | 0,39 |
| hsa-miR-15b-5p | *TPM3* | ENSG00000143549 | -4,19 | 0,39 |
| hsa-miR-15b-5p | *HIPK2* | ENSG00000064393 | -4,19 | 0,38 |
| hsa-miR-15b-5p | *PDCD4* | ENSG00000150593 | -4,19 | 0,37 |
| hsa-miR-15b-5p | *MORF4L1* | ENSG00000185787 | -4,19 | 0,37 |
| hsa-miR-15b-5p | *YTHDC1* | ENSG00000083896 | -4,19 | 0,37 |
| hsa-miR-15b-5p | *MDH1* | ENSG00000014641 | -4,19 | 0,36 |
| hsa-miR-15b-5p | *PHF19* | ENSG00000119403 | -4,19 | 0,35 |
| hsa-miR-15b-5p | *TNRC6B* | ENSG00000100354 | -4,19 | 0,34 |
| hsa-miR-15b-5p | *KIF2A* | ENSG00000068796 | -4,19 | 0,34 |
| hsa-miR-15b-5p | *WHSC1* | ENSG00000109685 | -4,19 | 0,34 |
| hsa-miR-15b-5p | *ABCF1* | ENSG00000204574 | -4,19 | 0,34 |
| hsa-miR-15b-5p | *SERBP1* | ENSG00000142864 | -4,19 | 0,33 |
| hsa-miR-15b-5p | *CEP350* | ENSG00000135837 | -4,19 | 0,32 |
| hsa-miR-15b-5p | *BCL11B* | ENSG00000127152 | -4,19 | 0,31 |
| hsa-miR-15b-5p | *LITAF* | ENSG00000189067 | -4,19 | 0,30 |
| hsa-miR-15b-5p | *CCND3* | ENSG00000112576 | -4,19 | 0,30 |
| hsa-miR-15b-5p | *TUBB* | ENSG00000196230 | -4,19 | 0,29 |
| hsa-miR-15b-5p | *DCAF7* | ENSG00000136485 | -4,19 | 0,28 |
| hsa-miR-15b-5p | *PRKCB* | ENSG00000166501 | -4,19 | 0,28 |
| hsa-miR-15b-5p | *CLEC2D* | ENSG00000069493 | -4,19 | 0,26 |
| hsa-miR-15b-5p | *CDC42SE2* | ENSG00000158985 | -4,19 | 0,25 |
| hsa-miR-16-1-3p | *CCNE1* | ENSG00000105173 | -2,39 | 1,25 |
| hsa-miR-16-1-3p | *CASK* | ENSG00000147044 | -2,39 | 0,92 |
| hsa-miR-16-1-3p | *TOP2A* | ENSG00000131747 | -2,39 | 0,72 |
| hsa-miR-16-1-3p | *SMC2* | ENSG00000136824 | -2,39 | 0,70 |
| hsa-miR-16-1-3p | *CBX5* | ENSG00000094916 | -2,39 | 0,66 |
| hsa-miR-16-1-3p | *KIF1C* | ENSG00000129250 | -2,39 | 0,66 |
| hsa-miR-16-1-3p | *WEE1* | ENSG00000166483 | -2,39 | 0,66 |
| hsa-miR-16-1-3p | *WDR76* | ENSG00000092470 | -2,39 | 0,62 |
| hsa-miR-16-1-3p | *GLCCI1* | ENSG00000106415 | -2,39 | 0,61 |
| hsa-miR-16-1-3p | *NPAT* | ENSG00000149308 | -2,39 | 0,60 |
| hsa-miR-16-1-3p | *RASA3* | ENSG00000185989 | -2,39 | 0,53 |
| hsa-miR-16-1-3p | *RORA* | ENSG00000069667 | -2,39 | 0,50 |
| hsa-miR-16-1-3p | *PDE3B* | ENSG00000152270 | -2,39 | 0,44 |
| hsa-miR-16-1-3p | *TNPO1* | ENSG00000083312 | -2,39 | 0,43 |
| hsa-miR-16-1-3p | *PIK3R1* | ENSG00000145675 | -2,39 | 0,42 |
| hsa-miR-16-1-3p | *NKTR* | ENSG00000114857 | -2,39 | 0,40 |
| hsa-miR-16-1-3p | *HELLS* | ENSG00000119969 | -2,39 | 0,38 |
| hsa-miR-16-1-3p | *YTHDC1* | ENSG00000083896 | -2,39 | 0,37 |
| hsa-miR-16-1-3p | *GLG1* | ENSG00000090863 | -2,39 | 0,35 |
| hsa-miR-16-1-3p | *BCL11B* | ENSG00000127152 | -2,39 | 0,31 |
| hsa-miR-16-2-3p | *KLF3* | ENSG00000109787 | -1,99 | 1,59 |
| hsa-miR-16-2-3p | *SNRNP48* | ENSG00000168566 | -1,99 | 0,60 |
| hsa-miR-16-2-3p | *RORA* | ENSG00000069667 | -1,99 | 0,50 |
| hsa-miR-16-2-3p | *CNOT6L* | ENSG00000138767 | -1,99 | 0,44 |
| hsa-miR-16-2-3p | *TOP1* | ENSG00000198900 | -1,99 | 0,43 |

| | | | | |
|---|---|---|---|---|
| hsa-miR-16-2-3p | *TNPO1* | ENSG00000083312 | -1,99 | 0,43 |
| hsa-miR-16-2-3p | *CDK1* | ENSG00000170312 | -1,99 | 0,42 |
| hsa-miR-16-2-3p | *ATAD2* | ENSG00000156802 | -1,99 | 0,38 |
| hsa-miR-16-2-3p | *YTHDC1* | ENSG00000083896 | -1,99 | 0,37 |
| hsa-miR-16-2-3p | *CXCR4* | ENSG00000121966 | -1,99 | 0,36 |
| hsa-miR-16-2-3p | *NCAPG2* | ENSG00000146918 | -1,99 | 0,32 |
| hsa-miR-16-2-3p | *RAD21* | ENSG00000164754 | -1,99 | 0,28 |
| hsa-miR-16-5p | *CCNE1* | ENSG00000105173 | -3,64 | 1,25 |
| hsa-miR-16-5p | *ARL2* | ENSG00000213465 | -3,64 | 1,17 |
| hsa-miR-16-5p | *KIF23* | ENSG00000137807 | -3,64 | 1,01 |
| hsa-miR-16-5p | *CASK* | ENSG00000147044 | -3,64 | 0,92 |
| hsa-miR-16-5p | *TSPAN5* | ENSG00000168785 | -3,64 | 0,92 |
| hsa-miR-16-5p | *FNIP1* | ENSG00000217128 | -3,64 | 0,86 |
| hsa-miR-16-5p | *ANAPC13* | ENSG00000129055 | -3,64 | 0,81 |
| hsa-miR-16-5p | *ZYG11B* | ENSG00000162378 | -3,64 | 0,79 |
| hsa-miR-16-5p | *MGAT5* | ENSG00000152127 | -3,64 | 0,75 |
| hsa-miR-16-5p | *SPRYD3* | ENSG00000167778 | -3,64 | 0,74 |
| hsa-miR-16-5p | *C12orf76* | ENSG00000174456 | -3,64 | 0,73 |
| hsa-miR-16-5p | *ATAD5* | ENSG00000176208 | -3,64 | 0,71 |
| hsa-miR-16-5p | *SMPD1* | ENSG00000166311 | -3,64 | 0,67 |
| hsa-miR-16-5p | *TMEM245* | ENSG00000106771 | -3,64 | 0,66 |
| hsa-miR-16-5p | *CBX5* | ENSG00000094916 | -3,64 | 0,66 |
| hsa-miR-16-5p | *KIF1C* | ENSG00000129250 | -3,64 | 0,66 |
| hsa-miR-16-5p | *WEE1* | ENSG00000166483 | -3,64 | 0,66 |
| hsa-miR-16-5p | *PAGR1* | ENSG00000280789 | -3,64 | 0,65 |
| hsa-miR-16-5p | *TPRG1L* | ENSG00000158109 | -3,64 | 0,63 |
| hsa-miR-16-5p | *SLC5A3* | ENSG00000198743 | -3,64 | 0,62 |
| hsa-miR-16-5p | *TMEM55B* | ENSG00000165782 | -3,64 | 0,61 |
| hsa-miR-16-5p | *HECTD1* | ENSG00000092148 | -3,64 | 0,58 |
| hsa-miR-16-5p | *GCC2* | ENSG00000135968 | -3,64 | 0,56 |
| hsa-miR-16-5p | *RBMS1* | ENSG00000153250 | -3,64 | 0,56 |
| hsa-miR-16-5p | *NUCKS1* | ENSG00000069275 | -3,64 | 0,55 |
| hsa-miR-16-5p | *SP1* | ENSG00000185591 | -3,64 | 0,55 |
| hsa-miR-16-5p | *TLE4* | ENSG00000106829 | -3,64 | 0,55 |
| hsa-miR-16-5p | *CEP55* | ENSG00000138180 | -3,64 | 0,55 |
| hsa-miR-16-5p | *BCL2L12* | ENSG00000126453 | -3,64 | 0,55 |
| hsa-miR-16-5p | *MXD3* | ENSG00000213347 | -3,64 | 0,54 |
| hsa-miR-16-5p | *INCENP* | ENSG00000149503 | -3,64 | 0,54 |
| hsa-miR-16-5p | *KPNA3* | ENSG00000102753 | -3,64 | 0,54 |
| hsa-miR-16-5p | *SPTBN1* | ENSG00000115306 | -3,64 | 0,52 |
| hsa-miR-16-5p | *STK38* | ENSG00000112079 | -3,64 | 0,52 |
| hsa-miR-16-5p | *RORA* | ENSG00000069667 | -3,64 | 0,50 |
| hsa-miR-16-5p | *UTRN* | ENSG00000152818 | -3,64 | 0,49 |
| hsa-miR-16-5p | *PHACTR2* | ENSG00000112419 | -3,64 | 0,49 |
| hsa-miR-16-5p | *WIPI2* | ENSG00000157954 | -3,64 | 0,49 |
| hsa-miR-16-5p | *BIRC5* | ENSG00000089685 | -3,64 | 0,48 |
| hsa-miR-16-5p | *MGAT4A* | ENSG00000071073 | -3,64 | 0,48 |
| hsa-miR-16-5p | *BZW1* | ENSG00000082153 | -3,64 | 0,48 |
| hsa-miR-16-5p | *BPTF* | ENSG00000171634 | -3,64 | 0,46 |
| hsa-miR-16-5p | *TUBA1A* | ENSG00000167552 | -3,64 | 0,45 |

| | | | | |
|---|---|---|---|---|
| hsa-miR-16-5p | *TNRC6C* | ENSG00000078687 | -3,64 | 0,45 |
| hsa-miR-16-5p | *CNOT6L* | ENSG00000138767 | -3,64 | 0,44 |
| hsa-miR-16-5p | *RYBP* | ENSG00000163602 | -3,64 | 0,44 |
| hsa-miR-16-5p | *PDE3B* | ENSG00000152270 | -3,64 | 0,44 |
| hsa-miR-16-5p | *PTCD3* | ENSG00000132300 | -3,64 | 0,43 |
| hsa-miR-16-5p | *PRR11* | ENSG00000068489 | -3,64 | 0,43 |
| hsa-miR-16-5p | *PIK3IP1* | ENSG00000100100 | -3,64 | 0,43 |
| hsa-miR-16-5p | *KIF2C* | ENSG00000142945 | -3,64 | 0,43 |
| hsa-miR-16-5p | *TNPO1* | ENSG00000083312 | -3,64 | 0,43 |
| hsa-miR-16-5p | *PIK3R1* | ENSG00000145675 | -3,64 | 0,42 |
| hsa-miR-16-5p | *CDK1* | ENSG00000170312 | -3,64 | 0,42 |
| hsa-miR-16-5p | *TSC22D3* | ENSG00000157514 | -3,64 | 0,41 |
| hsa-miR-16-5p | *YWHAH* | ENSG00000128245 | -3,64 | 0,40 |
| hsa-miR-16-5p | *CLSPN* | ENSG00000092853 | -3,64 | 0,39 |
| hsa-miR-16-5p | *IPO9* | ENSG00000198700 | -3,64 | 0,39 |
| hsa-miR-16-5p | *TPM3* | ENSG00000143549 | -3,64 | 0,39 |
| hsa-miR-16-5p | *HIPK2* | ENSG00000064393 | -3,64 | 0,38 |
| hsa-miR-16-5p | *PDCD4* | ENSG00000150593 | -3,64 | 0,37 |
| hsa-miR-16-5p | *MORF4L1* | ENSG00000185787 | -3,64 | 0,37 |
| hsa-miR-16-5p | *YTHDC1* | ENSG00000083896 | -3,64 | 0,37 |
| hsa-miR-16-5p | *PHF19* | ENSG00000119403 | -3,64 | 0,35 |
| hsa-miR-16-5p | *TNRC6B* | ENSG00000100354 | -3,64 | 0,34 |
| hsa-miR-16-5p | *KIF2A* | ENSG00000068796 | -3,64 | 0,34 |
| hsa-miR-16-5p | *WHSC1* | ENSG00000109685 | -3,64 | 0,34 |
| hsa-miR-16-5p | *ABCF1* | ENSG00000204574 | -3,64 | 0,34 |
| hsa-miR-16-5p | *SERBP1* | ENSG00000142864 | -3,64 | 0,33 |
| hsa-miR-16-5p | *CEP350* | ENSG00000135837 | -3,64 | 0,32 |
| hsa-miR-16-5p | *BCL11B* | ENSG00000127152 | -3,64 | 0,31 |
| hsa-miR-16-5p | *LITAF* | ENSG00000189067 | -3,64 | 0,30 |
| hsa-miR-16-5p | *CCND3* | ENSG00000112576 | -3,64 | 0,30 |
| hsa-miR-16-5p | *TUBB* | ENSG00000196230 | -3,64 | 0,29 |
| hsa-miR-16-5p | *DCAF7* | ENSG00000136485 | -3,64 | 0,28 |
| hsa-miR-16-5p | *PRKCB* | ENSG00000166501 | -3,64 | 0,28 |
| hsa-miR-16-5p | *CLEC2D* | ENSG00000069493 | -3,64 | 0,26 |
| hsa-miR-16-5p | *CDC42SE2* | ENSG00000158985 | -3,64 | 0,25 |

**Supplementary Table 5. List of genes downregulated (FDR < 0.05) upon miR-17~92 cluster KI**

| microRNA | Target Gene Symbol | ENSG | logFC_miRNA | logFC_mRNA |
|---|---|---|---|---|
| hsa-miR-17-3p | *PURB* | ENSG00000146676 | 1,25 | -0,61 |
| hsa-miR-17-3p | *RAP1A* | ENSG00000116473 | 1,25 | -0,49 |
| hsa-miR-17-3p | *SLC35E1* | ENSG00000127526 | 1,25 | -0,48 |
| hsa-miR-17-3p | *YTHDF2* | ENSG00000198492 | 1,25 | -0,42 |
| hsa-miR-17-3p | *SYNCRIP* | ENSG00000135316 | 1,25 | -0,36 |
| hsa-miR-17-3p | *DDX5* | ENSG00000108654 | 1,25 | -0,35 |
| hsa-miR-17-3p | *CDK2AP2* | ENSG00000167797 | 1,25 | -0,33 |
| hsa-miR-17-3p | *TWF2* | ENSG00000247596 | 1,25 | -0,32 |
| hsa-miR-17-3p | *HSP90AB1* | ENSG00000096384 | 1,25 | -0,32 |
| hsa-miR-17-3p | *RAN* | ENSG00000132341 | 1,25 | -0,32 |
| hsa-miR-17-3p | *RPL28* | ENSG00000108107 | 1,25 | -0,26 |
| hsa-miR-17-5p | *KAT2B* | ENSG00000114166 | 0,98 | -0,96 |
| hsa-miR-17-5p | *RTCA* | ENSG00000137996 | 0,98 | -0,66 |
| hsa-miR-17-5p | *TNFSF12* | ENSG00000239697 | 0,98 | -0,66 |
| hsa-miR-17-5p | *CREM* | ENSG00000095794 | 0,98 | -0,65 |
| hsa-miR-17-5p | *ORMDL3* | ENSG00000172057 | 0,98 | -0,63 |
| hsa-miR-17-5p | *PSMA5* | ENSG00000143106 | 0,98 | -0,62 |
| hsa-miR-17-5p | *PURB* | ENSG00000146676 | 0,98 | -0,61 |
| hsa-miR-17-5p | *PHTF2* | ENSG00000006576 | 0,98 | -0,59 |
| hsa-miR-17-5p | *FURIN* | ENSG00000140564 | 0,98 | -0,53 |
| hsa-miR-17-5p | *HEG1* | ENSG00000173706 | 0,98 | -0,51 |
| hsa-miR-17-5p | *SLC35E1* | ENSG00000127526 | 0,98 | -0,48 |
| hsa-miR-17-5p | *ZNF791* | ENSG00000173875 | 0,98 | -0,47 |
| hsa-miR-17-5p | *REEP5* | ENSG00000129625 | 0,98 | -0,47 |
| hsa-miR-17-5p | *BCL2* | ENSG00000171791 | 0,98 | -0,44 |
| hsa-miR-17-5p | *MLLT6* | ENSG00000275023 | 0,98 | -0,43 |
| hsa-miR-17-5p | *CCND2* | ENSG00000118971 | 0,98 | -0,42 |
| hsa-miR-17-5p | *EIF4G2* | ENSG00000110321 | 0,98 | -0,41 |
| hsa-miR-17-5p | *EEF1A1* | ENSG00000156508 | 0,98 | -0,40 |
| hsa-miR-17-5p | *LAMP1* | ENSG00000185896 | 0,98 | -0,38 |
| hsa-miR-17-5p | *STK17B* | ENSG00000081320 | 0,98 | -0,38 |
| hsa-miR-17-5p | *SYNCRIP* | ENSG00000135316 | 0,98 | -0,36 |
| hsa-miR-17-5p | *DDX5* | ENSG00000108654 | 0,98 | -0,35 |
| hsa-miR-17-5p | *NPM1* | ENSG00000181163 | 0,98 | -0,34 |
| hsa-miR-17-5p | *RSRP1* | ENSG00000117616 | 0,98 | -0,32 |
| hsa-miR-17-5p | *C1orf63* | ENSG00000117616 | 0,98 | -0,32 |
| hsa-miR-17-5p | *ZNFX1* | ENSG00000124201 | 0,98 | -0,29 |
| hsa-miR-18a-3p | *SRF* | ENSG00000112658 | 1,21 | -0,63 |
| hsa-miR-18a-3p | *ORMDL3* | ENSG00000172057 | 1,21 | -0,63 |
| hsa-miR-18a-3p | *MAD2L2* | ENSG00000116670 | 1,21 | -0,53 |
| hsa-miR-18a-3p | *PRELID1* | ENSG00000169230 | 1,21 | -0,48 |
| hsa-miR-18a-3p | *SSR3* | ENSG00000114850 | 1,21 | -0,46 |
| hsa-miR-18a-3p | *MLLT6* | ENSG00000275023 | 1,21 | -0,43 |
| hsa-miR-18a-5p | *MRPL35* | ENSG00000132313 | 1,16 | -0,86 |
| hsa-miR-18a-5p | *TBC1D9B* | ENSG00000197226 | 1,16 | -0,84 |
| hsa-miR-18a-5p | *SMCO4* | ENSG00000166002 | 1,16 | -0,74 |

| | | | | |
|---|---|---|---|---|
| hsa-miR-18a-5p | *ORMDL3* | ENSG00000172057 | 1,16 | -0,63 |
| hsa-miR-18a-5p | *PURB* | ENSG00000146676 | 1,16 | -0,61 |
| hsa-miR-18a-5p | *ITM2C* | ENSG00000135916 | 1,16 | -0,56 |
| hsa-miR-18a-5p | *HEG1* | ENSG00000173706 | 1,16 | -0,51 |
| hsa-miR-18a-5p | *RBM3* | ENSG00000102317 | 1,16 | -0,50 |
| hsa-miR-18a-5p | *RAP1A* | ENSG00000116473 | 1,16 | -0,49 |
| hsa-miR-18a-5p | *CCND2* | ENSG00000118971 | 1,16 | -0,42 |
| hsa-miR-18a-5p | *SYNCRIP* | ENSG00000135316 | 1,16 | -0,36 |
| hsa-miR-18a-5p | *DDX5* | ENSG00000108654 | 1,16 | -0,35 |
| hsa-miR-18a-5p | *ZNFX1* | ENSG00000124201 | 1,16 | -0,29 |
| hsa-miR-18a-5p | *ITM2B* | ENSG00000136156 | 1,16 | -0,28 |
| hsa-miR-18a-5p | *ACTB* | ENSG00000075624 | 1,16 | -0,26 |
| hsa-miR-19a-3p | *KAT2B* | ENSG00000114166 | 0,90 | -0,96 |
| hsa-miR-19a-3p | *NPTX1* | ENSG00000171246 | 0,90 | -0,72 |
| hsa-miR-19a-3p | *SOCS1* | ENSG00000185338 | 0,90 | -0,69 |
| hsa-miR-19a-3p | *EIF4B* | ENSG00000063046 | 0,90 | -0,68 |
| hsa-miR-19a-3p | *PAICS* | ENSG00000128050 | 0,90 | -0,67 |
| hsa-miR-19a-3p | *CREM* | ENSG00000095794 | 0,90 | -0,65 |
| hsa-miR-19a-3p | *PURB* | ENSG00000146676 | 0,90 | -0,61 |
| hsa-miR-19a-3p | *MDFIC* | ENSG00000135272 | 0,90 | -0,60 |
| hsa-miR-19a-3p | *PHTF2* | ENSG00000006576 | 0,90 | -0,59 |
| hsa-miR-19a-3p | *GRSF1* | ENSG00000132463 | 0,90 | -0,54 |
| hsa-miR-19a-3p | *FURIN* | ENSG00000140564 | 0,90 | -0,53 |
| hsa-miR-19a-3p | *HEG1* | ENSG00000173706 | 0,90 | -0,51 |
| hsa-miR-19a-3p | *RAP1A* | ENSG00000116473 | 0,90 | -0,49 |
| hsa-miR-19a-3p | *MSI2* | ENSG00000153944 | 0,90 | -0,49 |
| hsa-miR-19a-3p | *BCL3* | ENSG00000069399 | 0,90 | -0,47 |
| hsa-miR-19a-3p | *MLLT6* | ENSG00000275023 | 0,90 | -0,43 |
| hsa-miR-19a-3p | *CCND2* | ENSG00000118971 | 0,90 | -0,42 |
| hsa-miR-19a-3p | *YTHDF2* | ENSG00000198492 | 0,90 | -0,42 |
| hsa-miR-19a-3p | *EIF4G2* | ENSG00000110321 | 0,90 | -0,41 |
| hsa-miR-19a-3p | *IVNS1ABP* | ENSG00000116679 | 0,90 | -0,40 |
| hsa-miR-19a-3p | *TNFRSF1B* | ENSG00000028137 | 0,90 | -0,38 |
| hsa-miR-19a-3p | *SYNCRIP* | ENSG00000135316 | 0,90 | -0,36 |
| hsa-miR-19a-3p | *CD164* | ENSG00000135535 | 0,90 | -0,36 |
| hsa-miR-19a-3p | *NFKBIA* | ENSG00000100906 | 0,90 | -0,35 |
| hsa-miR-19a-3p | *RAN* | ENSG00000132341 | 0,90 | -0,32 |
| hsa-miR-19a-3p | *RPS4Y1* | ENSG00000129824 | 0,90 | -0,31 |
| hsa-miR-19a-3p | *HNRNPF* | ENSG00000169813 | 0,90 | -0,29 |
| hsa-miR-19a-3p | *ACTB* | ENSG00000075624 | 0,90 | -0,26 |
| hsa-miR-19a-3p | *ID2* | ENSG00000115738 | 0,90 | -0,22 |
| hsa-miR-19b-3p | *KAT2B* | ENSG00000114166 | 0,90 | -0,96 |
| hsa-miR-19b-3p | *NPTX1* | ENSG00000171246 | 0,90 | -0,72 |
| hsa-miR-19b-3p | *SOCS1* | ENSG00000185338 | 0,90 | -0,69 |
| hsa-miR-19b-3p | *EIF4B* | ENSG00000063046 | 0,90 | -0,68 |
| hsa-miR-19b-3p | *PAICS* | ENSG00000128050 | 0,90 | -0,67 |
| hsa-miR-19b-3p | *CREM* | ENSG00000095794 | 0,90 | -0,65 |
| hsa-miR-19b-3p | *PURB* | ENSG00000146676 | 0,90 | -0,61 |
| hsa-miR-19b-3p | *MDFIC* | ENSG00000135272 | 0,90 | -0,60 |
| hsa-miR-19b-3p | *PHTF2* | ENSG00000006576 | 0,90 | -0,59 |

| | | | | |
|---|---|---|---|---|
| hsa-miR-19b-3p | *GRSF1* | ENSG00000132463 | 0,90 | -0,54 |
| hsa-miR-19b-3p | *FURIN* | ENSG00000140564 | 0,90 | -0,53 |
| hsa-miR-19b-3p | *HEG1* | ENSG00000173706 | 0,90 | -0,51 |
| hsa-miR-19b-3p | *RAP1A* | ENSG00000116473 | 0,90 | -0,49 |
| hsa-miR-19b-3p | *MSI2* | ENSG00000153944 | 0,90 | -0,49 |
| hsa-miR-19b-3p | *BCL3* | ENSG00000069399 | 0,90 | -0,47 |
| hsa-miR-19b-3p | *MLLT6* | ENSG00000275023 | 0,90 | -0,43 |
| hsa-miR-19b-3p | *CCND2* | ENSG00000118971 | 0,90 | -0,42 |
| hsa-miR-19b-3p | *YTHDF2* | ENSG00000198492 | 0,90 | -0,42 |
| hsa-miR-19b-3p | *EIF4G2* | ENSG00000110321 | 0,90 | -0,41 |
| hsa-miR-19b-3p | *IVNS1ABP* | ENSG00000116679 | 0,90 | -0,40 |
| hsa-miR-19b-3p | *TNFRSF1B* | ENSG00000028137 | 0,90 | -0,38 |
| hsa-miR-19b-3p | *SYNCRIP* | ENSG00000135316 | 0,90 | -0,36 |
| hsa-miR-19b-3p | *CD164* | ENSG00000135535 | 0,90 | -0,36 |
| hsa-miR-19b-3p | *NFKBIA* | ENSG00000100906 | 0,90 | -0,35 |
| hsa-miR-19b-3p | *RAN* | ENSG00000132341 | 0,90 | -0,32 |
| hsa-miR-19b-3p | *RPS4Y1* | ENSG00000129824 | 0,90 | -0,31 |
| hsa-miR-19b-3p | *HNRNPF* | ENSG00000169813 | 0,90 | -0,29 |
| hsa-miR-19b-3p | *ACTB* | ENSG00000075624 | 0,90 | -0,26 |
| hsa-miR-19b-3p | *ID2* | ENSG00000115738 | 0,90 | -0,22 |
| hsa-miR-20a-5p | *KAT2B* | ENSG00000114166 | 1,19 | -0,96 |
| hsa-miR-20a-5p | *NPTX1* | ENSG00000171246 | 1,19 | -0,72 |
| hsa-miR-20a-5p | *GALNT6* | ENSG00000139629 | 1,19 | -0,69 |
| hsa-miR-20a-5p | *CEBPB* | ENSG00000172216 | 1,19 | -0,67 |
| hsa-miR-20a-5p | *RTCA* | ENSG00000137996 | 1,19 | -0,66 |
| hsa-miR-20a-5p | *CREM* | ENSG00000095794 | 1,19 | -0,65 |
| hsa-miR-20a-5p | *DNAJC2* | ENSG00000105821 | 1,19 | -0,65 |
| hsa-miR-20a-5p | *ORMDL3* | ENSG00000172057 | 1,19 | -0,63 |
| hsa-miR-20a-5p | *PURB* | ENSG00000146676 | 1,19 | -0,61 |
| hsa-miR-20a-5p | *PHTF2* | ENSG00000006576 | 1,19 | -0,59 |
| hsa-miR-20a-5p | *RSL1D1* | ENSG00000171490 | 1,19 | -0,58 |
| hsa-miR-20a-5p | *GRSF1* | ENSG00000132463 | 1,19 | -0,54 |
| hsa-miR-20a-5p | *FURIN* | ENSG00000140564 | 1,19 | -0,53 |
| hsa-miR-20a-5p | *HEG1* | ENSG00000173706 | 1,19 | -0,51 |
| hsa-miR-20a-5p | *RAP1A* | ENSG00000116473 | 1,19 | -0,49 |
| hsa-miR-20a-5p | *SLC35E1* | ENSG00000127526 | 1,19 | -0,48 |
| hsa-miR-20a-5p | *ZNF791* | ENSG00000173875 | 1,19 | -0,47 |
| hsa-miR-20a-5p | *REEP5* | ENSG00000129625 | 1,19 | -0,47 |
| hsa-miR-20a-5p | *BCL2* | ENSG00000171791 | 1,19 | -0,44 |
| hsa-miR-20a-5p | *MLLT6* | ENSG00000275023 | 1,19 | -0,43 |
| hsa-miR-20a-5p | *CCND2* | ENSG00000118971 | 1,19 | -0,42 |
| hsa-miR-20a-5p | *EIF4G2* | ENSG00000110321 | 1,19 | -0,41 |
| hsa-miR-20a-5p | *COMMD6* | ENSG00000188243 | 1,19 | -0,40 |
| hsa-miR-20a-5p | *IVNS1ABP* | ENSG00000116679 | 1,19 | -0,40 |
| hsa-miR-20a-5p | *LAMP1* | ENSG00000185896 | 1,19 | -0,38 |
| hsa-miR-20a-5p | *STK17B* | ENSG00000081320 | 1,19 | -0,38 |
| hsa-miR-20a-5p | *SYNCRIP* | ENSG00000135316 | 1,19 | -0,36 |
| hsa-miR-20a-5p | *DDX5* | ENSG00000108654 | 1,19 | -0,35 |
| hsa-miR-20a-5p | *NPM1* | ENSG00000181163 | 1,19 | -0,34 |
| hsa-miR-20a-5p | *RSRP1* | ENSG00000117616 | 1,19 | -0,32 |

| | | | | |
|---|---|---|---|---|
| hsa-miR-20a-5p | *C1orf63* | ENSG00000117616 | 1,19 | -0,32 |
| hsa-miR-20a-5p | *ZNFX1* | ENSG00000124201 | 1,19 | -0,29 |
| hsa-miR-92a-3p | *ADAM19* | ENSG00000135074 | 1,09 | -0,96 |
| hsa-miR-92a-3p | *KAT2B* | ENSG00000114166 | 1,09 | -0,96 |
| hsa-miR-92a-3p | *NPTX1* | ENSG00000171246 | 1,09 | -0,72 |
| hsa-miR-92a-3p | *CREM* | ENSG00000095794 | 1,09 | -0,65 |
| hsa-miR-92a-3p | *PHTF2* | ENSG00000006576 | 1,09 | -0,59 |
| hsa-miR-92a-3p | *CHCHD10* | ENSG00000250479 | 1,09 | -0,57 |
| hsa-miR-92a-3p | *HEG1* | ENSG00000173706 | 1,09 | -0,51 |
| hsa-miR-92a-3p | *RAP1A* | ENSG00000116473 | 1,09 | -0,49 |
| hsa-miR-92a-3p | *PITPNB* | ENSG00000180957 | 1,09 | -0,49 |
| hsa-miR-92a-3p | *CCND2* | ENSG00000118971 | 1,09 | -0,42 |
| hsa-miR-92a-3p | *EIF4G2* | ENSG00000110321 | 1,09 | -0,41 |
| hsa-miR-92a-3p | *RPL9* | ENSG00000163682 | 1,09 | -0,35 |
| hsa-miR-92a-3p | *NPM1* | ENSG00000181163 | 1,09 | -0,34 |
| hsa-miR-92a-3p | *RPL15* | ENSG00000174748 | 1,09 | -0,31 |
| hsa-miR-92a-3p | *ITM2B* | ENSG00000136156 | 1,09 | -0,28 |
| hsa-miR-92a-3p | *MYH9* | ENSG00000100345 | 1,09 | -0,26 |
| hsa-miR-92b-3p | *RPL9* | ENSG00000163682 | 0,36 | -0,35 |
| hsa-miR-92a-3p | *RPS20* | ENSG00000008988 | 1,09 | -0,17 |
| hsa-miR-92b-3p | *ADAM19* | ENSG00000135074 | 0,36 | -0,96 |
| hsa-miR-92b-3p | *KAT2B* | ENSG00000114166 | 0,36 | -0,96 |
| hsa-miR-92b-3p | *NPTX1* | ENSG00000171246 | 0,36 | -0,72 |
| hsa-miR-92b-3p | *CREM* | ENSG00000095794 | 0,36 | -0,65 |
| hsa-miR-92b-3p | *PHTF2* | ENSG00000006576 | 0,36 | -0,59 |
| hsa-miR-92b-3p | *CHCHD10* | ENSG00000250479 | 0,36 | -0,57 |
| hsa-miR-92b-3p | *HEG1* | ENSG00000173706 | 0,36 | -0,51 |
| hsa-miR-92b-3p | *RAP1A* | ENSG00000116473 | 0,36 | -0,49 |
| hsa-miR-92b-3p | *PITPNB* | ENSG00000180957 | 0,36 | -0,49 |
| hsa-miR-92b-3p | *CCND2* | ENSG00000118971 | 0,36 | -0,42 |
| hsa-miR-92b-3p | *EIF4G2* | ENSG00000110321 | 0,36 | -0,41 |
| hsa-miR-92b-3p | *RPL15* | ENSG00000174748 | 0,36 | -0,31 |
| hsa-miR-92b-3p | *ITM2B* | ENSG00000136156 | 0,36 | -0,28 |
| hsa-miR-92b-3p | *MYH9* | ENSG00000100345 | 0,36 | -0,26 |
| hsa-miR-92b-3p | *TPT1* | ENSG00000133112 | 0,36 | -0,23 |
| hsa-miR-92b-3p | *RPS20* | ENSG00000008988 | 0,36 | -0,17 |

**Supplementary Table 6. List of significantly deregulated genes upon *castling* of miR-17~92 cluster into the miR-15a/16-1 locus**

| Gene Symbol | RPM | | | | | | Fold Change (FC) *castling* /Mock | logFC | p-value | FDR |
|---|---|---|---|---|---|---|---|---|---|---|
| | Day 10 | | | | | | | | | |
| | Mock | | | *castling* | | | | | | |
| | Mock #1 | Mock #2 | Mean | miR-17~92 *castling* #1 | miR-17~92 *castling* #2 | Mean | | | | |
| ***AUTS2*** | 0,00 | 0,60 | 0,30 | 9,42 | 18,51 | 13,97 | 46,24 | 5,38 | 0,0000 | 0,0003 |
| ***MT1G*** | 0,56 | 0,20 | 0,38 | 17,71 | 17,22 | 17,46 | 46,14 | 5,35 | 0,0000 | 0,0000 |
| ***IGHM*** | 0,00 | 0,81 | 0,40 | 17,14 | 16,71 | 16,92 | 42,04 | 5,27 | 0,0000 | 0,0000 |
| ***PAX5*** | 0,00 | 1,21 | 0,60 | 15,14 | 7,71 | 11,42 | 18,93 | 4,15 | 0,0008 | 0,0120 |
| ***PAQR6*** | 0,37 | 1,41 | 0,89 | 8,28 | 23,91 | 16,09 | 18,09 | 4,09 | 0,0000 | 0,0008 |
| ***CD244*** | 1,30 | 0,40 | 0,85 | 11,71 | 16,71 | 14,21 | 16,73 | 3,95 | 0,0002 | 0,0034 |
| ***P2RY11*** | 1,48 | 2,42 | 1,95 | 17,71 | 32,90 | 25,30 | 12,99 | 3,62 | 0,0000 | 0,0000 |
| ***FGFBP2*** | 2,04 | 1,41 | 1,72 | 22,28 | 13,62 | 17,95 | 10,42 | 3,30 | 0,0001 | 0,0023 |
| ***7SK*** | 20,20 | 15,09 | 17,64 | 177,06 | 182,76 | 179,91 | 10,20 | 3,29 | 0,0000 | 0,0000 |
| ***PXDN*** | 2,59 | 0,60 | 1,60 | 12,28 | 19,79 | 16,04 | 10,03 | 3,23 | 0,0004 | 0,0063 |
| ***TBKBP1*** | 1,67 | 2,62 | 2,14 | 18,28 | 21,34 | 19,81 | 9,25 | 3,14 | 0,0001 | 0,0016 |
| ***TRBV28*** | 23,16 | 38,63 | 30,90 | 283,87 | 276,32 | 280,10 | 9,07 | 3,12 | 0,0000 | 0,0000 |
| ***GP6-AS1*** | 3,34 | 3,02 | 3,18 | 16,85 | 40,61 | 28,73 | 9,04 | 3,11 | 0,0000 | 0,0000 |
| ***KLHL25*** | 2,59 | 0,81 | 1,70 | 18,85 | 11,57 | 15,21 | 8,95 | 3,07 | 0,0009 | 0,0128 |
| ***PRKD3*** | 7,78 | 1,41 | 4,60 | 27,99 | 51,41 | 39,70 | 8,64 | 3,04 | 0,0000 | 0,0000 |
| ***U2*** | 0,56 | 3,22 | 1,89 | 17,14 | 14,91 | 16,02 | 8,49 | 3,02 | 0,0007 | 0,0105 |
| ***PLAU*** | 2,04 | 5,03 | 3,53 | 31,99 | 26,73 | 29,36 | 8,31 | 2,99 | 0,0000 | 0,0000 |
| ***TRGC2*** | 26,87 | 11,27 | 19,07 | 163,64 | 122,35 | 143,00 | 7,50 | 2,84 | 0,0000 | 0,0000 |
| ***PIK3AP1*** | 3,52 | 0,00 | 1,76 | 7,71 | 18,51 | 13,11 | 7,45 | 2,80 | 0,0046 | 0,0437 |
| ***IGLC7*** | 0,37 | 3,42 | 1,90 | 16,85 | 10,28 | 13,57 | 7,15 | 2,77 | 0,0041 | 0,0405 |
| ***RNU4-2*** | 2,22 | 4,43 | 3,33 | 17,99 | 25,45 | 21,72 | 6,53 | 2,64 | 0,0002 | 0,0037 |
| ***PECAM1*** | 5,93 | 5,43 | 5,68 | 40,27 | 33,93 | 37,10 | 6,53 | 2,64 | 0,0000 | 0,0000 |
| ***F8A1*** | 2,78 | 1,61 | 2,19 | 7,14 | 21,08 | 14,11 | 6,43 | 2,61 | 0,0048 | 0,0449 |
| ***ZKSCAN2-DT*** | 2,78 | 4,63 | 3,70 | 14,85 | 30,33 | 22,59 | 6,10 | 2,54 | 0,0002 | 0,0039 |
| ***MRC2*** | 9,45 | 23,34 | 16,40 | 77,39 | 118,50 | 97,95 | 5,97 | 2,52 | 0,0000 | 0,0000 |
| ***KLRC1*** | 4,26 | 7,04 | 5,65 | 37,13 | 29,82 | 33,47 | 5,92 | 2,50 | 0,0000 | 0,0001 |
| ***Y_RNA*** | 7,60 | 2,82 | 5,21 | 33,41 | 25,70 | 29,56 | 5,68 | 2,44 | 0,0000 | 0,0007 |
| ***CCL5*** | 784,56 | 680,10 | 732,33 | 4114,99 | 3990,84 | 4052,92 | 5,53 | 2,41 | 0,0000 | 0,0000 |
| ***LINC00944*** | 3,52 | 2,82 | 3,17 | 23,13 | 11,82 | 17,48 | 5,52 | 2,39 | 0,0024 | 0,0275 |
| ***SLC25A53*** | 1,48 | 4,83 | 3,16 | 7,71 | 26,99 | 17,35 | 5,50 | 2,39 | 0,0025 | 0,0283 |
| ***ITGA1*** | 77,83 | 107,25 | 92,54 | 442,37 | 563,18 | 502,78 | 5,43 | 2,38 | 0,0000 | 0,0000 |
| ***KLRC2*** | 40,58 | 55,74 | 48,16 | 253,89 | 265,53 | 259,71 | 5,39 | 2,37 | 0,0000 | 0,0000 |
| ***RGS17P1*** | 4,63 | 2,42 | 3,52 | 10,28 | 26,99 | 18,64 | 5,29 | 2,33 | 0,0019 | 0,0234 |
| ***PITPNC1*** | 10,01 | 6,24 | 8,12 | 26,56 | 56,04 | 41,30 | 5,08 | 2,28 | 0,0000 | 0,0001 |
| ***ZNF683*** | 11,67 | 5,63 | 8,65 | 53,12 | 34,70 | 43,91 | 5,07 | 2,28 | 0,0000 | 0,0000 |
| ***CXXC5*** | 3,52 | 7,85 | 5,68 | 24,85 | 31,62 | 28,23 | 4,97 | 2,25 | 0,0001 | 0,0024 |

| | | | | | | | | | | |
|---|---|---|---|---|---|---|---|---|---|---|
| ***SRGAP3*** | 23,53 | 26,16 | 24,85 | 128,23 | 107,44 | 117,84 | 4,74 | 2,19 | 0,0000 | 0,0000 |
| ***PDGFA*** | 7,60 | 10,06 | 8,83 | 41,70 | 39,33 | 40,51 | 4,59 | 2,14 | 0,0000 | 0,0002 |
| ***CAPS*** | 4,08 | 3,82 | 3,95 | 13,14 | 22,36 | 17,75 | 4,49 | 2,10 | 0,0054 | 0,0494 |
| ***DAPK2*** | 9,64 | 11,27 | 10,45 | 34,27 | 57,84 | 46,05 | 4,41 | 2,08 | 0,0000 | 0,0001 |
| ***DGKD*** | 7,60 | 16,50 | 12,05 | 36,84 | 69,14 | 52,99 | 4,40 | 2,08 | 0,0000 | 0,0000 |
| ***TRGC1*** | 7,04 | 6,84 | 6,94 | 28,27 | 31,62 | 29,94 | 4,31 | 2,04 | 0,0002 | 0,0039 |
| ***KLRC4*** | 24,83 | 29,98 | 27,41 | 121,09 | 112,84 | 116,97 | 4,27 | 2,03 | 0,0000 | 0,0000 |
| ***SEMA7A*** | 4,82 | 5,84 | 5,33 | 26,27 | 18,51 | 22,39 | 4,20 | 2,01 | 0,0020 | 0,0239 |
| ***HOPX*** | 39,10 | 31,59 | 35,34 | 110,24 | 185,07 | 147,65 | 4,18 | 2,00 | 0,0000 | 0,0000 |
| ***ZNF549*** | 5,56 | 3,82 | 4,69 | 22,85 | 16,19 | 19,52 | 4,16 | 1,99 | 0,0046 | 0,0440 |
| ***NT5C3AP2*** | 12,04 | 18,11 | 15,08 | 42,55 | 80,97 | 61,76 | 4,10 | 1,97 | 0,0000 | 0,0000 |
| ***LPAR5*** | 8,15 | 10,46 | 9,31 | 41,12 | 34,19 | 37,66 | 4,05 | 1,95 | 0,0000 | 0,0011 |
| ***LY9*** | 7,23 | 10,26 | 8,74 | 34,27 | 36,24 | 35,26 | 4,03 | 1,95 | 0,0001 | 0,0019 |
| ***CD8B*** | 48,55 | 44,47 | 46,51 | 165,35 | 208,98 | 187,16 | 4,02 | 1,95 | 0,0000 | 0,0000 |
| ***SLAMF7*** | 49,85 | 64,79 | 57,32 | 217,04 | 238,54 | 227,79 | 3,97 | 1,93 | 0,0000 | 0,0000 |
| ***GATD1-DT*** | 3,89 | 11,67 | 7,78 | 28,27 | 32,39 | 30,33 | 3,90 | 1,90 | 0,0004 | 0,0067 |
| ***INTS3*** | 8,52 | 9,46 | 8,99 | 46,55 | 23,13 | 34,84 | 3,88 | 1,89 | 0,0001 | 0,0028 |
| ***ITGAE*** | 57,63 | 77,87 | 67,75 | 282,16 | 235,45 | 258,80 | 3,82 | 1,87 | 0,0000 | 0,0000 |
| ***TTC24*** | 10,01 | 13,08 | 11,54 | 27,42 | 59,38 | 43,40 | 3,76 | 1,85 | 0,0000 | 0,0007 |
| ***PPAN*** | 7,78 | 4,23 | 6,00 | 20,56 | 23,65 | 22,11 | 3,68 | 1,81 | 0,0045 | 0,0432 |
| ***GPR157*** | 12,97 | 10,66 | 11,82 | 47,69 | 38,81 | 43,25 | 3,66 | 1,81 | 0,0000 | 0,0009 |
| ***KLRB1*** | 29,83 | 32,19 | 31,01 | 140,51 | 74,80 | 107,65 | 3,47 | 1,74 | 0,0000 | 0,0000 |
| ***MT1X*** | 11,86 | 4,43 | 8,14 | 22,85 | 33,42 | 28,13 | 3,45 | 1,72 | 0,0017 | 0,0214 |
| ***GABARAPL1*** | 63,56 | 68,21 | 65,88 | 225,04 | 225,94 | 225,49 | 3,42 | 1,72 | 0,0000 | 0,0000 |
| ***ZFHX2-AS1*** | 11,12 | 3,82 | 7,47 | 21,13 | 29,56 | 25,35 | 3,39 | 1,70 | 0,0035 | 0,0363 |
| ***AOAH*** | 58,74 | 65,39 | 62,07 | 192,20 | 222,86 | 207,53 | 3,34 | 1,68 | 0,0000 | 0,0000 |
| ***CD8A*** | 127,49 | 118,11 | 122,80 | 415,24 | 393,53 | 404,39 | 3,29 | 1,66 | 0,0000 | 0,0000 |
| ***BMF*** | 30,20 | 15,49 | 22,85 | 78,82 | 71,46 | 75,14 | 3,29 | 1,66 | 0,0000 | 0,0000 |
| ***LINC01012*** | 8,89 | 7,65 | 8,27 | 17,71 | 36,50 | 27,10 | 3,28 | 1,65 | 0,0030 | 0,0325 |
| ***RIN3*** | 21,12 | 20,93 | 21,03 | 66,83 | 70,94 | 68,89 | 3,28 | 1,65 | 0,0000 | 0,0000 |
| ***JAML*** | 283,51 | 278,48 | 280,99 | 825,34 | 1004,01 | 914,67 | 3,26 | 1,64 | 0,0000 | 0,0000 |
| ***ARL2*** | 35,95 | 22,54 | 29,24 | 91,39 | 98,45 | 94,92 | 3,25 | 1,64 | 0,0000 | 0,0000 |
| ***NHS*** | 19,64 | 19,72 | 19,68 | 52,83 | 74,80 | 63,82 | 3,24 | 1,64 | 0,0000 | 0,0001 |
| ***SNAPC4*** | 6,67 | 10,87 | 8,77 | 26,27 | 30,07 | 28,17 | 3,21 | 1,62 | 0,0028 | 0,0306 |
| ***CD101*** | 15,01 | 11,07 | 13,04 | 45,12 | 37,79 | 41,45 | 3,18 | 1,61 | 0,0002 | 0,0044 |
| ***MPZL3*** | 64,67 | 68,82 | 66,74 | 213,05 | 210,78 | 211,91 | 3,18 | 1,61 | 0,0000 | 0,0000 |
| ***TRIB2*** | 7,97 | 13,68 | 10,83 | 16,85 | 50,64 | 33,74 | 3,12 | 1,58 | 0,0012 | 0,0162 |
| ***AKAP5*** | 8,15 | 13,08 | 10,62 | 32,84 | 32,64 | 32,74 | 3,08 | 1,56 | 0,0016 | 0,0200 |
| ***MTFP1*** | 10,01 | 12,88 | 11,44 | 31,41 | 38,81 | 35,11 | 3,07 | 1,56 | 0,0011 | 0,0148 |
| ***SYTL2*** | 84,68 | 89,34 | 87,01 | 261,88 | 268,61 | 265,25 | 3,05 | 1,55 | 0,0000 | 0,0000 |
| ***RGS16*** | 65,04 | 66,40 | 65,72 | 204,48 | 193,30 | 198,89 | 3,03 | 1,54 | 0,0000 | 0,0000 |
| ***PRKAR1B*** | 9,82 | 12,88 | 11,35 | 46,27 | 22,11 | 34,19 | 3,01 | 1,53 | 0,0015 | 0,0190 |
| ***P2RX5*** | 10,56 | 11,67 | 11,12 | 30,56 | 34,96 | 32,76 | 2,95 | 1,50 | 0,0023 | 0,0264 |
| ***LRRC28*** | 21,31 | 28,17 | 24,74 | 65,97 | 79,43 | 72,70 | 2,94 | 1,49 | 0,0000 | 0,0001 |
| ***SLC9B2*** | 8,89 | 11,07 | 9,98 | 26,56 | 31,87 | 29,22 | 2,93 | 1,49 | 0,0044 | 0,0425 |
| ***TRAV17*** | 8,15 | 16,10 | 12,13 | 42,55 | 28,28 | 35,41 | 2,92 | 1,49 | 0,0015 | 0,0197 |
| ***CRACR2A*** | 8,15 | 12,88 | 10,52 | 25,13 | 36,24 | 30,69 | 2,92 | 1,48 | 0,0035 | 0,0362 |

| | | | | | | | | | | |
|---|---|---|---|---|---|---|---|---|---|---|
| ***SLC12A9*** | 9,45 | 22,94 | 16,19 | 47,12 | 47,30 | 47,21 | 2,92 | 1,48 | 0,0002 | 0,0041 |
| ***LINC02481*** | 46,88 | 30,58 | 38,73 | 104,81 | 119,27 | 112,04 | 2,89 | 1,47 | 0,0000 | 0,0000 |
| ***KIT*** | 22,05 | 21,13 | 21,59 | 65,11 | 59,63 | 62,37 | 2,89 | 1,47 | 0,0000 | 0,0006 |
| ***SCML4*** | 17,60 | 12,88 | 15,24 | 36,84 | 50,89 | 43,87 | 2,88 | 1,46 | 0,0005 | 0,0073 |
| ***SNORD3A*** | 100,99 | 85,72 | 93,35 | 282,44 | 247,79 | 265,12 | 2,84 | 1,45 | 0,0000 | 0,0000 |
| ***CLNK*** | 8,34 | 20,73 | 14,53 | 45,41 | 36,76 | 41,08 | 2,83 | 1,44 | 0,0008 | 0,0119 |
| ***DBN1*** | 35,21 | 15,29 | 25,25 | 56,83 | 85,85 | 71,34 | 2,83 | 1,44 | 0,0000 | 0,0003 |
| ***GRAP*** | 8,52 | 15,09 | 11,81 | 37,98 | 28,53 | 33,26 | 2,82 | 1,43 | 0,0030 | 0,0320 |
| ***TNFRSF25*** | 18,35 | 18,31 | 18,33 | 44,55 | 58,61 | 51,58 | 2,81 | 1,43 | 0,0002 | 0,0034 |
| ***MTERF4*** | 31,32 | 25,96 | 28,64 | 83,68 | 77,11 | 80,39 | 2,81 | 1,43 | 0,0000 | 0,0001 |
| ***GAB3*** | 23,16 | 7,65 | 15,40 | 40,55 | 44,73 | 42,64 | 2,77 | 1,41 | 0,0008 | 0,0119 |
| ***TTLL3*** | 24,09 | 14,29 | 19,19 | 35,13 | 70,94 | 53,04 | 2,76 | 1,40 | 0,0002 | 0,0034 |
| ***LINC02908*** | 15,57 | 12,88 | 14,22 | 47,41 | 30,85 | 39,13 | 2,75 | 1,40 | 0,0015 | 0,0190 |
| ***ABLIM1*** | 18,72 | 40,65 | 29,68 | 82,82 | 78,91 | 80,87 | 2,72 | 1,39 | 0,0000 | 0,0002 |
| ***EMB*** | 72,64 | 85,92 | 79,28 | 219,04 | 212,83 | 215,94 | 2,72 | 1,39 | 0,0000 | 0,0000 |
| ***PARVG*** | 13,90 | 13,28 | 13,59 | 32,56 | 41,13 | 36,84 | 2,71 | 1,38 | 0,0024 | 0,0272 |
| ***PTPN6*** | 26,50 | 45,07 | 35,79 | 99,10 | 94,59 | 96,85 | 2,71 | 1,38 | 0,0000 | 0,0000 |
| ***SMOX*** | 15,38 | 15,90 | 15,64 | 41,41 | 42,93 | 42,17 | 2,70 | 1,37 | 0,0011 | 0,0154 |
| ***ZNF544*** | 11,86 | 14,89 | 13,37 | 32,56 | 39,07 | 35,81 | 2,68 | 1,36 | 0,0031 | 0,0327 |
| ***SYCP2*** | 23,72 | 21,73 | 22,72 | 52,26 | 67,35 | 59,80 | 2,63 | 1,33 | 0,0001 | 0,0027 |
| ***SLC2A8*** | 18,90 | 23,14 | 21,02 | 53,12 | 57,06 | 55,09 | 2,62 | 1,33 | 0,0003 | 0,0047 |
| ***ZNF652*** | 46,33 | 54,73 | 50,53 | 123,94 | 139,57 | 131,76 | 2,61 | 1,32 | 0,0000 | 0,0000 |
| ***APOBR*** | 20,57 | 19,92 | 20,24 | 30,56 | 74,54 | 52,55 | 2,60 | 1,31 | 0,0004 | 0,0069 |
| ***LPAR2*** | 18,35 | 12,68 | 15,51 | 28,27 | 51,41 | 39,84 | 2,57 | 1,30 | 0,0025 | 0,0286 |
| ***SLFN13*** | 14,82 | 15,09 | 14,96 | 23,70 | 52,69 | 38,20 | 2,55 | 1,29 | 0,0033 | 0,0348 |
| ***IKZF3*** | 89,50 | 86,52 | 88,01 | 192,77 | 253,96 | 223,36 | 2,54 | 1,28 | 0,0000 | 0,0000 |
| ***NELL2*** | 62,45 | 65,39 | 63,92 | 157,36 | 166,31 | 161,83 | 2,53 | 1,28 | 0,0000 | 0,0000 |
| ***CAPRIN2*** | 16,86 | 22,94 | 19,90 | 41,41 | 59,12 | 50,27 | 2,53 | 1,28 | 0,0008 | 0,0112 |
| ***GNLY*** | 1041,01 | 1046,51 | 1043,76 | 2598,53 | 2658,59 | 2628,56 | 2,52 | 1,27 | 0,0000 | 0,0000 |
| ***TPT1-AS1*** | 40,03 | 39,44 | 39,73 | 110,81 | 88,68 | 99,74 | 2,51 | 1,27 | 0,0000 | 0,0001 |
| ***PRKCH-AS1*** | 13,16 | 17,30 | 15,23 | 38,27 | 38,04 | 38,16 | 2,51 | 1,26 | 0,0039 | 0,0391 |
| ***MYO1F*** | 83,94 | 88,33 | 86,14 | 219,04 | 212,06 | 215,55 | 2,50 | 1,26 | 0,0000 | 0,0000 |
| ***KLRD1*** | 78,20 | 84,91 | 81,55 | 181,63 | 225,94 | 203,79 | 2,50 | 1,26 | 0,0000 | 0,0000 |
| ***ARRB1*** | 19,09 | 22,94 | 21,01 | 42,55 | 61,69 | 52,12 | 2,48 | 1,25 | 0,0008 | 0,0111 |
| ***FCMR*** | 39,65 | 23,54 | 31,60 | 88,25 | 68,37 | 78,31 | 2,48 | 1,25 | 0,0000 | 0,0009 |
| ***RHBDF2*** | 17,97 | 17,30 | 17,64 | 40,55 | 46,78 | 43,67 | 2,48 | 1,25 | 0,0022 | 0,0256 |
| ***WAKMAR2*** | 26,50 | 35,62 | 31,06 | 87,10 | 65,80 | 76,45 | 2,46 | 1,24 | 0,0001 | 0,0012 |
| ***VEGFB*** | 19,64 | 17,71 | 18,67 | 52,55 | 38,30 | 45,42 | 2,43 | 1,22 | 0,0021 | 0,0248 |
| ***CTSD*** | 162,14 | 168,01 | 165,08 | 352,70 | 449,83 | 401,26 | 2,43 | 1,22 | 0,0000 | 0,0000 |
| ***DEDD2*** | 25,20 | 20,52 | 22,86 | 60,83 | 50,12 | 55,48 | 2,43 | 1,22 | 0,0007 | 0,0100 |
| ***NT5E*** | 16,68 | 18,51 | 17,59 | 48,84 | 36,50 | 42,67 | 2,43 | 1,22 | 0,0030 | 0,0321 |
| ***UBASH3B*** | 20,75 | 21,73 | 21,24 | 49,69 | 52,69 | 51,19 | 2,41 | 1,21 | 0,0012 | 0,0158 |
| ***PLEKHA1*** | 34,28 | 26,96 | 30,62 | 69,40 | 77,88 | 73,64 | 2,40 | 1,21 | 0,0001 | 0,0022 |
| ***CD9*** | 33,73 | 26,36 | 30,04 | 84,82 | 58,35 | 71,58 | 2,38 | 1,19 | 0,0001 | 0,0029 |
| ***DGCR8*** | 15,01 | 23,34 | 19,18 | 43,98 | 47,04 | 45,51 | 2,37 | 1,19 | 0,0027 | 0,0295 |
| ***SLC39A10*** | 18,16 | 19,32 | 18,74 | 41,41 | 47,04 | 44,22 | 2,36 | 1,18 | 0,0033 | 0,0344 |
| ***GPR68*** | 83,20 | 68,21 | 75,71 | 190,77 | 166,31 | 178,54 | 2,36 | 1,18 | 0,0000 | 0,0000 |

| | | | | | | | | | | |
|---|---|---|---|---|---|---|---|---|---|---|
| ***MGAT5*** | 28,91 | 14,49 | 21,70 | 61,40 | 40,36 | 50,88 | 2,34 | 1,17 | 0,0017 | 0,0209 |
| ***FUCA1*** | 24,65 | 25,35 | 25,00 | 69,68 | 47,04 | 58,36 | 2,33 | 1,16 | 0,0008 | 0,0115 |
| ***AK1*** | 23,72 | 25,55 | 24,64 | 68,54 | 46,27 | 57,40 | 2,33 | 1,16 | 0,0009 | 0,0127 |
| ***ARFGAP2*** | 31,50 | 23,34 | 27,42 | 65,97 | 61,43 | 63,70 | 2,32 | 1,16 | 0,0005 | 0,0078 |
| ***UBL3*** | 35,02 | 30,79 | 32,90 | 72,54 | 80,20 | 76,37 | 2,32 | 1,15 | 0,0001 | 0,0028 |
| ***EEIG1*** | 123,78 | 134,81 | 129,30 | 285,87 | 311,79 | 298,83 | 2,31 | 1,15 | 0,0000 | 0,0000 |
| ***LINC00623*** | 18,16 | 20,32 | 19,24 | 38,84 | 49,87 | 44,35 | 2,31 | 1,14 | 0,0040 | 0,0401 |
| ***LINC00996*** | 61,33 | 55,33 | 58,33 | 123,66 | 143,17 | 133,42 | 2,29 | 1,13 | 0,0000 | 0,0000 |
| ***CD27*** | 31,69 | 44,07 | 37,88 | 110,52 | 62,72 | 86,62 | 2,29 | 1,13 | 0,0001 | 0,0015 |
| ***STK38*** | 63,93 | 53,12 | 58,52 | 135,37 | 129,81 | 132,59 | 2,27 | 1,12 | 0,0000 | 0,0000 |
| ***CAPN7*** | 22,24 | 21,93 | 22,08 | 51,98 | 47,81 | 49,89 | 2,26 | 1,12 | 0,0028 | 0,0306 |
| ***NECTIN3*** | 46,70 | 39,24 | 42,97 | 117,95 | 76,09 | 97,02 | 2,26 | 1,12 | 0,0000 | 0,0008 |
| ***STK38L*** | 29,28 | 19,52 | 24,40 | 30,84 | 78,66 | 54,75 | 2,24 | 1,10 | 0,0019 | 0,0230 |
| ***YPEL2*** | 33,73 | 37,02 | 35,37 | 70,54 | 87,91 | 79,22 | 2,24 | 1,10 | 0,0002 | 0,0036 |
| ***LDLRAD4*** | 69,30 | 74,45 | 71,88 | 133,37 | 188,41 | 160,89 | 2,24 | 1,10 | 0,0000 | 0,0000 |
| ***SLAMF6*** | 19,83 | 30,38 | 25,11 | 47,98 | 64,26 | 56,12 | 2,24 | 1,10 | 0,0017 | 0,0212 |
| ***IRF2BPL*** | 30,76 | 28,98 | 29,87 | 80,54 | 52,95 | 66,74 | 2,23 | 1,10 | 0,0006 | 0,0095 |
| ***EPB41*** | 108,22 | 132,40 | 120,31 | 289,01 | 245,73 | 267,37 | 2,22 | 1,09 | 0,0000 | 0,0000 |
| ***PASK*** | 29,46 | 39,64 | 34,55 | 80,25 | 73,26 | 76,75 | 2,22 | 1,09 | 0,0003 | 0,0048 |
| ***ABHD3*** | 23,72 | 26,96 | 25,34 | 63,97 | 48,58 | 56,28 | 2,22 | 1,09 | 0,0018 | 0,0222 |
| ***SMIM27*** | 32,06 | 30,79 | 31,42 | 53,12 | 86,37 | 69,74 | 2,22 | 1,09 | 0,0005 | 0,0083 |
| ***ABHD14B*** | 30,39 | 14,69 | 22,54 | 67,40 | 32,64 | 50,02 | 2,22 | 1,09 | 0,0033 | 0,0350 |
| ***CRNDE*** | 14,45 | 29,58 | 22,02 | 41,41 | 56,29 | 48,85 | 2,22 | 1,09 | 0,0037 | 0,0376 |
| ***ADCY3*** | 17,23 | 29,98 | 23,61 | 47,12 | 57,06 | 52,09 | 2,21 | 1,08 | 0,0029 | 0,0315 |
| ***CCDC12*** | 34,28 | 34,21 | 34,24 | 84,53 | 66,57 | 75,55 | 2,21 | 1,08 | 0,0003 | 0,0058 |
| ***RNF157*** | 31,13 | 15,49 | 23,31 | 46,27 | 56,29 | 51,28 | 2,20 | 1,08 | 0,0033 | 0,0347 |
| ***ATP10D*** | 32,61 | 40,65 | 36,63 | 72,82 | 87,91 | 80,37 | 2,19 | 1,07 | 0,0002 | 0,0044 |
| ***GLUL*** | 18,72 | 34,81 | 26,76 | 55,40 | 61,95 | 58,68 | 2,19 | 1,07 | 0,0017 | 0,0211 |
| ***DIP2A*** | 60,59 | 63,18 | 61,89 | 111,66 | 159,37 | 135,52 | 2,19 | 1,07 | 0,0000 | 0,0001 |
| ***SKIL*** | 35,76 | 34,41 | 35,09 | 77,97 | 75,31 | 76,64 | 2,18 | 1,07 | 0,0004 | 0,0061 |
| ***FHL3*** | 20,01 | 24,35 | 22,18 | 22,85 | 74,03 | 48,44 | 2,18 | 1,07 | 0,0046 | 0,0441 |
| ***G2E3*** | 28,72 | 27,16 | 27,94 | 43,98 | 77,88 | 60,93 | 2,18 | 1,06 | 0,0015 | 0,0192 |
| ***MIAT*** | 190,86 | 189,14 | 190,00 | 378,97 | 448,80 | 413,88 | 2,18 | 1,06 | 0,0000 | 0,0000 |
| ***GCNT1*** | 33,54 | 29,38 | 31,46 | 87,67 | 49,10 | 68,38 | 2,17 | 1,06 | 0,0008 | 0,0115 |
| ***LINC02446*** | 136,38 | 144,27 | 140,33 | 287,30 | 320,53 | 303,92 | 2,17 | 1,06 | 0,0000 | 0,0000 |
| ***PLEKHO1*** | 143,05 | 130,19 | 136,62 | 274,16 | 316,93 | 295,55 | 2,16 | 1,05 | 0,0000 | 0,0000 |
| ***CSGALNACT1*** | 46,88 | 38,63 | 42,76 | 82,53 | 102,05 | 92,29 | 2,16 | 1,05 | 0,0001 | 0,0025 |
| ***IRF3*** | 54,29 | 52,11 | 53,20 | 126,23 | 102,56 | 114,39 | 2,15 | 1,04 | 0,0000 | 0,0006 |
| ***SNX9*** | 56,15 | 49,10 | 52,62 | 91,39 | 134,69 | 113,04 | 2,15 | 1,04 | 0,0000 | 0,0006 |
| ***ZNF84*** | 29,46 | 23,54 | 26,50 | 50,26 | 63,49 | 56,88 | 2,15 | 1,04 | 0,0026 | 0,0292 |
| ***FAM13A-AS1*** | 35,95 | 14,09 | 25,02 | 57,12 | 49,61 | 53,36 | 2,13 | 1,03 | 0,0038 | 0,0383 |
| ***TBC1D10C*** | 61,15 | 83,91 | 72,53 | 140,22 | 168,88 | 154,55 | 2,13 | 1,03 | 0,0000 | 0,0000 |
| ***SHPRH*** | 32,24 | 39,64 | 35,94 | 86,53 | 65,80 | 76,17 | 2,12 | 1,02 | 0,0006 | 0,0092 |
| ***SERINC1*** | 55,41 | 68,61 | 62,01 | 111,09 | 151,66 | 131,37 | 2,12 | 1,02 | 0,0000 | 0,0003 |
| ***STRBP*** | 30,39 | 47,49 | 38,94 | 88,82 | 75,83 | 82,32 | 2,11 | 1,02 | 0,0004 | 0,0063 |
| ***ZFYVE16*** | 24,09 | 26,76 | 25,43 | 55,98 | 51,15 | 53,56 | 2,11 | 1,01 | 0,0042 | 0,0415 |

| | | | | | | | | | | |
|---|---|---|---|---|---|---|---|---|---|---|
| ***NLRC3*** | 104,51 | 105,64 | 105,07 | 209,62 | 232,88 | 221,25 | 2,11 | 1,01 | 0,0000 | 0,0000 |
| ***SCPEP1*** | 48,55 | 34,41 | 41,48 | 106,24 | 68,37 | 87,31 | 2,10 | 1,01 | 0,0003 | 0,0049 |
| ***PLD3*** | 20,01 | 28,37 | 24,19 | 43,98 | 57,84 | 50,91 | 2,10 | 1,01 | 0,0054 | 0,0497 |
| ***TRAF5*** | 51,70 | 46,68 | 49,19 | 87,67 | 119,27 | 103,47 | 2,10 | 1,01 | 0,0001 | 0,0018 |
| ***CTSA*** | 59,85 | 70,22 | 65,04 | 134,80 | 138,80 | 136,80 | 2,10 | 1,01 | 0,0000 | 0,0002 |
| ***RCBTB2*** | 46,88 | 45,27 | 46,08 | 90,25 | 103,59 | 96,92 | 2,10 | 1,01 | 0,0001 | 0,0027 |
| ***ENTPD1*** | 158,99 | 152,12 | 155,55 | 348,41 | 305,11 | 326,76 | 2,10 | 1,01 | 0,0000 | 0,0000 |
| ***PIK3IP1*** | 72,27 | 79,68 | 75,97 | 141,94 | 176,59 | 159,26 | 2,10 | 1,01 | 0,0000 | 0,0001 |
| ***CLIP4*** | 31,13 | 28,37 | 29,75 | 62,26 | 62,46 | 62,36 | 2,10 | 1,01 | 0,0022 | 0,0256 |
| ***DDI2*** | 92,65 | 98,80 | 95,72 | 198,77 | 200,75 | 199,76 | 2,09 | 1,00 | 0,0000 | 0,0000 |
| ***SLC7A5*** | 31,69 | 36,62 | 34,15 | 60,54 | 81,74 | 71,14 | 2,08 | 1,00 | 0,0012 | 0,0158 |
| ***NKG7*** | 53,37 | 45,27 | 49,32 | 87,96 | 116,95 | 102,46 | 2,08 | 0,99 | 0,0001 | 0,0024 |
| ***PCED1B-AS1*** | 66,71 | 71,23 | 68,97 | 131,08 | 155,00 | 143,04 | 2,07 | 0,99 | 0,0000 | 0,0002 |
| ***CDK17*** | 45,58 | 40,24 | 42,91 | 72,25 | 105,65 | 88,95 | 2,07 | 0,99 | 0,0003 | 0,0056 |
| ***NFE2L3*** | 28,91 | 32,19 | 30,55 | 41,12 | 85,08 | 63,10 | 2,07 | 0,99 | 0,0025 | 0,0283 |
| ***SLC12A6*** | 57,63 | 74,65 | 66,14 | 102,53 | 170,16 | 136,34 | 2,06 | 0,98 | 0,0000 | 0,0003 |
| ***OXNAD1*** | 34,65 | 47,49 | 41,07 | 72,54 | 96,39 | 84,46 | 2,06 | 0,98 | 0,0005 | 0,0082 |
| ***SGSM2*** | 28,72 | 35,62 | 32,17 | 76,82 | 55,01 | 65,91 | 2,05 | 0,98 | 0,0022 | 0,0256 |
| ***HDLBP*** | 217,91 | 228,98 | 223,45 | 438,66 | 474,24 | 456,45 | 2,04 | 0,97 | 0,0000 | 0,0000 |
| ***INPP5F*** | 35,21 | 31,39 | 33,30 | 45,69 | 89,97 | 67,83 | 2,04 | 0,97 | 0,0021 | 0,0248 |
| ***SLFN12L*** | 196,42 | 245,08 | 220,75 | 377,83 | 521,03 | 449,43 | 2,04 | 0,97 | 0,0000 | 0,0000 |
| ***CARD19*** | 74,12 | 57,95 | 66,04 | 134,80 | 132,38 | 133,59 | 2,02 | 0,96 | 0,0000 | 0,0006 |
| ***NME3*** | 30,95 | 37,23 | 34,09 | 62,83 | 75,06 | 68,94 | 2,02 | 0,96 | 0,0021 | 0,0248 |
| ***ARHGAP45*** | 190,86 | 166,81 | 178,83 | 326,14 | 395,33 | 360,73 | 2,02 | 0,95 | 0,0000 | 0,0000 |
| ***CD96*** | 468,62 | 450,52 | 459,57 | 855,90 | 998,10 | 927,00 | 2,02 | 0,95 | 0,0000 | 0,0000 |
| ***RN7SL1*** | 83,20 | 84,91 | 84,06 | 157,36 | 181,22 | 169,29 | 2,01 | 0,95 | 0,0000 | 0,0001 |
| ***PDCD4*** | 160,10 | 165,40 | 162,75 | 331,56 | 322,59 | 327,08 | 2,01 | 0,95 | 0,0000 | 0,0000 |
| ***NOTCH1*** | 30,20 | 32,40 | 31,30 | 57,12 | 68,63 | 62,87 | 2,01 | 0,95 | 0,0036 | 0,0365 |
| ***PIK3R5*** | 45,95 | 67,61 | 56,78 | 117,09 | 110,27 | 113,68 | 2,00 | 0,94 | 0,0001 | 0,0023 |
| ***LINC02195*** | 75,42 | 50,51 | 62,96 | 39,41 | 23,91 | 31,66 | 0,50 | -1,05 | 0,0011 | 0,0152 |
| ***GSTO1*** | 216,80 | 237,63 | 227,22 | 134,51 | 93,82 | 114,17 | 0,50 | -1,05 | 0,0000 | 0,0000 |
| ***NDUFB5*** | 109,14 | 109,46 | 109,30 | 64,54 | 45,24 | 54,89 | 0,50 | -1,05 | 0,0000 | 0,0005 |
| ***NT5C3A*** | 51,70 | 45,27 | 48,49 | 22,85 | 25,70 | 24,28 | 0,50 | -1,06 | 0,0040 | 0,0401 |
| ***AURKB*** | 113,59 | 85,52 | 99,55 | 53,12 | 46,53 | 49,82 | 0,50 | -1,06 | 0,0000 | 0,0010 |
| ***CD74*** | 572,39 | 611,09 | 591,74 | 331,85 | 259,10 | 295,47 | 0,50 | -1,06 | 0,0000 | 0,0000 |
| ***CKAP2L*** | 68,19 | 71,83 | 70,01 | 35,13 | 34,70 | 34,91 | 0,50 | -1,06 | 0,0005 | 0,0083 |
| ***APEX1*** | 108,96 | 77,87 | 93,41 | 47,98 | 44,98 | 46,48 | 0,50 | -1,07 | 0,0001 | 0,0014 |
| ***KIF15*** | 70,97 | 55,54 | 63,25 | 30,56 | 32,39 | 31,47 | 0,50 | -1,07 | 0,0009 | 0,0132 |
| ***SLC25A24*** | 59,67 | 41,65 | 50,66 | 27,99 | 22,36 | 25,18 | 0,50 | -1,07 | 0,0030 | 0,0324 |
| ***CHST2*** | 55,22 | 59,36 | 57,29 | 34,56 | 22,36 | 28,46 | 0,50 | -1,07 | 0,0016 | 0,0203 |
| ***KIF2C*** | 77,27 | 59,16 | 68,21 | 31,99 | 35,73 | 33,86 | 0,50 | -1,07 | 0,0006 | 0,0088 |
| ***SMIM14*** | 80,42 | 70,63 | 75,52 | 34,56 | 40,36 | 37,46 | 0,50 | -1,07 | 0,0003 | 0,0051 |
| ***CSRNP1*** | 52,81 | 44,87 | 48,84 | 26,56 | 21,85 | 24,20 | 0,50 | -1,07 | 0,0035 | 0,0363 |
| ***RRM2*** | 256,27 | 229,99 | 243,13 | 119,37 | 121,58 | 120,48 | 0,50 | -1,07 | 0,0000 | 0,0000 |
| ***PARP9*** | 157,69 | 137,23 | 147,46 | 70,54 | 75,31 | 72,93 | 0,49 | -1,08 | 0,0000 | 0,0000 |
| ***ESD*** | 119,33 | 95,38 | 107,35 | 59,69 | 46,27 | 52,98 | 0,49 | -1,08 | 0,0000 | 0,0004 |
| ***TMA16*** | 61,52 | 55,74 | 58,63 | 30,84 | 26,73 | 28,79 | 0,49 | -1,08 | 0,0012 | 0,0162 |

| | | | | | | | | | | |
|---|---|---|---|---|---|---|---|---|---|---|
| ***MIR222HG*** | 109,70 | 78,07 | 93,88 | 37,13 | 55,01 | 46,07 | 0,49 | -1,09 | 0,0000 | 0,0011 |
| ***CEP55*** | 53,18 | 61,37 | 57,28 | 34,56 | 21,59 | 28,07 | 0,49 | -1,09 | 0,0014 | 0,0178 |
| ***CD38*** | 57,81 | 48,09 | 52,95 | 29,99 | 21,85 | 25,92 | 0,49 | -1,09 | 0,0020 | 0,0243 |
| ***RAN*** | 519,39 | 490,76 | 505,08 | 287,87 | 204,61 | 246,24 | 0,49 | -1,10 | 0,0000 | 0,0000 |
| ***ACTG1*** | 5514,51 | 5265,97 | 5390,24 | 2699,06 | 2527,24 | 2613,15 | 0,48 | -1,10 | 0,0000 | 0,0000 |
| ***AK2*** | 73,01 | 68,01 | 70,51 | 44,27 | 23,91 | 34,09 | 0,48 | -1,11 | 0,0003 | 0,0054 |
| ***EGR1*** | 145,28 | 141,05 | 143,16 | 69,97 | 67,35 | 68,66 | 0,48 | -1,12 | 0,0000 | 0,0000 |
| ***AURKA*** | 62,08 | 53,72 | 57,90 | 36,56 | 18,76 | 27,66 | 0,48 | -1,12 | 0,0009 | 0,0130 |
| ***DUT*** | 342,99 | 315,50 | 329,25 | 166,21 | 148,31 | 157,26 | 0,48 | -1,13 | 0,0000 | 0,0000 |
| ***PDE4B*** | 87,46 | 91,75 | 89,61 | 43,12 | 42,41 | 42,77 | 0,48 | -1,13 | 0,0000 | 0,0009 |
| ***MTHFD2*** | 224,03 | 183,51 | 203,77 | 90,53 | 103,07 | 96,80 | 0,48 | -1,13 | 0,0000 | 0,0000 |
| ***DTD1*** | 40,21 | 52,72 | 46,46 | 17,99 | 25,96 | 21,98 | 0,47 | -1,14 | 0,0027 | 0,0302 |
| ***INSIG1*** | 140,64 | 114,69 | 127,67 | 51,69 | 68,12 | 59,90 | 0,47 | -1,15 | 0,0000 | 0,0000 |
| ***FOS*** | 80,79 | 52,92 | 66,86 | 19,71 | 42,93 | 31,32 | 0,47 | -1,15 | 0,0003 | 0,0048 |
| ***SAMD9L*** | 259,79 | 321,34 | 290,56 | 140,51 | 129,55 | 135,03 | 0,46 | -1,16 | 0,0000 | 0,0000 |
| ***ADI1*** | 80,98 | 77,47 | 79,22 | 31,70 | 41,90 | 36,80 | 0,46 | -1,17 | 0,0001 | 0,0015 |
| ***GAB2*** | 49,48 | 43,46 | 46,47 | 20,28 | 22,88 | 21,58 | 0,46 | -1,17 | 0,0022 | 0,0261 |
| ***HADHB*** | 70,41 | 55,74 | 63,08 | 35,13 | 23,39 | 29,26 | 0,46 | -1,17 | 0,0004 | 0,0060 |
| ***POLD2*** | 44,10 | 44,07 | 44,08 | 24,85 | 15,94 | 20,39 | 0,46 | -1,17 | 0,0028 | 0,0310 |
| ***DIAPH3*** | 55,59 | 50,71 | 53,15 | 18,28 | 30,85 | 24,56 | 0,46 | -1,17 | 0,0010 | 0,0138 |
| ***HERC6*** | 42,25 | 70,83 | 56,54 | 19,71 | 31,87 | 25,79 | 0,46 | -1,19 | 0,0006 | 0,0090 |
| ***BIRC3*** | 1483,14 | 1559,81 | 1521,47 | 703,11 | 682,45 | 692,78 | 0,46 | -1,19 | 0,0000 | 0,0000 |
| ***MDH1*** | 141,75 | 130,59 | 136,17 | 76,25 | 47,30 | 61,77 | 0,45 | -1,20 | 0,0000 | 0,0000 |
| ***MRPS33*** | 73,56 | 87,33 | 80,45 | 47,12 | 25,70 | 36,41 | 0,45 | -1,20 | 0,0000 | 0,0009 |
| ***FH*** | 47,81 | 44,87 | 46,34 | 25,70 | 16,19 | 20,95 | 0,45 | -1,20 | 0,0017 | 0,0213 |
| ***ITGB3BP*** | 63,56 | 58,96 | 61,26 | 27,42 | 27,76 | 27,59 | 0,45 | -1,21 | 0,0003 | 0,0050 |
| ***CMTM7*** | 56,52 | 44,07 | 50,29 | 10,57 | 34,70 | 22,63 | 0,45 | -1,21 | 0,0010 | 0,0139 |
| ***BCL2L1*** | 73,56 | 70,02 | 71,79 | 36,27 | 28,02 | 32,14 | 0,45 | -1,22 | 0,0001 | 0,0017 |
| ***GXYLT1*** | 39,10 | 47,29 | 43,19 | 21,70 | 16,97 | 19,33 | 0,45 | -1,22 | 0,0023 | 0,0264 |
| ***MB21D2*** | 78,20 | 47,49 | 62,84 | 30,27 | 25,96 | 28,12 | 0,45 | -1,22 | 0,0002 | 0,0040 |
| ***ALDH6A1*** | 53,55 | 62,58 | 58,06 | 24,56 | 27,25 | 25,90 | 0,45 | -1,22 | 0,0004 | 0,0061 |
| ***TCF19*** | 59,67 | 56,54 | 58,10 | 26,56 | 25,19 | 25,87 | 0,45 | -1,23 | 0,0004 | 0,0059 |
| ***MSMO1*** | 47,07 | 43,87 | 45,47 | 25,42 | 14,91 | 20,16 | 0,44 | -1,23 | 0,0016 | 0,0198 |
| ***GNA15*** | 76,34 | 103,02 | 89,68 | 43,69 | 35,73 | 39,71 | 0,44 | -1,23 | 0,0000 | 0,0003 |
| ***ITGA4*** | 74,12 | 63,99 | 69,05 | 32,84 | 28,28 | 30,56 | 0,44 | -1,23 | 0,0001 | 0,0019 |
| ***ALYREF*** | 34,84 | 45,68 | 40,26 | 11,71 | 23,91 | 17,81 | 0,44 | -1,23 | 0,0029 | 0,0316 |
| ***LGALS3BP*** | 46,51 | 46,08 | 46,29 | 23,42 | 17,48 | 20,45 | 0,44 | -1,24 | 0,0013 | 0,0175 |
| ***CHMP5*** | 122,30 | 134,21 | 128,25 | 53,40 | 59,38 | 56,39 | 0,44 | -1,24 | 0,0000 | 0,0000 |
| ***ACAT1*** | 63,37 | 72,84 | 68,11 | 40,27 | 19,54 | 29,90 | 0,44 | -1,25 | 0,0001 | 0,0019 |
| ***DPP4*** | 87,28 | 104,03 | 95,65 | 51,41 | 32,13 | 41,77 | 0,44 | -1,25 | 0,0000 | 0,0001 |
| ***CCNA2*** | 48,36 | 61,77 | 55,07 | 30,84 | 17,22 | 24,03 | 0,44 | -1,25 | 0,0004 | 0,0065 |
| ***ABRACL*** | 174,74 | 154,53 | 164,63 | 72,25 | 70,69 | 71,47 | 0,43 | -1,26 | 0,0000 | 0,0000 |
| ***TUBA1C*** | 163,25 | 149,50 | 156,38 | 77,68 | 57,84 | 67,76 | 0,43 | -1,27 | 0,0000 | 0,0000 |
| ***TMEM14A*** | 42,25 | 54,13 | 48,19 | 16,28 | 25,19 | 20,73 | 0,43 | -1,27 | 0,0008 | 0,0114 |
| ***CAVIN3*** | 58,00 | 55,94 | 56,97 | 21,99 | 26,99 | 24,49 | 0,43 | -1,28 | 0,0003 | 0,0045 |
| ***FOXP3*** | 179,93 | 140,45 | 160,19 | 83,68 | 53,98 | 68,83 | 0,43 | -1,28 | 0,0000 | 0,0000 |
| ***ADAM19*** | 342,99 | 339,25 | 341,12 | 180,20 | 112,59 | 146,39 | 0,43 | -1,28 | 0,0000 | 0,0000 |
| ***TYMS*** | 182,34 | 197,79 | 190,06 | 77,97 | 84,31 | 81,14 | 0,43 | -1,29 | 0,0000 | 0,0000 |

| | | | | | | | | | | |
|---|---|---|---|---|---|---|---|---|---|---|
| ***ANKRD33B*** | 33,73 | 37,43 | 35,58 | 20,85 | 9,51 | 15,18 | 0,43 | -1,29 | 0,0039 | 0,0395 |
| ***CLIC4*** | 34,10 | 42,26 | 38,18 | 19,42 | 13,11 | 16,26 | 0,43 | -1,29 | 0,0027 | 0,0301 |
| ***SOX4*** | 569,80 | 561,19 | 565,49 | 209,33 | 271,69 | 240,51 | 0,43 | -1,29 | 0,0000 | 0,0000 |
| ***BEX5*** | 32,43 | 36,42 | 34,42 | 19,42 | 9,77 | 14,59 | 0,42 | -1,30 | 0,0044 | 0,0425 |
| ***PLPP1*** | 81,53 | 73,44 | 77,49 | 28,56 | 37,01 | 32,79 | 0,42 | -1,30 | 0,0000 | 0,0004 |
| ***CDT1*** | 68,38 | 63,99 | 66,18 | 18,28 | 37,27 | 27,77 | 0,42 | -1,31 | 0,0001 | 0,0012 |
| ***LGALS1*** | 1384,19 | 1243,90 | 1314,05 | 597,16 | 504,32 | 550,74 | 0,42 | -1,31 | 0,0000 | 0,0000 |
| ***TMSB10*** | 2708,15 | 2775,95 | 2742,05 | 1188,89 | 1108,88 | 1148,89 | 0,42 | -1,31 | 0,0000 | 0,0000 |
| ***GNG4*** | 44,29 | 44,47 | 44,38 | 12,00 | 25,19 | 18,59 | 0,42 | -1,31 | 0,0010 | 0,0135 |
| ***NRP2*** | 74,68 | 86,72 | 80,70 | 38,55 | 29,05 | 33,80 | 0,42 | -1,31 | 0,0000 | 0,0002 |
| ***MXD1*** | 58,00 | 42,66 | 50,33 | 17,99 | 24,16 | 21,08 | 0,42 | -1,31 | 0,0004 | 0,0069 |
| ***GNG8*** | 295,92 | 263,99 | 279,96 | 118,23 | 116,18 | 117,21 | 0,42 | -1,32 | 0,0000 | 0,0000 |
| ***CCNB1*** | 47,81 | 53,93 | 50,87 | 12,85 | 29,56 | 21,21 | 0,42 | -1,32 | 0,0004 | 0,0062 |
| ***PRDX2*** | 52,44 | 49,90 | 51,17 | 15,42 | 26,99 | 21,21 | 0,41 | -1,33 | 0,0003 | 0,0057 |
| ***KIF4A*** | 32,06 | 34,41 | 33,23 | 15,71 | 11,82 | 13,77 | 0,41 | -1,33 | 0,0043 | 0,0420 |
| ***ECT2*** | 36,50 | 42,46 | 39,48 | 26,27 | 6,17 | 16,22 | 0,41 | -1,34 | 0,0016 | 0,0204 |
| ***BMAL2*** | 40,58 | 39,04 | 39,81 | 16,56 | 15,94 | 16,25 | 0,41 | -1,35 | 0,0014 | 0,0186 |
| ***LST1*** | 73,75 | 70,22 | 71,99 | 27,13 | 31,62 | 29,37 | 0,41 | -1,35 | 0,0000 | 0,0004 |
| ***PKD2*** | 42,62 | 50,51 | 46,56 | 32,27 | 5,66 | 18,96 | 0,41 | -1,35 | 0,0006 | 0,0086 |
| ***PCLAF*** | 263,31 | 246,08 | 254,70 | 94,24 | 112,84 | 103,54 | 0,41 | -1,36 | 0,0000 | 0,0000 |
| ***HLA-DRB5*** | 52,63 | 60,36 | 56,49 | 26,56 | 19,28 | 22,92 | 0,41 | -1,36 | 0,0001 | 0,0025 |
| ***CCR7*** | 164,18 | 193,77 | 178,97 | 71,40 | 73,77 | 72,58 | 0,41 | -1,36 | 0,0000 | 0,0000 |
| ***DENND5A*** | 29,83 | 39,44 | 34,64 | 10,85 | 17,22 | 14,04 | 0,41 | -1,36 | 0,0029 | 0,0317 |
| ***IGF1*** | 127,12 | 132,40 | 129,76 | 41,70 | 62,46 | 52,08 | 0,40 | -1,38 | 0,0000 | 0,0000 |
| ***ENTPD1-AS1*** | 25,76 | 49,30 | 37,53 | 15,71 | 14,39 | 15,05 | 0,40 | -1,37 | 0,0017 | 0,0215 |
| ***DPH3*** | 35,02 | 37,63 | 36,32 | 15,99 | 13,11 | 14,55 | 0,40 | -1,38 | 0,0020 | 0,0244 |
| ***PLSCR1*** | 72,45 | 85,92 | 79,19 | 31,13 | 31,87 | 31,50 | 0,40 | -1,39 | 0,0000 | 0,0001 |
| ***GNS*** | 53,92 | 53,12 | 53,52 | 30,27 | 12,08 | 21,18 | 0,40 | -1,40 | 0,0001 | 0,0027 |
| ***H2AC6*** | 60,59 | 53,72 | 57,16 | 31,41 | 13,62 | 22,52 | 0,39 | -1,40 | 0,0001 | 0,0016 |
| ***MCM2*** | 92,09 | 75,25 | 83,67 | 30,84 | 34,96 | 32,90 | 0,39 | -1,41 | 0,0000 | 0,0001 |
| ***ASF1B*** | 61,89 | 50,91 | 56,40 | 23,99 | 20,31 | 22,15 | 0,39 | -1,41 | 0,0001 | 0,0017 |
| ***MT2A*** | 707,84 | 730,00 | 718,92 | 321,28 | 242,39 | 281,84 | 0,39 | -1,41 | 0,0000 | 0,0000 |
| ***OAS2*** | 243,30 | 221,94 | 232,62 | 79,68 | 102,56 | 91,12 | 0,39 | -1,41 | 0,0000 | 0,0000 |
| ***TNFSF4*** | 146,39 | 104,43 | 125,41 | 45,69 | 51,92 | 48,81 | 0,39 | -1,42 | 0,0000 | 0,0000 |
| ***HERC5*** | 201,61 | 185,72 | 193,66 | 68,26 | 81,74 | 75,00 | 0,39 | -1,43 | 0,0000 | 0,0000 |
| ***HIVEP1*** | 36,13 | 57,35 | 46,74 | 19,42 | 16,71 | 18,06 | 0,39 | -1,43 | 0,0003 | 0,0049 |
| ***LAP3*** | 66,52 | 47,29 | 56,90 | 25,42 | 18,51 | 21,96 | 0,39 | -1,43 | 0,0001 | 0,0013 |
| ***RNU5F-1*** | 31,32 | 31,99 | 31,65 | 12,57 | 11,82 | 12,20 | 0,39 | -1,43 | 0,0031 | 0,0332 |
| ***UBE2Q2*** | 84,31 | 57,55 | 70,93 | 17,99 | 36,50 | 27,25 | 0,38 | -1,44 | 0,0000 | 0,0002 |
| ***IRF7*** | 121,93 | 83,10 | 102,51 | 41,41 | 37,27 | 39,34 | 0,38 | -1,44 | 0,0000 | 0,0000 |
| ***GRINA*** | 64,48 | 71,63 | 68,06 | 31,70 | 20,05 | 25,87 | 0,38 | -1,45 | 0,0000 | 0,0002 |
| ***SOCS4*** | 45,03 | 39,64 | 42,33 | 17,71 | 14,39 | 16,05 | 0,38 | -1,46 | 0,0005 | 0,0073 |
| ***TSHZ2*** | 81,72 | 63,79 | 72,75 | 21,13 | 33,42 | 27,27 | 0,37 | -1,47 | 0,0000 | 0,0001 |
| ***PTPN14*** | 46,70 | 45,07 | 45,88 | 11,71 | 22,62 | 17,16 | 0,37 | -1,48 | 0,0002 | 0,0040 |
| ***FAM111B*** | 53,74 | 56,14 | 54,94 | 16,28 | 24,68 | 20,48 | 0,37 | -1,48 | 0,0000 | 0,0011 |
| ***PRDX4*** | 35,58 | 21,53 | 28,55 | 9,42 | 11,82 | 10,62 | 0,37 | -1,48 | 0,0041 | 0,0403 |
| ***RAD51AP1*** | 26,13 | 32,60 | 29,36 | 10,00 | 11,82 | 10,91 | 0,37 | -1,48 | 0,0035 | 0,0363 |

| | | | | | | | | | | |
|---|---|---|---|---|---|---|---|---|---|---|
| ***ELAPOR1*** | 29,83 | 33,60 | 31,72 | 14,57 | 9,00 | 11,78 | 0,37 | -1,49 | 0,0023 | 0,0267 |
| ***PHGDH*** | 41,88 | 50,71 | 46,29 | 13,14 | 21,08 | 17,11 | 0,37 | -1,49 | 0,0002 | 0,0034 |
| ***PDZD11*** | 22,05 | 33,60 | 27,83 | 10,28 | 10,28 | 10,28 | 0,37 | -1,49 | 0,0045 | 0,0430 |
| ***CORO1B*** | 31,69 | 47,69 | 39,69 | 19,42 | 9,77 | 14,59 | 0,37 | -1,50 | 0,0005 | 0,0083 |
| ***FZD3*** | 40,03 | 26,76 | 33,39 | 13,99 | 10,54 | 12,27 | 0,37 | -1,50 | 0,0016 | 0,0198 |
| ***GZMB*** | 1929,34 | 1682,55 | 1805,94 | 648,85 | 675,51 | 662,18 | 0,37 | -1,51 | 0,0000 | 0,0000 |
| ***TRAPPC13*** | 29,83 | 24,95 | 27,39 | 12,57 | 7,45 | 10,01 | 0,37 | -1,51 | 0,0044 | 0,0428 |
| ***MGAT2*** | 30,76 | 39,04 | 34,90 | 15,14 | 10,28 | 12,71 | 0,36 | -1,51 | 0,0011 | 0,0152 |
| ***RSAD2*** | 203,83 | 194,57 | 199,20 | 71,97 | 72,74 | 72,36 | 0,36 | -1,52 | 0,0000 | 0,0000 |
| ***SELL*** | 150,46 | 125,96 | 138,21 | 63,11 | 36,76 | 49,94 | 0,36 | -1,53 | 0,0000 | 0,0000 |
| ***CMSS1*** | 29,65 | 25,15 | 27,40 | 15,99 | 3,60 | 9,80 | 0,36 | -1,54 | 0,0038 | 0,0383 |
| ***DDX60*** | 120,07 | 116,10 | 118,09 | 39,41 | 44,98 | 42,20 | 0,36 | -1,54 | 0,0000 | 0,0000 |
| ***STARD4*** | 71,16 | 75,86 | 73,51 | 36,27 | 16,19 | 26,23 | 0,36 | -1,54 | 0,0000 | 0,0000 |
| ***CABLES1*** | 35,95 | 32,19 | 34,07 | 9,14 | 15,17 | 12,15 | 0,36 | -1,54 | 0,0011 | 0,0148 |
| ***PGRMC1*** | 31,50 | 30,18 | 30,84 | 13,42 | 8,48 | 10,95 | 0,36 | -1,55 | 0,0019 | 0,0230 |
| ***XAF1*** | 479,74 | 512,29 | 496,02 | 130,51 | 221,06 | 175,78 | 0,35 | -1,56 | 0,0000 | 0,0000 |
| ***MT-TC*** | 137,12 | 178,07 | 157,60 | 44,84 | 66,57 | 55,71 | 0,35 | -1,56 | 0,0000 | 0,0000 |
| ***TFRC*** | 433,97 | 401,02 | 417,50 | 153,36 | 141,63 | 147,49 | 0,35 | -1,56 | 0,0000 | 0,0000 |
| ***TRABD2A*** | 32,80 | 53,52 | 43,16 | 20,85 | 9,51 | 15,18 | 0,35 | -1,56 | 0,0002 | 0,0034 |
| ***UBE2C*** | 76,53 | 66,60 | 71,57 | 24,56 | 25,70 | 25,13 | 0,35 | -1,57 | 0,0000 | 0,0000 |
| ***CMPK2*** | 41,51 | 41,65 | 41,58 | 11,71 | 17,48 | 14,59 | 0,35 | -1,57 | 0,0002 | 0,0042 |
| ***BLVRA*** | 41,32 | 16,50 | 28,91 | 5,14 | 14,91 | 10,02 | 0,35 | -1,59 | 0,0023 | 0,0264 |
| ***SOCS1*** | 76,53 | 85,32 | 80,92 | 21,99 | 33,67 | 27,83 | 0,34 | -1,60 | 0,0000 | 0,0000 |
| ***CHRNA6*** | 75,05 | 65,60 | 70,32 | 33,41 | 14,91 | 24,16 | 0,34 | -1,60 | 0,0000 | 0,0000 |
| ***ZMYM5*** | 30,20 | 24,35 | 27,28 | 8,28 | 10,28 | 9,28 | 0,34 | -1,61 | 0,0028 | 0,0306 |
| ***CEACAM1*** | 66,89 | 74,25 | 70,57 | 22,56 | 25,19 | 23,88 | 0,34 | -1,62 | 0,0000 | 0,0000 |
| ***PPA1*** | 54,29 | 52,72 | 53,51 | 24,85 | 11,31 | 18,08 | 0,34 | -1,62 | 0,0000 | 0,0004 |
| ***DPP3*** | 43,55 | 37,43 | 40,49 | 18,56 | 8,74 | 13,65 | 0,34 | -1,63 | 0,0002 | 0,0035 |
| ***DUSP6*** | 142,31 | 163,18 | 152,75 | 64,83 | 38,04 | 51,44 | 0,34 | -1,63 | 0,0000 | 0,0000 |
| ***HLA-DPB1*** | 74,68 | 68,01 | 71,34 | 16,56 | 31,36 | 23,96 | 0,34 | -1,63 | 0,0000 | 0,0000 |
| ***SLC12A2-DT*** | 25,20 | 24,55 | 24,87 | 12,57 | 4,11 | 8,34 | 0,34 | -1,63 | 0,0041 | 0,0406 |
| ***ZNF282*** | 159,91 | 138,84 | 149,38 | 58,83 | 41,13 | 49,98 | 0,33 | -1,64 | 0,0000 | 0,0000 |
| ***CASC15*** | 27,98 | 27,16 | 27,57 | 7,14 | 11,31 | 9,23 | 0,33 | -1,64 | 0,0023 | 0,0269 |
| ***CCR4*** | 151,76 | 155,74 | 153,75 | 44,27 | 58,35 | 51,31 | 0,33 | -1,64 | 0,0000 | 0,0000 |
| ***APP*** | 76,53 | 65,60 | 71,06 | 16,56 | 30,85 | 23,70 | 0,33 | -1,64 | 0,0000 | 0,0000 |
| ***IER3*** | 162,14 | 154,13 | 158,13 | 47,12 | 58,35 | 52,74 | 0,33 | -1,64 | 0,0000 | 0,0000 |
| ***SERPINB6*** | 32,61 | 27,57 | 30,09 | 7,14 | 12,85 | 10,00 | 0,33 | -1,65 | 0,0013 | 0,0173 |
| ***AHCY*** | 31,13 | 28,98 | 30,05 | 12,00 | 7,97 | 9,98 | 0,33 | -1,65 | 0,0013 | 0,0173 |
| ***GMPPB*** | 32,43 | 22,54 | 27,48 | 10,00 | 8,23 | 9,11 | 0,33 | -1,65 | 0,0022 | 0,0259 |
| ***MX2*** | 55,78 | 42,66 | 49,22 | 11,14 | 21,34 | 16,24 | 0,33 | -1,66 | 0,0000 | 0,0006 |
| ***GINS2*** | 30,02 | 33,00 | 31,51 | 19,71 | 1,03 | 10,37 | 0,33 | -1,66 | 0,0013 | 0,0170 |
| ***CDKN2D*** | 77,27 | 83,10 | 80,19 | 35,13 | 17,22 | 26,17 | 0,33 | -1,67 | 0,0000 | 0,0000 |
| ***RCCD1*** | 25,02 | 26,96 | 25,99 | 10,00 | 6,94 | 8,47 | 0,33 | -1,67 | 0,0027 | 0,0301 |
| ***GPR108*** | 30,20 | 29,98 | 30,09 | 15,99 | 3,60 | 9,80 | 0,33 | -1,68 | 0,0011 | 0,0152 |
| ***SLC8A1-AS1*** | 53,00 | 33,00 | 43,00 | 16,56 | 10,80 | 13,68 | 0,32 | -1,71 | 0,0001 | 0,0013 |
| ***ACOT7*** | 55,96 | 40,85 | 48,40 | 6,85 | 23,91 | 15,38 | 0,32 | -1,71 | 0,0000 | 0,0005 |

| | | | | | | | | | | |
|---|---|---|---|---|---|---|---|---|---|---|
| ***WARS1*** | 182,71 | 177,67 | 180,19 | 59,97 | 54,49 | 57,23 | 0,32 | -1,71 | 0,0000 | 0,0000 |
| ***SLC35F3*** | 27,61 | 22,54 | 25,07 | 2,29 | 13,62 | 7,95 | 0,32 | -1,71 | 0,0028 | 0,0306 |
| ***PRKCD*** | 26,68 | 43,06 | 34,87 | 12,57 | 9,51 | 11,04 | 0,32 | -1,72 | 0,0003 | 0,0054 |
| ***RIOX2*** | 60,41 | 64,99 | 62,70 | 24,56 | 14,65 | 19,61 | 0,31 | -1,73 | 0,0000 | 0,0000 |
| ***PLAAT3*** | 50,40 | 59,56 | 54,98 | 16,56 | 17,22 | 16,89 | 0,31 | -1,76 | 0,0000 | 0,0001 |
| ***QSER1*** | 14,27 | 36,62 | 25,44 | 7,14 | 8,48 | 7,81 | 0,31 | -1,76 | 0,0021 | 0,0249 |
| ***CREM*** | 300,56 | 309,87 | 305,21 | 91,10 | 96,13 | 93,62 | 0,31 | -1,76 | 0,0000 | 0,0000 |
| ***AIF1*** | 30,39 | 26,56 | 28,47 | 16,56 | 0,77 | 8,67 | 0,30 | -1,77 | 0,0012 | 0,0158 |
| ***C4orf33*** | 26,50 | 30,18 | 28,34 | 12,00 | 5,14 | 8,57 | 0,30 | -1,78 | 0,0009 | 0,0133 |
| ***OAS3*** | 142,87 | 136,02 | 139,44 | 35,98 | 48,32 | 42,15 | 0,30 | -1,79 | 0,0000 | 0,0000 |
| ***F5*** | 45,40 | 41,25 | 43,32 | 10,00 | 16,19 | 13,09 | 0,30 | -1,78 | 0,0000 | 0,0007 |
| ***BASP1*** | 70,97 | 63,38 | 67,18 | 25,42 | 14,91 | 20,16 | 0,30 | -1,79 | 0,0000 | 0,0000 |
| ***EIF2AK2*** | 473,26 | 421,95 | 447,60 | 142,51 | 123,90 | 133,20 | 0,30 | -1,81 | 0,0000 | 0,0000 |
| ***MZB1*** | 23,72 | 43,06 | 33,39 | 14,85 | 4,88 | 9,87 | 0,30 | -1,82 | 0,0002 | 0,0044 |
| ***HLA-DRA*** | 111,37 | 138,03 | 124,70 | 47,41 | 25,96 | 36,68 | 0,29 | -1,82 | 0,0000 | 0,0000 |
| ***BPGM*** | 26,68 | 33,80 | 30,24 | 9,14 | 8,48 | 8,81 | 0,29 | -1,84 | 0,0005 | 0,0073 |
| ***NUDT19*** | 13,71 | 26,36 | 20,04 | 4,86 | 6,68 | 5,77 | 0,29 | -1,85 | 0,0054 | 0,0494 |
| ***IFT56*** | 27,61 | 24,95 | 26,28 | 7,14 | 7,97 | 7,55 | 0,29 | -1,85 | 0,0011 | 0,0148 |
| ***LIMA1*** | 111,92 | 138,64 | 125,28 | 34,56 | 37,01 | 35,79 | 0,29 | -1,87 | 0,0000 | 0,0000 |
| ***IFI44*** | 89,31 | 70,43 | 79,87 | 26,27 | 19,28 | 22,78 | 0,29 | -1,87 | 0,0000 | 0,0000 |
| ***IFIT3*** | 68,56 | 46,08 | 57,32 | 12,57 | 20,05 | 16,31 | 0,28 | -1,87 | 0,0000 | 0,0000 |
| ***FRMD4B*** | 84,13 | 78,47 | 81,30 | 23,70 | 22,36 | 23,03 | 0,28 | -1,88 | 0,0000 | 0,0000 |
| ***CD40LG*** | 120,82 | 141,45 | 131,13 | 22,28 | 51,92 | 37,10 | 0,28 | -1,88 | 0,0000 | 0,0000 |
| ***LINC01806*** | 27,05 | 16,90 | 21,98 | 11,14 | 1,29 | 6,21 | 0,28 | -1,88 | 0,0029 | 0,0312 |
| ***TRAV6*** | 31,13 | 24,95 | 28,04 | 1,43 | 14,39 | 7,91 | 0,28 | -1,88 | 0,0006 | 0,0094 |
| ***TNF*** | 78,01 | 58,96 | 68,48 | 14,28 | 24,16 | 19,22 | 0,28 | -1,89 | 0,0000 | 0,0000 |
| ***SESN3*** | 303,15 | 271,64 | 287,39 | 68,26 | 90,74 | 79,50 | 0,28 | -1,91 | 0,0000 | 0,0000 |
| ***TTLL7*** | 59,30 | 64,59 | 61,94 | 15,42 | 18,76 | 17,09 | 0,28 | -1,92 | 0,0000 | 0,0000 |
| ***CHDH*** | 50,96 | 43,06 | 47,01 | 12,00 | 13,88 | 12,94 | 0,28 | -1,92 | 0,0000 | 0,0001 |
| ***HLA-DPA1*** | 88,02 | 88,74 | 88,38 | 25,99 | 22,62 | 24,30 | 0,28 | -1,92 | 0,0000 | 0,0000 |
| ***NEDD4L*** | 18,16 | 22,13 | 20,15 | 5,14 | 5,91 | 5,53 | 0,27 | -1,92 | 0,0040 | 0,0400 |
| ***EPSTI1*** | 94,50 | 112,28 | 103,39 | 43,41 | 13,11 | 28,26 | 0,27 | -1,93 | 0,0000 | 0,0000 |
| ***LGALS9*** | 123,22 | 152,12 | 137,67 | 25,13 | 50,12 | 37,63 | 0,27 | -1,93 | 0,0000 | 0,0000 |
| ***SCFD2*** | 22,98 | 21,53 | 22,25 | 7,71 | 4,37 | 6,04 | 0,27 | -1,94 | 0,0021 | 0,0252 |
| ***LY6E*** | 166,77 | 145,48 | 156,12 | 45,69 | 38,81 | 42,25 | 0,27 | -1,94 | 0,0000 | 0,0000 |
| ***UQCC3*** | 23,16 | 18,51 | 20,84 | 6,85 | 4,37 | 5,61 | 0,27 | -1,95 | 0,0030 | 0,0321 |
| ***LYRM4*** | 42,80 | 42,86 | 42,83 | 9,42 | 13,62 | 11,52 | 0,27 | -1,95 | 0,0000 | 0,0002 |
| ***PTGIR*** | 137,49 | 155,14 | 146,31 | 28,56 | 50,12 | 39,34 | 0,27 | -1,95 | 0,0000 | 0,0000 |
| ***NCDN*** | 29,83 | 39,84 | 34,84 | 11,14 | 7,45 | 9,30 | 0,27 | -1,96 | 0,0001 | 0,0014 |
| ***DDIT4*** | 261,64 | 262,58 | 262,11 | 61,97 | 77,63 | 69,80 | 0,27 | -1,97 | 0,0000 | 0,0000 |
| ***CDC7*** | 23,90 | 20,32 | 22,11 | 3,14 | 8,48 | 5,81 | 0,26 | -1,98 | 0,0019 | 0,0225 |
| ***USP18*** | 18,16 | 19,52 | 18,84 | 6,28 | 3,60 | 4,94 | 0,26 | -1,99 | 0,0046 | 0,0437 |
| ***TNFRSF4*** | 58,74 | 66,20 | 62,47 | 22,56 | 10,03 | 16,29 | 0,26 | -2,00 | 0,0000 | 0,0000 |
| ***DEPDC1*** | 21,68 | 31,19 | 26,43 | 6,28 | 7,45 | 6,87 | 0,26 | -2,00 | 0,0005 | 0,0081 |
| ***CYFIP1*** | 14,45 | 23,74 | 19,10 | 7,14 | 2,57 | 4,86 | 0,25 | -2,03 | 0,0037 | 0,0371 |
| ***ATL1*** | 20,38 | 19,12 | 19,75 | 2,57 | 7,45 | 5,01 | 0,25 | -2,03 | 0,0030 | 0,0323 |
| ***LHX9*** | 18,53 | 21,93 | 20,23 | 2,29 | 7,97 | 5,13 | 0,25 | -2,03 | 0,0026 | 0,0292 |
| ***PLA2G12A*** | 17,60 | 18,71 | 18,16 | 2,57 | 6,43 | 4,50 | 0,25 | -2,06 | 0,0043 | 0,0419 |

| | | | | | | | | | | |
|---|---|---|---|---|---|---|---|---|---|---|
| ***C6orf120*** | 32,06 | 21,53 | 26,79 | 2,29 | 10,80 | 6,54 | 0,24 | -2,09 | 0,0003 | 0,0051 |
| ***IL2RA*** | 1070,11 | 1163,62 | 1116,86 | 258,17 | 278,63 | 268,40 | 0,24 | -2,12 | 0,0000 | 0,0000 |
| ***TLR5*** | 25,39 | 18,71 | 22,05 | 5,71 | 4,88 | 5,30 | 0,24 | -2,11 | 0,0011 | 0,0151 |
| ***IFITM2*** | 168,99 | 208,26 | 188,62 | 55,12 | 34,19 | 44,65 | 0,24 | -2,14 | 0,0000 | 0,0000 |
| ***LAIR2*** | 66,71 | 89,94 | 78,33 | 31,41 | 5,14 | 18,28 | 0,23 | -2,16 | 0,0000 | 0,0000 |
| ***NET1*** | 27,61 | 22,94 | 25,27 | 9,71 | 2,06 | 5,88 | 0,23 | -2,16 | 0,0003 | 0,0056 |
| ***NFKBIZ*** | 274,24 | 330,39 | 302,32 | 54,55 | 84,82 | 69,69 | 0,23 | -2,18 | 0,0000 | 0,0000 |
| ***PDE7B*** | 188,08 | 210,67 | 199,38 | 51,69 | 38,81 | 45,25 | 0,23 | -2,20 | 0,0000 | 0,0000 |
| ***SOCS2*** | 43,36 | 56,14 | 49,75 | 7,14 | 15,42 | 11,28 | 0,23 | -2,20 | 0,0000 | 0,0000 |
| ***F11R*** | 17,97 | 16,90 | 17,44 | 2,00 | 5,66 | 3,83 | 0,22 | -2,23 | 0,0031 | 0,0331 |
| ***PMAIP1*** | 64,67 | 70,83 | 67,75 | 21,70 | 7,97 | 14,84 | 0,22 | -2,25 | 0,0000 | 0,0000 |
| ***CCDC50*** | 64,30 | 69,42 | 66,86 | 17,99 | 11,05 | 14,52 | 0,22 | -2,26 | 0,0000 | 0,0000 |
| ***EVI5*** | 66,52 | 62,18 | 64,35 | 19,42 | 7,97 | 13,69 | 0,21 | -2,29 | 0,0000 | 0,0000 |
| ***H2BC5*** | 25,94 | 24,15 | 25,04 | 8,57 | 2,06 | 5,31 | 0,21 | -2,29 | 0,0002 | 0,0035 |
| ***OAS1*** | 92,46 | 71,43 | 81,95 | 14,57 | 19,54 | 17,05 | 0,21 | -2,32 | 0,0000 | 0,0000 |
| ***ADCY1*** | 49,48 | 42,26 | 45,87 | 8,85 | 9,77 | 9,31 | 0,20 | -2,36 | 0,0000 | 0,0000 |
| ***NEK2*** | 24,27 | 32,80 | 28,54 | 7,14 | 4,37 | 5,76 | 0,20 | -2,36 | 0,0000 | 0,0009 |
| ***PHLDA2*** | 84,50 | 72,24 | 78,37 | 17,14 | 14,39 | 15,76 | 0,20 | -2,37 | 0,0000 | 0,0000 |
| ***PPM1L*** | 20,20 | 17,91 | 19,05 | 4,57 | 3,09 | 3,83 | 0,20 | -2,37 | 0,0011 | 0,0154 |
| ***HSD3B7*** | 16,12 | 25,76 | 20,94 | 3,43 | 4,88 | 4,16 | 0,20 | -2,38 | 0,0005 | 0,0085 |
| ***HLA-DQA1*** | 16,49 | 17,10 | 16,80 | 2,29 | 4,37 | 3,33 | 0,20 | -2,38 | 0,0025 | 0,0281 |
| ***MCOLN3*** | 23,90 | 25,35 | 24,63 | 1,43 | 8,23 | 4,83 | 0,20 | -2,40 | 0,0001 | 0,0026 |
| ***ISG15*** | 253,86 | 240,65 | 247,26 | 56,55 | 38,04 | 47,29 | 0,19 | -2,44 | 0,0000 | 0,0000 |
| ***PMCH*** | 447,68 | 446,09 | 446,89 | 45,41 | 107,70 | 76,55 | 0,17 | -2,61 | 0,0000 | 0,0000 |
| ***DSCC1*** | 20,75 | 12,48 | 16,61 | 4,86 | 0,77 | 2,81 | 0,17 | -2,62 | 0,0013 | 0,0171 |
| ***ACP5*** | 21,87 | 14,49 | 18,18 | 2,29 | 3,86 | 3,07 | 0,17 | -2,61 | 0,0007 | 0,0102 |
| ***LIF*** | 49,85 | 40,24 | 45,04 | 4,00 | 11,05 | 7,53 | 0,17 | -2,64 | 0,0000 | 0,0000 |
| ***ARG2*** | 34,84 | 24,35 | 29,59 | 1,71 | 7,97 | 4,84 | 0,16 | -2,66 | 0,0000 | 0,0001 |
| ***DNAJC12*** | 69,67 | 50,51 | 60,09 | 11,14 | 8,48 | 9,81 | 0,16 | -2,67 | 0,0000 | 0,0000 |
| ***PTGS2*** | 130,82 | 133,00 | 131,91 | 12,00 | 30,07 | 21,03 | 0,16 | -2,71 | 0,0000 | 0,0000 |
| ***ST8SIA1*** | 23,72 | 16,90 | 20,31 | 4,28 | 2,06 | 3,17 | 0,16 | -2,73 | 0,0002 | 0,0034 |
| ***IL13*** | 122,85 | 145,08 | 133,96 | 11,14 | 30,33 | 20,73 | 0,15 | -2,75 | 0,0000 | 0,0000 |
| ***TSPAN2*** | 19,83 | 16,50 | 18,16 | 2,00 | 3,60 | 2,80 | 0,15 | -2,74 | 0,0004 | 0,0072 |
| ***NTAQ1*** | 20,57 | 13,88 | 17,23 | 3,14 | 2,06 | 2,60 | 0,15 | -2,78 | 0,0006 | 0,0093 |
| ***IFITM1*** | 16,49 | 22,54 | 19,51 | 2,00 | 3,86 | 2,93 | 0,15 | -2,78 | 0,0002 | 0,0039 |
| ***NEIL3*** | 19,46 | 9,46 | 14,46 | 3,43 | 0,77 | 2,10 | 0,15 | -2,84 | 0,0018 | 0,0225 |
| ***ZBED2*** | 87,65 | 68,01 | 77,83 | 14,57 | 7,71 | 11,14 | 0,14 | -2,86 | 0,0000 | 0,0000 |
| ***P4HA2*** | 11,67 | 14,49 | 13,08 | 3,71 | 0,00 | 1,86 | 0,14 | -2,88 | 0,0033 | 0,0344 |
| ***SIGLEC17P*** | 36,32 | 33,80 | 35,06 | 8,28 | 1,54 | 4,91 | 0,14 | -2,89 | 0,0000 | 0,0000 |
| ***ACVR2A*** | 22,79 | 16,90 | 19,85 | 4,00 | 1,54 | 2,77 | 0,14 | -2,89 | 0,0001 | 0,0024 |
| ***OSBPL6*** | 13,53 | 15,29 | 14,41 | 4,00 | 0,00 | 2,00 | 0,14 | -2,91 | 0,0016 | 0,0202 |
| ***ADGRE1*** | 16,12 | 9,86 | 12,99 | 2,57 | 1,03 | 1,80 | 0,14 | -2,90 | 0,0032 | 0,0335 |
| ***IL17RB*** | 89,31 | 95,38 | 92,34 | 8,00 | 17,48 | 12,74 | 0,14 | -2,91 | 0,0000 | 0,0000 |
| ***ACTN1*** | 159,17 | 153,73 | 156,45 | 24,85 | 17,74 | 21,29 | 0,14 | -2,94 | 0,0000 | 0,0000 |
| ***ARL4A*** | 19,46 | 22,54 | 21,00 | 5,71 | 0,00 | 2,86 | 0,14 | -2,94 | 0,0001 | 0,0014 |
| ***IRX3*** | 51,33 | 36,82 | 44,08 | 7,71 | 3,86 | 5,78 | 0,13 | -2,99 | 0,0000 | 0,0000 |
| ***HMHB1*** | 10,01 | 20,73 | 15,37 | 0,57 | 3,34 | 1,96 | 0,13 | -3,00 | 0,0008 | 0,0112 |

| | | | | | | | | | | |
|---|---|---|---|---|---|---|---|---|---|---|
| ***SCD*** | 30,95 | 16,50 | 23,72 | 2,86 | 3,09 | 2,97 | 0,13 | -3,05 | 0,0000 | 0,0003 |
| ***MT1E*** | 103,03 | 143,26 | 123,15 | 14,28 | 16,19 | 15,24 | 0,12 | -3,07 | 0,0000 | 0,0000 |
| ***IL1RN*** | 21,50 | 16,10 | 18,80 | 2,00 | 2,57 | 2,28 | 0,12 | -3,08 | 0,0001 | 0,0021 |
| ***CYSLTR1*** | 158,99 | 165,60 | 162,29 | 21,99 | 16,71 | 19,35 | 0,12 | -3,13 | 0,0000 | 0,0000 |
| ***WFDC1*** | 16,12 | 7,85 | 11,98 | 0,29 | 2,57 | 1,43 | 0,12 | -3,09 | 0,0034 | 0,0356 |
| ***FN1*** | 74,68 | 53,72 | 64,20 | 6,28 | 9,00 | 7,64 | 0,12 | -3,13 | 0,0000 | 0,0000 |
| ***CD4*** | 88,02 | 89,74 | 88,88 | 10,57 | 9,25 | 9,91 | 0,11 | -3,22 | 0,0000 | 0,0000 |
| ***RHOBTB3*** | 17,42 | 16,30 | 16,86 | 2,00 | 1,29 | 1,64 | 0,10 | -3,40 | 0,0001 | 0,0022 |
| ***HLF*** | 30,57 | 41,85 | 36,21 | 1,43 | 5,14 | 3,28 | 0,09 | -3,50 | 0,0000 | 0,0000 |
| ***FSD1L*** | 6,86 | 13,28 | 10,07 | 1,71 | 0,00 | 0,86 | 0,09 | -3,60 | 0,0041 | 0,0402 |
| ***LINC02851*** | 7,23 | 13,48 | 10,35 | 1,71 | 0,00 | 0,86 | 0,08 | -3,64 | 0,0032 | 0,0341 |
| ***GPR63*** | 15,75 | 18,91 | 17,33 | 0,00 | 2,83 | 1,41 | 0,08 | -3,63 | 0,0000 | 0,0009 |
| ***PIP4P2*** | 19,83 | 14,89 | 17,36 | 1,43 | 1,29 | 1,36 | 0,08 | -3,71 | 0,0000 | 0,0007 |
| ***ITGA11*** | 13,16 | 7,04 | 10,10 | 1,43 | 0,00 | 0,71 | 0,07 | -3,87 | 0,0026 | 0,0292 |
| ***LYSET*** | 9,45 | 10,87 | 10,16 | 1,43 | 0,00 | 0,71 | 0,07 | -3,87 | 0,0025 | 0,0285 |
| ***MX1*** | 433,23 | 381,10 | 407,16 | 24,85 | 25,45 | 25,15 | 0,06 | -4,08 | 0,0000 | 0,0000 |
| ***SEMA3A*** | 10,93 | 8,65 | 9,79 | 1,14 | 0,00 | 0,57 | 0,06 | -4,13 | 0,0022 | 0,0256 |
| ***NEDD4*** | 8,71 | 21,73 | 15,22 | 1,71 | 0,00 | 0,86 | 0,06 | -4,20 | 0,0000 | 0,0009 |
| ***FRY*** | 26,68 | 18,91 | 22,80 | 1,71 | 0,77 | 1,24 | 0,05 | -4,24 | 0,0000 | 0,0000 |
| ***PRKCE*** | 9,45 | 13,08 | 11,26 | 1,14 | 0,00 | 0,57 | 0,05 | -4,33 | 0,0006 | 0,0087 |
| ***IFI44L*** | 482,89 | 519,74 | 501,31 | 23,99 | 26,73 | 25,36 | 0,05 | -4,36 | 0,0000 | 0,0000 |
| ***BCL7A*** | 16,12 | 7,65 | 11,88 | 0,00 | 1,03 | 0,51 | 0,04 | -4,48 | 0,0003 | 0,0046 |
| ***SREBF1*** | 19,27 | 9,66 | 14,46 | 0,86 | 0,26 | 0,56 | 0,04 | -4,71 | 0,0000 | 0,0005 |
| ***RAB31*** | 7,78 | 7,45 | 7,61 | 0,57 | 0,00 | 0,29 | 0,04 | -4,71 | 0,0053 | 0,0493 |
| ***OSM*** | 84,68 | 92,16 | 88,42 | 3,14 | 2,83 | 2,98 | 0,03 | -4,94 | 0,0000 | 0,0000 |
| ***IFIT1*** | 29,46 | 24,55 | 27,01 | 1,43 | 0,00 | 0,71 | 0,03 | -5,28 | 0,0000 | 0,0000 |
| ***CEND1*** | 8,52 | 8,05 | 8,29 | 0,29 | 0,00 | 0,14 | 0,02 | -5,71 | 0,0013 | 0,0173 |
| ***CCHCR1*** | 11,30 | 7,04 | 9,17 | 0,00 | 0,00 | 0,00 | 0,00 | -8,38 | 0,0001 | 0,0021 |
| ***SIX1*** | 6,86 | 7,65 | 7,25 | 0,00 | 0,00 | 0,00 | 0,00 | -8,04 | 0,0011 | 0,0147 |
| ***ANK2*** | 7,78 | 7,45 | 7,61 | 0,00 | 0,00 | 0,00 | 0,00 | -8,11 | 0,0007 | 0,0103 |
| ***DNAJC6*** | 19,09 | 7,45 | 13,27 | 0,00 | 0,00 | 0,00 | 0,00 | -8,91 | 0,0000 | 0,0000 |
| ***PLA2G4A*** | 14,45 | 8,25 | 11,35 | 0,00 | 0,00 | 0,00 | 0,00 | -8,69 | 0,0000 | 0,0002 |
| ***KIAA1217*** | 9,64 | 14,09 | 11,86 | 0,00 | 0,00 | 0,00 | 0,00 | -8,75 | 0,0000 | 0,0001 |
| ***CASQ1*** | 9,64 | 10,06 | 9,85 | 0,00 | 0,00 | 0,00 | 0,00 | -8,48 | 0,0000 | 0,0010 |
| ***C19orf38*** | 9,45 | 8,25 | 8,85 | 0,00 | 0,00 | 0,00 | 0,00 | -8,33 | 0,0002 | 0,0030 |
| ***GUCY1B1*** | 9,64 | 10,06 | 9,85 | 0,00 | 0,00 | 0,00 | 0,00 | -8,48 | 0,0000 | 0,0010 |

**Supplementary Table 7. Sequence of the repair templates used in this study**

| Template ID | Sequence (5'--> 3')[a, b] |
|---|---|
| miR-92b | ttactgtatcaaatgatatccaactctaagagaacttctagtatgaagatcaatgttgaaagtaaaatatttagattattaataacaaattatttaa<br>gggagaattagtttgtttagagctttttttcttaactatatttggaattaaaatgtcttaattagatatttaataaaattaacatttagaattaaagta<br>tttttctttaaaagtatatttgcattttgctagcttaagtttattgaaactgagctcttgtatgtcatttcatgtttttcccttgatgtgaagtcttaccta<br>tttgatgaagatgtcttttgaaaggtgtactgcaaggaacaaaatgtttgtaaattctccttttaccaaggtaaagatcaaattttataaatttactt<br>gtttgtttatacaaggaaaaataacttcatatattgaatatattcaaaagtttaagcatttagttgtattgccctgttaagttggcatagcaaataaa<br>tgcttttcttttcctcatttattctttgtgtttcctaacctatagcactgtgctgggcacagaatggacttcagttaagtttttgatgtagaaatgtttt<br>attattctacttaaaatctccttaaaaataattatgcatattacatcaatgttataatgtttaaacatagattttttacatgcattcttttttcctgaa<br>agaaaatatttttatattctttaggcgcgaatgtgtgtttaaaaaaaataaaa**cgggccccgggcgggcgggagggacgggacgcggtgcagt**<br>**gttgtttttccccgccaatattgcactcgtcccggcctccggccccccggccc** catactctacagttgtgttttaatgtatattaatgttactaat<br>gtgttttcagttttattgatagtcttttcagtattattgataatcttgttatttttagtatgattctgtaaaaatgaattaatactaatttttcagatgtat<br>catctcttaaaatactgtaattgcaatttaataattgtattgaatgccatcaagttttttaaaaagcttatgcagcattagaggaatttattttaatg<br>cacatttatattcaacatagacattaattcagatttttacttgggataaaacaaattctagttttccctttgttttgaaattacttttaaaatatgtcttt<br>acagataaatataaaatatattaagcattttgaacagagcttagaagacaatatttagtactgtttctgaatatttctttatatctgaaggggaaaa<br>gccatcaaaatatgtgaattaaatacctaaaattctggttgtcaaaacgtcacacttaaccataactttaaaggtagaaaaacccttacaagtga<br>ccaacccactcattggatagctaaggaaacacaaaactcaaataatttgctacacatcacagattggggtcttagtcttatgtcttctaactttata<br>atggtatactatttttttcatgtttttattaaaatgttttcttaaatccttagccaaagatttttgtttgataagtcaaaaataaggtcttgtttactttgt<br>ttactttttgtat |
| miR-17~92 | ttactgtatcaaatgatatccaactctaagagaacttctagtatgaagatcaatgttgaaagtaaaatatttagattattaataacaaattatttaa<br>gggagaattagtttgtttagagctttttttcttaactatatttggaattaaaatgtcttaattagatatttaataaaattaacatttagaattaaagta<br>tttttctttaaaagtatatttgcattttgctagcttaagtttattgaaactgagctcttgtatgtcatttcatgtttttcccttgatgtgaagtcttaccta<br>tttgatgaagatgtcttttgaaaggtgtactgcaaggaacaaaatgtttgtaaattctccttttaccaaggtaaagatcaaattttataaatttactt<br>gtttgtttatacaaggaaaaataacttcatatattgaatatattcaaaagtttaagcatttagttgtattgccctgttaagttggcatagcaaataaa<br>tgcttttcttttcctcatttattctttgtgtttcctaacctatagcactgtgctgggcacagaatggacttcagttaagtttttgatgtagaaatgtttt<br>attattctacttaaaatctccttaaaaataattatgcatattacatcaatgttataatgtttaaacatagattttttacatgcattcttttttcctgaa<br>agaaaatatttttatattctttaggcgcgaatgtgtgtttaaaaaaaataaaa**gtttgaggtgttaattctaattatctatttcaaatttagcagg**<br>**aaaaaagagaacatcaccttgtaaaactgaagattgtgaccagtcagaataatgtcaaagtgcttacagtgcaggtagtgatatgtgcatcta**<br>**ctgcagtgaaggcacttgtagcattatggtgacagctgcctcgggaagccaagttgggctttaaagtgcagggcctgctgatgttgagtgcttt**<br>**ttgttctaaggtgcatctagtgcagatagtgaagtagattagcatctactgccctaagtgctccttctggcataagaagttatgtattcatccaat**<br>**aattcaagccaagcaagtatataggtgttttaatagtttttgtttgcagtcctctgttagttttgcatagttgcactacaagaagaatgtagttgt**<br>**gcaaatctatgcaaaactgatggtggcctgctatttccttcaaatgaatgatttttactaattttgtgtacttttattgtgtcgatgtagaatctgc**<br>**ctggtctatctgatgtgacagcttctgtagcactaaagtgcttatagtgcaggtagtgtttagttatctactgcattatgagcacttaaagtactg**<br>**ctagctgtagaactccagcttcggcctgtcgcccaatcaaactgtcctgttactgaacactgttctatggttagttttgcaggtttgcatccagct**<br>**gtgtgatattctgctgtgcaaatccatgcaaaactgactgtggtagtgaaaagtctgtagaaaagtaagggaaactcaaacccctttctacac**<br>**aggttgggatcggttgcaatgctgtgtttctgtatggtattgcacttgtcccggcctgttgagtttggtggggattgtgaccagaagattttgaaa**<br>**attaaatattactgaagatttcgacttccactgttaaatgtacaagatacatgaa**catactctacagttgtgttttaatgtatattaatgttactaa<br>tgtgttttcagttttattgatagtcttttcagtattattgataatcttgttatttttagtatgattctgtaaaaatgaattaatactaatttttcagatgta<br>tcatctcttaaaatactgtaattgcaatttaataattgtattgaatgccatcaagttttttaaaaagcttatgcagcattagaggaatttattttaat<br>gcacatttatattcaacatagacattaattcagatttttacttgggataaaacaaattctagttttccctttgttttgaaattacttttaaaatatgtct<br>ttacagataaatataaaatatattaagcattttgaacagagcttagaagacaatatttagtactgtttctgaatatttctttatatctgaaggggaa<br>aagccatcaaaatatgtgaattaaatacctaaaattctggttgtcaaaacgtcacacttaaccataactttaaaggtagaaaaacccttacaagt<br>gaccaacccactcattggatagctaaggaaacacaaaactcaaataatttgctacacatcacagattggggtcttagtcttatgtcttctaacttta<br>taatggtatactatttttttcatgtttttattaaaatgttttcttaaatccttagccaaagatttttgtttgataagtcaaaaataaggtcttgtttacttt<br>gtttactttttgtat |

[a] **Bold** indicates the integrated sequence

**Supplementary Table 8. Landmark sequences used for the long read amplicon sequencing analysis**

| Landmark ID | Sequence (5'--> 3') |
|---|---|
| full_insertion | GTTTGAGGTGTTAATTCTAATTATCTATTTCAAATTTAGCAGGAAAAAAGAGAACATCACCTTGTAAAACTGAAGATTGTGACCAGTCAGAATAATGTCAAAGTGCTTACAGTGCAGGTAGTGATATGTGCATCTACTGCAGTGAAGGCACTTGTAGCATTATGGTGACAGCTGCCTCGGGAAGCCAAGTTGGGCTTTAAAGTGCAGGGCCTGCTGATGTTGAGTGCTTTTTGTTCTAAGGTGCATCTAGTGCAGATAGTGAAGTAGATTAGCATCTACTGCCCTAAGTGCTCCTTCTGGCATAAGAAGTTATGTATTCATCCAATAATTCAAGCCAAGCAAGTATATAGGTGTTTTAATAGTTTTTGTTTGCAGTCCTCTGTTAGTTTTGCATAGTTGCACTACAAGAAGAATGTAGTTGTGCAAATCTATGCAAAACTGATGGTGGCCTGCTATTTCCTTCAAATGAATGATTTTTACTAATTTTGTGTACTTTTATTGTGTCGATGTAGAATCTGCCTGGTCTATCTGATGTGACAGCTTCTGTAGCACTAAAGTGCTTATAGTGCAGGTAGTGTTTAGTTATCTACTGCATTATGAGCACTTAAAGTACTGCTAGCTGTAGAACTCCAGCTTCGGCCTGTCGCCCAATCAAACTGTCCTGTTACTGAACACTGTTCTATGGTTAGTTTTGCAGGTTTGCATCCAGCTGTGTGATATTCTGCTGTGCAAATCCATGCAAAACTGACTGTGGTAGTGAAAAGTCTGTAGAAAAGTAAGGGAAACTCAAACCCCTTTCTACACAGGTTGGGATCGGTTGCAATGCTGTGTTTCTGTATGGTATTGCACTTGTCCCGGCCTGTTGAGTTTGGTGGGGATTGTGACCAGAAGATTTTGAAAATTAAATATTACTGAAGATTTCGACTTCCACTGTTAAATGTACAAGATACATGAA |
| full_WT | CCTTGGAGTAAAGTAGCAGCACATAATGGTTTGTGGATTTTGAAAAGGTGCAGGCCATATTGTGCTGCCTCAAAAATACAAGGATCTGATCTTCTGAAGAAAATATATTTCTTTTTATTCATAGCTCTTATGATAGCAATGTCAGCAGTGCCTTAGCAGCACGTAAATATTGGCGTTAAGATTCTAAAATTATCTCCAGTATTAACTGTGCTGCTGAAGTAAGGTTGAC |
| gRNA1 | TGTGCTGCTACTTTACTCCA |
| gRNA2 | TTAACTGTGCTGCTGAAGTA |
| pcr_fwd | GTATGTGATGAGGGGGACCCTG |
| pcr_rev | AGAGACAGGGTTTCACCACATTG |
